# Mutually Antagonistic Protein Pairs of Cancer

**DOI:** 10.1101/2021.08.30.458281

**Authors:** Ertuğrul Dalgıç

**Author notes:** Address correspondence to: Ertuğrul Dalgıç, Zonguldak Bulent Ecevit Universitesi Tip Fakultesi Dekanligi, Tibbi Biyoloji Anabilim Dali, Kozlu, Zonguldak, 67600, Turkey.

## Abstract

Switch-like behavior of tumorigenesis could be governed by antagonistic gene and protein pairs with mutual inhibition. Unlike extensive analysis of gene expression, search for protein level antagonistic pairs has been limited. Here, potential cancer type specific antagonist protein pairs with mutual inhibition were obtained from large scale datasets. Cancer samples or cancer types were compared to retrieve potential protein pairs with contrasting differential expression patterns. Analysis of two different protein expression datasets showed that a few proteins participate in most of the mutually antagonistic relationships. Some proteins with highly antagonistic profile were identified, which could not be attained from a differential expression or a correlation based analysis. The antagonistic protein pairs are sparsely connected by molecular interactions. Glioma, melanoma, and cervical cancer, are more frequently associated with antagonistic proteins than most of the other cancer types. Integrative analysis of mutually antagonist protein pairs contributes to our understanding of systems level changes of cancer.

## Introduction

Cancer could be viewed as the result of switch like behavior of cells (Mills et al. 2010, Balentine et al. 2011, Torres et al. 2018). Robustness is preserved, as cells transform from normal state to cancer state. The switch like behavior could be best understood by a systems level view of molecular networks. Some specialized molecular interaction patterns, or circuits, could play significant roles for the switch (Siegal-Gaskins et al. 2011). Previously gene circuits which play a role in the bistability of normal network transition to cancer network, were investigated (Shiraishi et al. 2010). Interaction network neighbor genes that show contrasting expression levels for cancer were investigated and shown as candidate players for circuits underlying the switch like behavior.

Antagonist protein pairs with mutual inhibition have critical roles for generating bistability (Gardner et al. 2000, Yao et al. 2008, Parashar et al. 2009, Shin et al. 2017, Matsuoka et al. 2018, Nguyen et al. 2018, Rata et al. 2018]. Two proteins of such antagonist pairs, negatively regulate each other directly or indirectly. Mutually antagonist proteins show contrasting expression and activity levels in two different stable states. Inhibitory effects of Retinoblastoma (RB) and E2F Transcription Factor on each other, is important for the bistability during a critical cell cycle decision (Yao et al. 2008). A mutual inhibitory mechanism was shown for Phosphatidylinositol 3,4,5-trisphosphate (PIP3) and PIP3 phosphatase, providing polarity such that one part of a cell would have PIP3, and another part would have PIP3 phosphatase (Matsuoka et al. 2018). The interaction between PIP3 and its phosphatase provide the bistable behavior of directional movement. Another mutual inhibitory mechanism was shown for RhoA and Rac1, providing bistability for cell migration (Nguyen et al. 2018). Under cellular stress conditions, XIAP-associated factor 1 (XAF1) and metallothionein 2A (MT2A) induce the degradation of each other and this mutual regulation is important for the switch like behavior for tumorigenesis (Shin et al. 2017). Such antagonistic mechanisms were confirmed in expression levels as XAF1 and MT2A were negatively coexpressed in several primary tumors and cell lines (Shin et al. 2017). For the tumors where XAF1 was low and MT2A was high, induction of XAF1 lowered MT2A levels and caused a regression in tumorigenesis. Cyclin dependent kinase 1 (CDK1) and protein phosphatase 2A (PP2A) indirectly inhibit each other and this mutual inhibitory mechanism is critical for mitosis entry switch (Rata et al. 2018). When the mutual inhibitory effects of CDK1 and PP2A were disrupted, the switch like entry into mitosis was disrupted. Such examples suggest mutually antagonistic relationships between proteins as important mechanisms for several cellular events including tumorigenesis.

Previous studies for the estimation of antagonistic pairs focused on gene expression datasets rather than protein datasets (Shiraishi et al. 2010). However, there is no strong correlation between mRNA and protein expression levels for the majority of human genes (Kosti et al. 2016). Protein level changes, which are not reflected in genomic level could play role in cancer mechanisms (Jarnuczak et al. 2021). Therefore, protein level analysis is required for understanding cancer related antagonistic relationships.

Cancer is a complex heterogenic disease as each cancer sample has its own unique gene and protein expression patterns (Axelsen et al. 2007, Kosti et al. 2016, Schneider et al. 2017). There are different tissue specific oncogenic mechanisms (Schneider et al. 2017). Proteins could depict contrasting expression and activity levels in different cancer samples and cancer types. Therefore, two proteins underlying a bistable switch could show opposite behavior in two different cancer samples or cancer types. Here this is exploited to analyze antagonistic pairs across various cancer samples (CS) and cancer types (CT). Protein pairs with antagonistic behavior are such that when the expression level of a protein is up, the expression of the other protein is down and vice versa. Here mutually antagonistic protein pairs (MAPP) are identified by selecting pairs of proteins which are ON (upregulated) and OFF (downregulated) in at least one CS or CT, and OFF and ON in at least one other CS or CT. Some proteins were found to have high number of antagonistic links with other proteins. A few protein pairs were linked by molecular interactions. Members of MAPP could potentially be key players for the switch like behavior of cancer.

## Results

### Uncovering MAPP

MAPP were explored by selecting protein pairs with opposing differential expression profiles; which are ON and OFF respectively in at least one CS or CT, and OFF and ON in at least one other CS or CT (Figure 1A-B). 342 distinct MAPP among 80 proteins were identified in The Cancer Proteome Atlas (TCPA) dataset based on comparison of the CS (Supplementary Table 1). On the other hand, 341 distinct protein pairs were found by comparing the CT of the TCPA dataset (Supplementary Table 1). All of the CT based MAPP were the same with the CS based pairs except for the missing HSP70-PKCPANBETAII_pS660 pair. Significance of the MAPP was analyzed by randomization tests. 187 CS based pairs and 111 CT based pairs are significant (FDR corrected p-value < 0.05) (Supplementary Table 1). 97 pairs are significant in both CS and CT based MAPP. Network view of the combined significant MAPP of CS and CT based analyses shows that a few proteins such as EPPK1, MYH11, ESR1, CAV1, ANXA1, PEA15, TFRC, TUBA1B, GAPDH, and SRC are highly connected whereas most of the proteins are lowly connected in the combined list of MAPP which include both CS and CT based pairs (Figure 2A, Supplementary Figure 1). Degree distribution of the MAPP networks are similar for individual CS or CT based analyses (Supplementary Figure 2).

**Figure 1.**
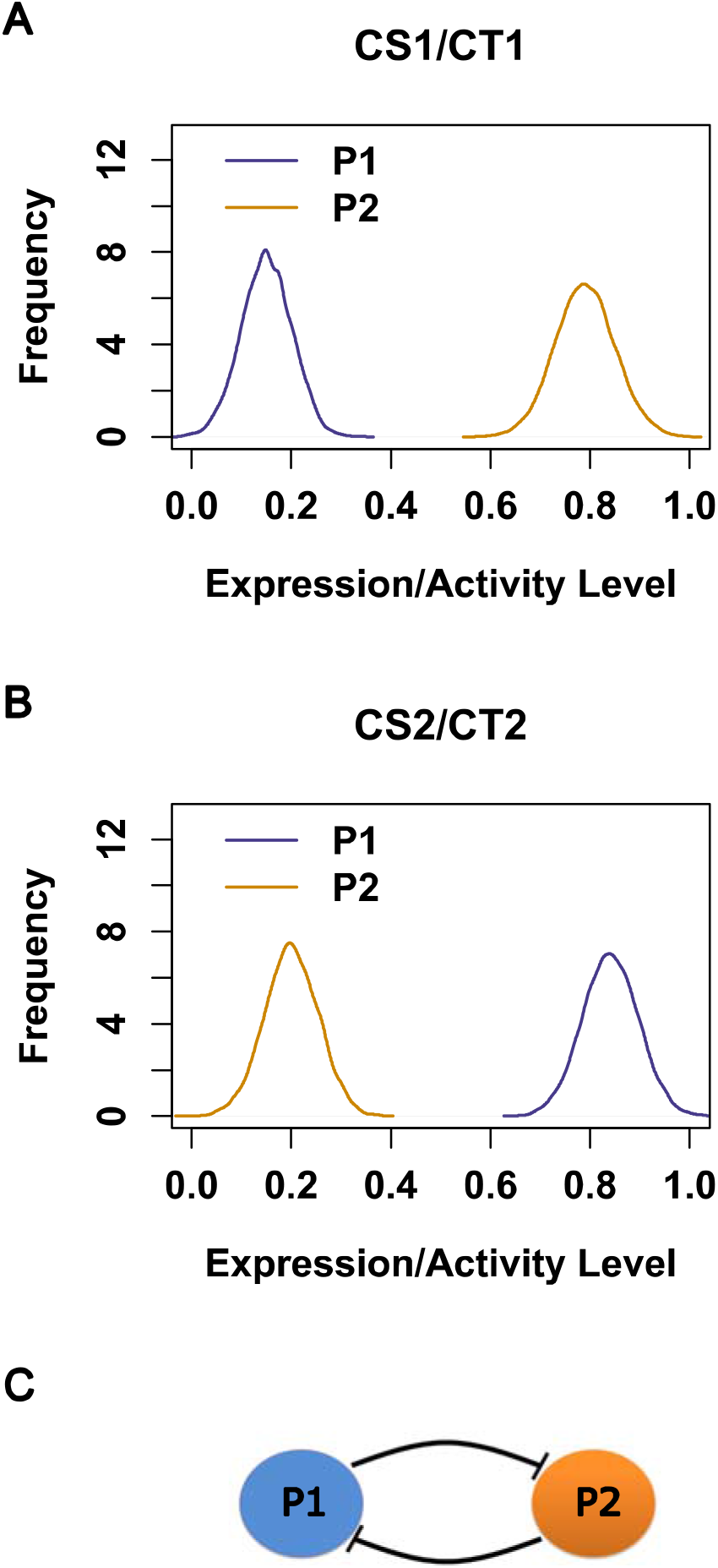
Hypothetical MAPP expression and interaction motif. **(A)** Expression or activity levels of two proteins (P1 and P2) which have a mutually antagonistic relationship with each other across two different cancer samples (CS1 and CS2) or cancer types (CT1 and CT2). **(B)** Potential direct or indirect interactions between the proteins in a MAPP. One explanation for the opposing expression or activity levels of P1 and P2, is the negative effect of each protein to the other one in the pair.

**Figure 2.**
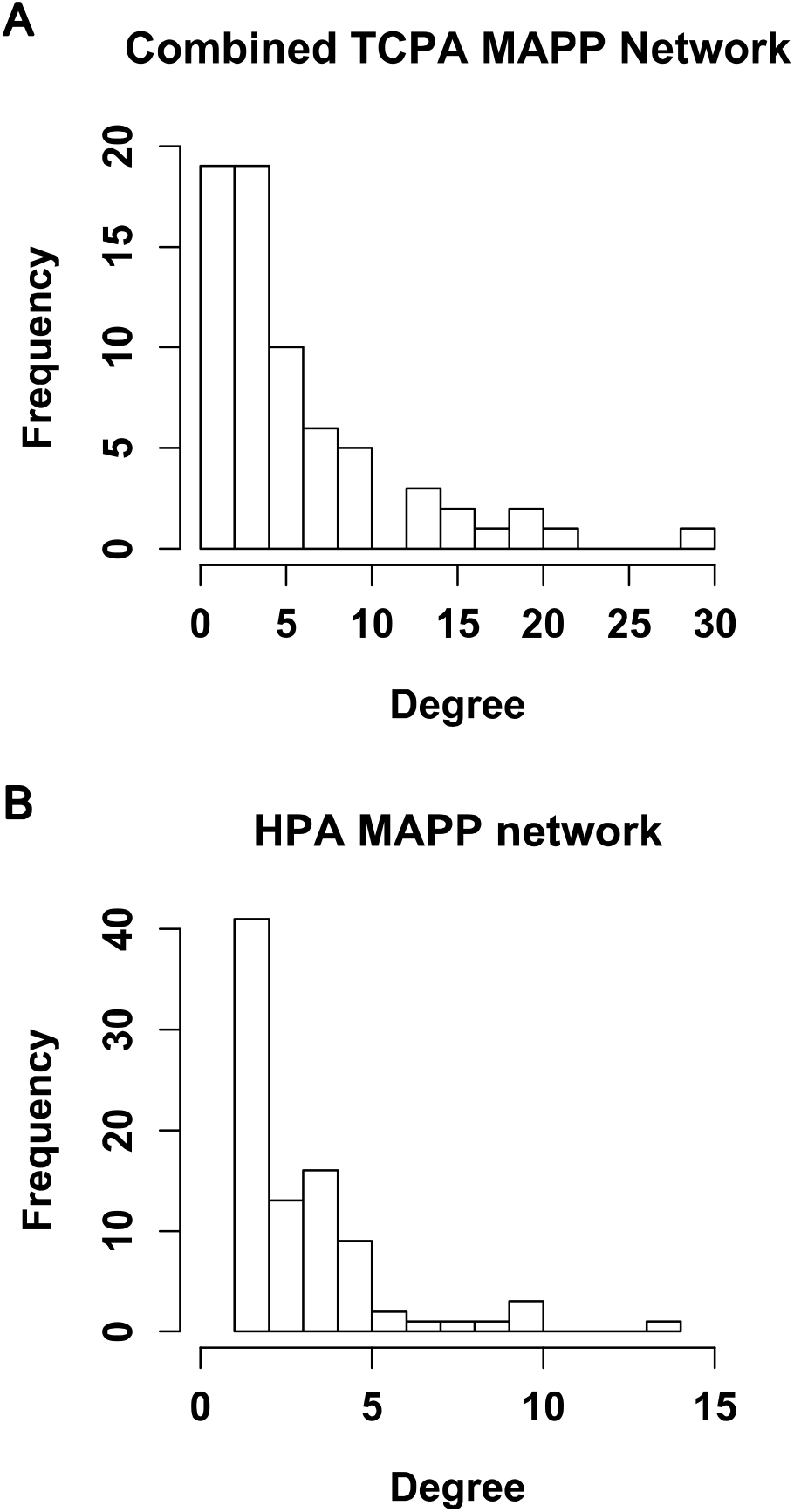
Degree distribution of MAPP networks. **(A)** CS and CT based MAPP networks of the TCPA dataset were combined. Degree values (number of links) of the unique MAPP of the combined list were used for the distribution. **(B)** MAPP network of the HPA dataset were used for the distribution.

Human Protein Atlas (HPA) dataset provides only cancer type based values unlike TCPA. However, it has a higher coverage of human proteome. There are 19651 unique proteins in the HPA dataset and 217 unique proteins in the TCPA dataset based on NCBI gene symbols. All of the 217 TCPA proteins are also present in the HPA dataset. MAPP were defined based on the comparison of the HPA cancer types. 148 distinct MAPP among 90 proteins were identified (Supplementary Table 2). 142 pairs are significant based on the randomization test (FDR corrected p-value < .0.05). Remarkably, there is no common protein pair between the TCPA and the HPA MAPP lists, and GAPDH is the only common protein. HPA MAPP network has a few highly connected proteins such as LRRC26, ABHD3, SPIDR, PKP3, PC, and KRT5 while most of the proteins are lowly connected (Figure 2B, Supplementary Figure 3). Thus, analyses of both datasets showed that a few proteins participate in most of the mutually antagonistic relationships.

### Comparison of Antagonistic Relationships to Differential Expression and Correlation

Next, assessment of cancer proteins for mutual antagonism was compared to the typical analyses of differential expression and coexpression. Parameters for MAPP, differential expression and expression correlation (PCPP; positive correlation based protein pairs, NCPP; negative correlation based protein pairs) were defined for each protein in the TCPA and HPA datasets (see Methods). MAPP, PCPP, and NCPP based differential ratio values were calculated as the difference of their ratio values from the differential expression based ratio value for each protein. The profile of mutual antagonism based values differs from positive and negative correlation for the CS and CT based analyses of the TCPA and HPA datasets (Figure 3, Supplementary Tables 3-5). For the CT based analyses of both datasets, MAPP based values of some proteins are considerably different from differential expression unlike correlation based values. Unlike PCPP and NCPP, MAPP based differential ratio values are highly positive for some proteins. Therefore, the current analysis provides molecules with distinct antagonistic characteristics which could not be obtained from a differential expression or a correlation based analysis.

**Figure 3.**
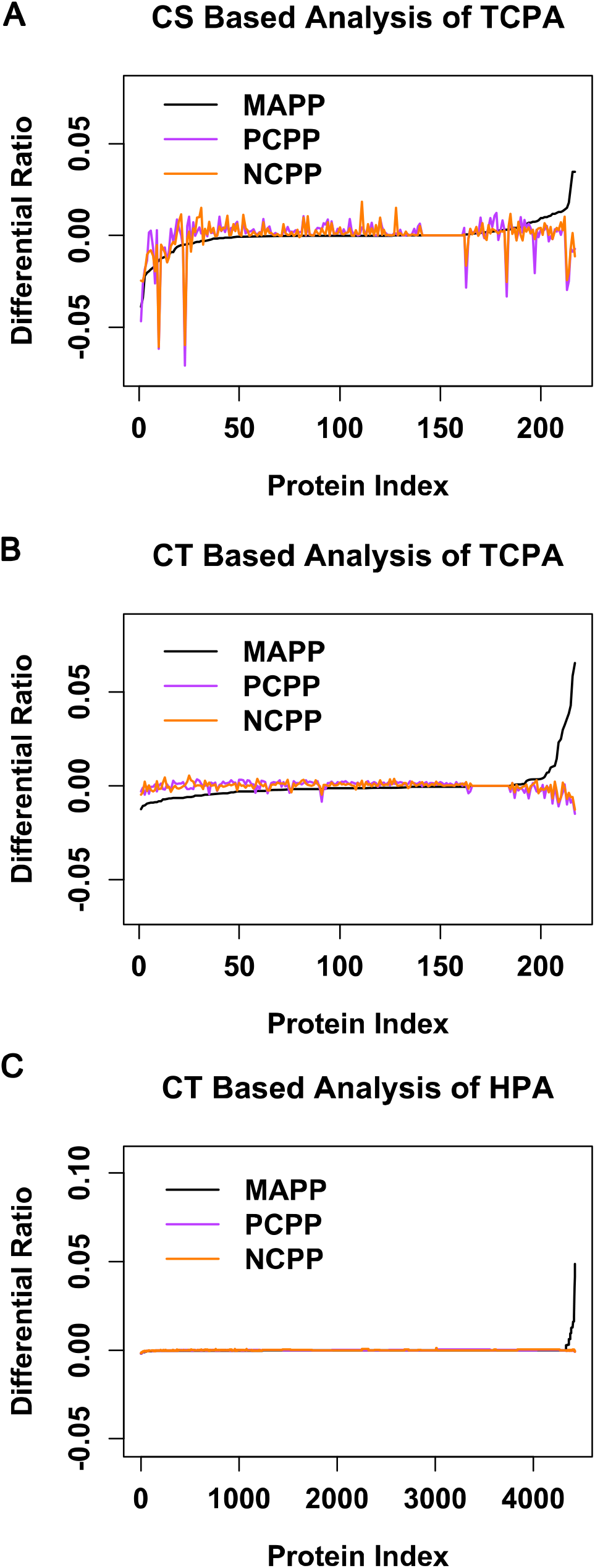
Differential ratio analysis of differentially expressed proteins in TCPA and HPA datasets. Difference of the weighted ratio of proteins based on their mutually antagonistic protein pairs (MAPP), positive correlation protein pairs (PCPP) and negative correlation protein pairs (NCPP) from the weighted ratio value for differential expression. Protein index is sorted from the lowest to highest MAPP differential ratio, **(A)** Cancer sample (CS) based analysis of TCPA dataset, **(B)** Cancer type (CT) based analysis of TCPA dataset, **(C)** Cancer type (CT) based analysis of HPA dataset.

TCPA proteins such as EPPK1, MYH11, TUBA1B, ANXA1, TFRC, PEA15, ESR1, PEA15, CAV1, GAPDH, and CCNB1 have highly positive MAPP based differential ratio values in the CS or CT based analyses, thus are highly antagonistic with other proteins (Table 1). Some proteins such as TUBA1B and ANXA1 are common to the CS and CT based lists, others such as EPPK1 and MYH11 are specific to only one of the lists. Although EPPK1 and MYH11 have the highest differential expression values in the CS based analysis, they have positive MAPP based differential ratio values only in the CT based analysis (Supplementary Tables 3-4). On the other hand, GSK3ALPHABETA_pS21S9 (GSK3A, GSK3B) and ATM have relatively lower differential expression values but positive MAPP based ratio values. Although, a similar profile was observed for both CS and CT based analyses of differential ratio values, some proteins have disparate values. Also most of the top proteins with the highest positive and negative correlation numbers do not have highly positive MAPP based differential ratio values (Supplementary tables 3-4).

**Table 1.** Top ranked proteins for the MAPP based differential ratio values of the TCPA dataset. Top 10 proteins in the combined list of CS and CT based analysis were listed. In addition to MAPP, PCPP and NCPP based values were also shown.

| TCPA protein symbol | NCBI<br>symbol | Differential<br>ratio for<br>PCPP | Differential<br>ratio for<br>NCPP | Differential<br>ratio for<br>MAPP | Type of<br>analysis<br>(CS or<br>CT) |
| --- | --- | --- | --- | --- | --- |
| EPPK1 | EPPK1 | -0.0149 | -0.01257 | 0.065482 | CT |
| MYH11 | MYH11 | -0.00722 | -0.00584 | 0.05841 | CT |
| ACETYLATUBULINLYS40 | TUBA1B | -0.00888 | -0.00596 | 0.042741 | CT |
| ANNEXIN1 | ANXA1 | -0.00421 | -0.00371 | 0.038592 | CT |
| TFRC | TFRC | -0.00993 | -0.00689 | 0.035851 | CT |
| ACETYLATUBULINLYS40 | TUBA1B | -0.00732 | -0.0114 | 0.03476 | CS |
| ANNEXIN1 | ANXA1 | -0.00894 | 0.001294 | 0.03458 | CS |
| PEA15_pS116 | PEA15 | 0.000745 | -0.00105 | 0.032279 | CT |
| ERALPHA | ESR1 | -0.00337 | 0.001688 | 0.029912 | CT |
| PEA15_pS116 | PEA15 | -0.00695 | -0.00707 | 0.026293 | CS |
| CAVEOLIN1 | CAV1 | -0.00609 | -0.00618 | 0.024879 | CT |
| GAPDH | GAPDH | -0.01095 | -0.00856 | 0.0231 | CT |
| CAVEOLIN1 | CAV1 | -0.02496 | -0.01968 | 0.017898 | CS |
| ERALPHA | ESR1 | -0.02972 | -0.02444 | 0.015211 | CS |
| CYCLINB1 | CCNB1 | -0.00586 | -0.00269 | 0.014624 | CT |

HPA proteins such as LRRC26, PC, ABHD3, PKP3, SPIDR, KRT5, KRT17, TRPS1, RNASEH2C, and GIGYF1 have highly positive MAPP based differential ratio values (Table 2, Supplementary Table 5). None of the top proteins with the highest differential expression number, positive and negative correlation numbers have positive MAPP based differential ratio values. Analyses of both TCPA and HPA datasets showed that mutual antagonism analysis provides unique proteins. Differential expression or correlation based values do not estimate the mutual antagonistic parameters of a protein.

**Table 2.**
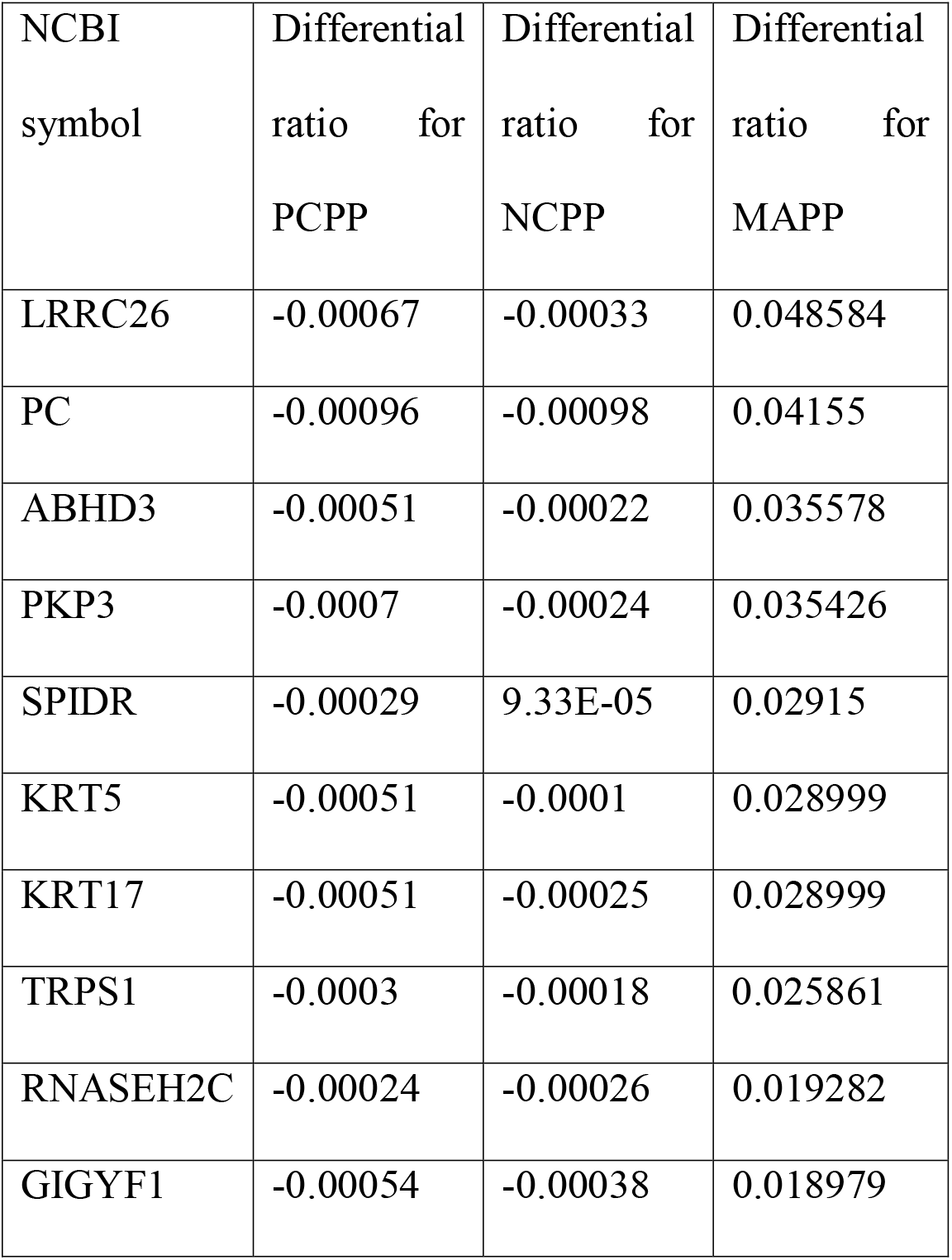
Top 10 ranked proteins for the MAPP based differential ratio values of the HPA dataset. In addition to MAPP, PCPP and NCPP based values were also shown.

| NCBI<br>symbol | Differential<br>ratio for<br>PCPP | Differential<br>ratio for<br>NCPP | Differential<br>ratio for<br>MAPP |
| --- | --- | --- | --- |
| LRRC26 | -0.00067 | -0.00033 | 0.048584 |
| PC | -0.00096 | -0.00098 | 0.04155 |
| ABHD3 | -0.00051 | -0.00022 | 0.035578 |
| PKP3 | -0.0007 | -0.00024 | 0.035426 |
| SPIDR | -0.00029 | 9.33E-05 | 0.02915 |
| KRT5 | -0.00051 | -0.0001 | 0.028999 |
| KRT17 | -0.00051 | -0.00025 | 0.028999 |
| TRPS1 | -0.0003 | -0.00018 | 0.025861 |
| RNASEH2C | -0.00024 | -0.00026 | 0.019282 |
| GIGYF1 | -0.00054 | -0.00038 | 0.018979 |

### Interaction Networks of MAPP

Protein-protein interactions (PPI) and protein-DNA interactions (PDI) between the antagonist proteins were investigated. Significant MAPP of the TCPA and HPA datasets were integrated with human PPI and PDI datasets (see Methods). Only 4 pairs from both CS and CT based MAPP of the TCPA dataset have direct protein-protein interactions; STAT5A-ESR1, CAV1-SRC, CTNNB1-YAP1, and MAP2K1-YAP1. CS based MAPP also has CAV1-SQSTM1 as a directly interacting pair which is not a significant CT based MAPP. Only one pair from both CS and CT based MAPP has a direct PDI; STAT5A-ESR1, where STAT5A is the TF of ESR1 gene. 2 other PDI are present in the CS based MAPP; AR-ANXA1 and ESR1-CTNNB1; where AR and ESR1 are the TF of ANXA1 and CTNNB1 genes respectively. STAT5A-ESR1 is the only pair which has both direct PPI and PDI. Furthermore, more TCPA MAPP have indirect PPI between them (Figure 4, Supplementary Table 6). Among the 210 significant TCPA MAPP (based on NCBI symbols), 34 CS and CT based pairs, 35 only CS based pairs, and 5 only CT based pairs have indirect interactions. ESR1, CAV1, SRC, YAP1, CTNNB1, GAPDH, and CCNB1 are highly connected in the PPI integrated network (Figure 4). On the other hand, none of the significant HPA MAPP have direct PPI or PDI between them. Furthermore, only one pair, which is HSPB1-PARD6A, has indirect PPI (Supplementary Table 7). Overall, there are very few antagonist pairs with direct molecular interactions. Some TCPA pairs have indirect interactions and HPA pairs have no direct or indirect interactions between them except for one.

**Figure 4.**
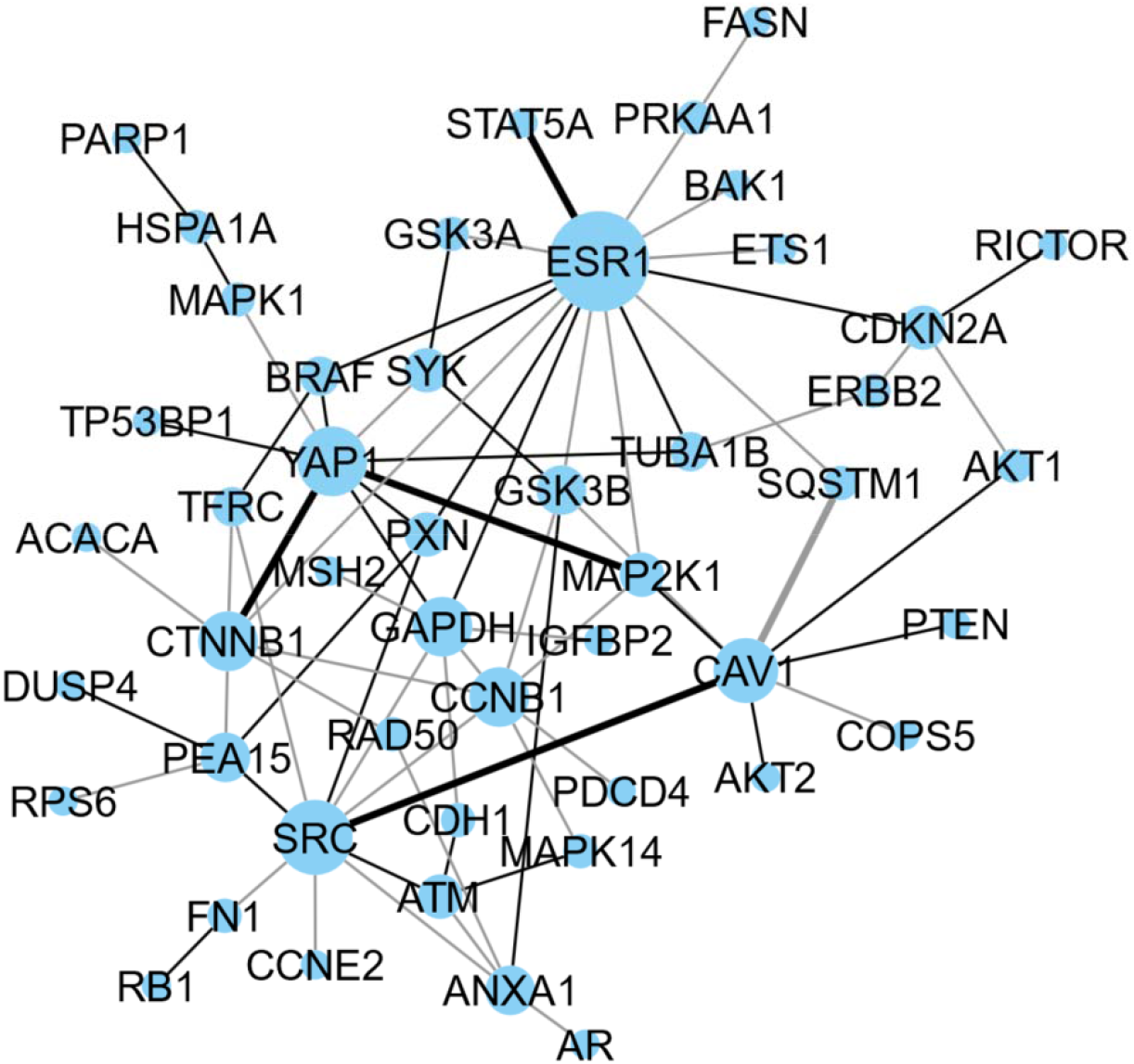
PPI interaction network of MAPP in the TCPA dataset. MAPP which are linked directly or indirectly in the human PPI network. Thickness of the edge represents direct PPI connection, color of the edge represents the presence of MAPP in one or both of the CS and CT based analysis. Size of the nodes are correlated to their degree values (number of links).

### Cancer Pairs of MAPP

MAPP definition of this study was based on the comparison of expression levels of two proteins in two different CS or CT (Figure 1). Finally CT pairs, which were used for the designation of significant MAPP of the TCPA and HPA datasets, were listed (Supplementary Tables 8-9). CT of significant MAPP of the TCPA and HPA datasets, include some cancer more frequently than most of the other cancers (Figure 5). LGG (Brain Lower Grade Glioma), SARC (Sarcoma), CESC (Cervical squamous cell carcinoma and endocervical adenocarcinoma), SKCM (Skin Cutaneous Melanoma) and BLCA (Bladder Urothelial Carcinoma) have higher counts of appearance in the CT lists of the TCPA dataset. Breast cancer, testis cancer, melanoma, glioma, cervical cancer, and endometrial cancer have high counts in the CT lists of the HPA dataset. None of these cancers were overrepresented in the datasets (Supplementary Table 10-11). Glioma (LGG), melanoma (SKCM), and cervical cancer (CESC) are frequently used by the antagonist proteins of both datasets.

**Figure 5.**
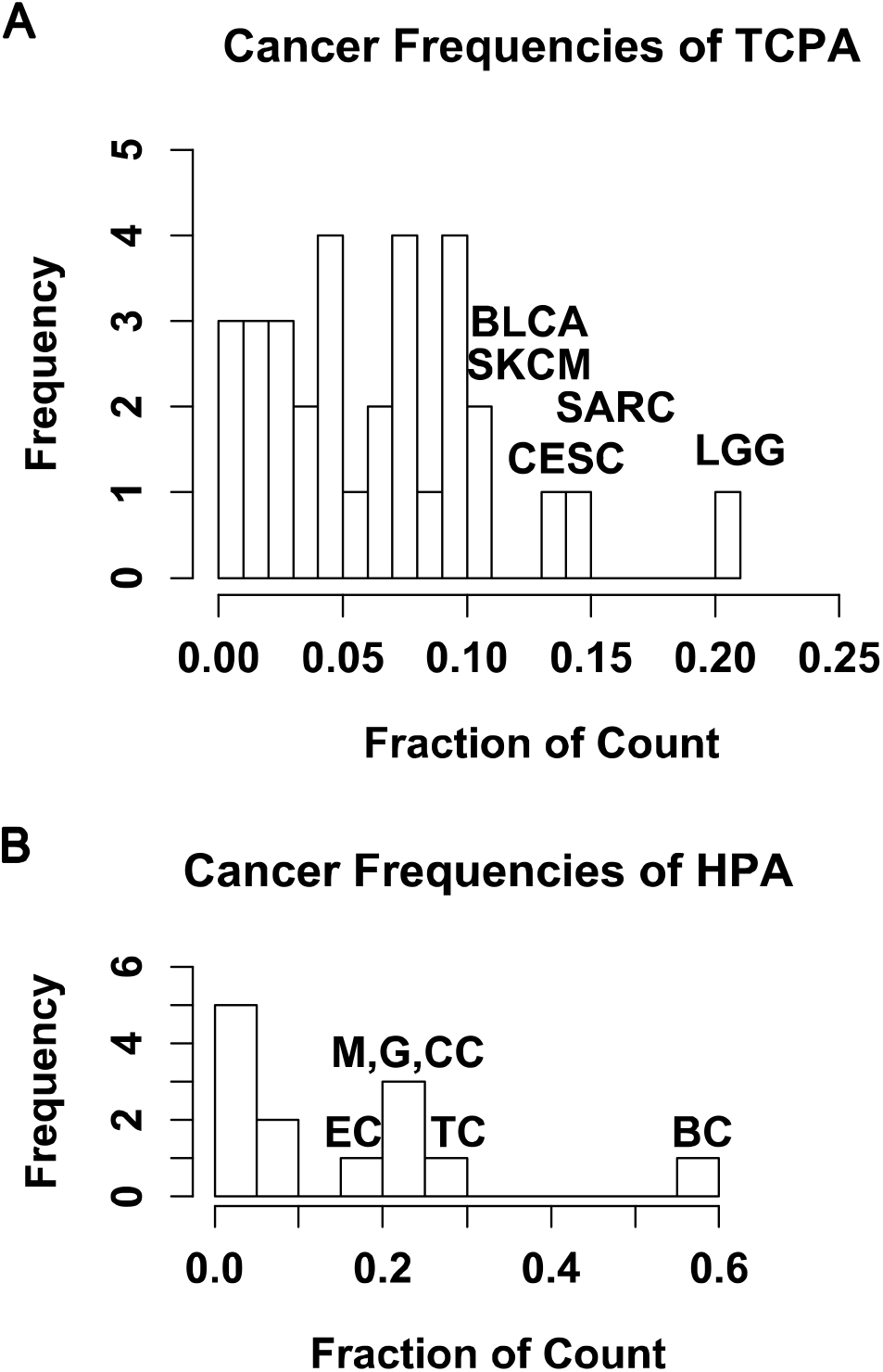
Frequencies of CT pairs associated with significant MAPP. **(A)** Frequency of CT counts for the TCPA dataset. **(B)** Frequency of counts for the HPA dataset. High ranking frequent cancers were shown. (BC: breast cancer, TC: testis cancer, CC: cervical cancer, G: glioma, M: melanoms, EC: endometrial cancer)

## Discussion

In this study, samples and types of cancer are compared among themselves to identify pairs of proteins with contrasting levels, unlike typical studies focusing on only normal and cancer comparison. For some proteins, MAPP associated ratio values, their associated MAPP count relative to all other proteins, appear to be distinctive. Such information for those proteins cannot be received from differential equation or correlation based analyses. Even, negative correlation values of such proteins are not distinguishing. Therefore, examination of mutual antagonism by comparing CS or CT to each other, provides unique proteins with distinct characteristics. MAPP analyses showed that a few proteins participate in most of the mutually antagonistic relationships, and these proteins have generally positive MAPP based ratio values. EPPK1, MYH11, TUBA1B, ANXA1, TFRC, PEA15, ESR1, PEA15, CAV1, GAPDH, CCNB1, LRRC26, ABHD3, SPIDR, PKP3, PC, and KRT5 are some of the proteins with a high number appearances in MAPP networks and distinctive MAPP based ratio values.

STAT5A and ESR1 (ER alpha) pair is in both CS and CT based MAPP of the TCPA dataset analysis and has both PPI and PDI. There is experimental evidence for potential antagonistic or redundant relationships between them (Wang and Cheng 2004, Miermont et al. 2010). Another MAPP of the TCPA is SRC-CAV1. They also have a direct interaction. The relationship between SRC and CAV1 was shown previously (Thomas et al. 2011). EPPK1-PEA15_pS116 pair has the highest number of occurrences in the TCPA dataset analysis. Therefore a potential antagonistic relationship between the two proteins could be investigated. Currently there is no experimental evidence for a direct relationship between these two proteins. PEA15_pS116 specifically represents the form of PEA15 (proliferation and apoptosis adaptor protein 15) which is phosphorylated at Ser-116, thus actively inhibiting apoptosis (Renganathan et al. 2005). EPPK1 (Epiplakin 1), knock-down is associated with cell migration and proliferation (Kokado et al. 2016). Therefore, EPPK1 and PEA15 might have mutually exclusive anti-proliferative effects in several cancer samples. Since the current study is based on the comparison of CS or CT to each other, identified pairs could have roles in bistability or they could be dispensable for each other. Thus, determination of their mechanistic roles for cancer needs further evidence.

For protein expression or activity levels, TCPA and HPA databases were used. HPA dataset has a high number of proteins, however, only the values for distinct cancer types could be analyzed. On the other hand, TCPA dataset has very few number of proteins but a high number of samples across various cancer types. There is no common pair between the significant MAPP of TCPA and HPA. This could be due to different sizes of the datasets. Expression scores of the two datasets are also different, contributing to the differences of the results. Significant MAPP network of HPA also does not have any direct PPI and has only one indirect PPI, which could be due to incomplete interaction databases. Besides, some pairs might be connected by more indirect connections.

HPA dataset values ignore modifications such as phosphorylation which could determine the activity of some proteins. On the other hand, TCPA dataset values have some modification-specific values. Therefore, the reason for the absence of some TCPA MAPP from HPA MAPP could be due to the negligence of such modification-specific values. When several normal and cancer tissues were examined, 64-80% of cases with significant expression differences, had low XAF1 and high MT2A levels confirming the role of antagonistic mechanism between these proteins in cancer (Shin et al. 2017). Therefore when using generic datasets like HPA, some antagonistic pairs will be missed. In order to capture a larger scale identification of such pairs, individual sample based datasets are needed. Furthermore, because of the heterogenic nature of tumor tissues, single cell based proteomics datasets might be needed for elucidating all antagonistic pairs. Noting the importance of chemical modifications for protein function, phosphorylation specific measurements provide a more biologically meaningful perspective. The pairs obtained by the analysis of HPA dataset ignores such modifications, therefore the high expression of a certain protein does not guarantee an increase in its activity. Therefore, such pairs should be further examines computationally and experimentally for changes in their activity, for if there is actually an antagonistic relationship between the two proteins with respect to their activities. With the availability of datasets of bigger size, in terms of the number of proteins and the number of samples, in the future, could enable us to find a higher number of significant protein pairs that could play a role in cancer.

Different bistability generating feedback circuits could work together to have a robust control on cellular events (Rata et al. 2018). Similarly different antagonistic pairs could also operate cooperatively to achieve important transitions such as tumorigenesis. The potential bistability generating antagonistic circuits identified here could work in different combinations for different cases. Frequently observed proteins could have important roles for different cancer types. Elucidating such mechanisms could be critical for systems level understanding of cancer.

## Methods

### Prediction of MAPP across Cancer Samples

Cancer sample expression values for different proteins were obtained from The Cancer Proteome Atlas (TCPA) database version 4.2 (https://tcpaportal.org) (Li et al. 2013). TCPA dataset values are based on Reverse phase protein arrays (RPPA). TCPA has 4 levels of data processing steps. Level 4 values have normalized values such that different batch effects were removed and values for different cancer samples were merged. Here, relative quantitative values of Pan-Cancer Level 4 TCPA dataset were used to assess differential expression of 217 proteins (including different modifications of the same protein, for instance P27_pT198, a phosphorylated form of P27, was treated as a separate protein, see Supplementary Information). The dataset has a total of 7694 cancer samples for 32 different cancer types. The threshold values -2 and 2 were chosen to define differentially expressed proteins in the TCPA dataset (see Supplementary Information). Proteins with an expression value higher than 2 were defined as ON and the ones with an expression value lower than -2 were defined as OFF for each cancer sample. Then, mutually antagonistic protein pairs (MAPP) were defined as those proteins which are ON-OFF in at least one sample, and OFF-ON in at least one other sample ignoring the cancer type of the sample.

Next, a permutation test was done by shuffling the expression values of each sample. Antagonist pairs were similarly found in the random datasets. The number of sample pairs, for which the antagonist relationship was defined for each protein pair, was compared to random values and p-values were defined as the fraction of random iterations with an equal or higher number of samples pairs. Multiple testing correction of the p-values was done by False Discovery Rate (FDR). 0.05 was chosen as the FDR correct p-value cutoff to define the significant MAPP.

### Prediction of MAPP across Cancer Types

TCPA dataset also has cancer type information for all samples. Therefore, a similar analysis was also done based on cancer types, in which differential expression of a protein in at least one of the samples belonging to a cancer type was considered when searching for protein pairs with contrasting ON-OFF trends. MAPP were defined as those proteins which are ON-OFF in at least cancer type, and OFF-ON in at least one other cancer type. Permutation test based p-values for protein pairs were defined similarly and multiple testing correction was done by FDR.

In addition to TCPA, Human Protein Atlas (HPA) database protein expression values were used to assess antagonistic relationships (http://www.proteinatlas.org) (Uhlen et al. 2015). Normal tissue and cancer expression (pathology) values for different proteins were obtained from HPA database version 20. Normal tissue and pathology datasets were used to determine semi-quantitative differential expression of proteins. Manual scoring of immunohistochemistry analysis is shown as ‘not detected’, ‘low’, ‘medium’, and ‘high’. Cancer dataset has scoring values for 15308 proteins and 17 different cancer types after removing rows with missing values and very low number of measurements (see Supplementary Information). There are multiple measurements for different cases of a cancer type and there is a variation among the different samples. However, there is a single scoring value for normal tissue cell type samples. To determine the differential expression of a protein, proteins with binary expression (present or absent) in cancer and normal were determined. For the pathology dataset values, the Cancer Expression parameter (CE) was defined.

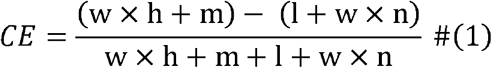

In equation 1; n, l, m, and h are the number of samples with ‘not detected’, ‘low’, ‘medium’, ‘high’ expression respectively, and w is the weight for ‘not detected’ and ‘high’ scores. As CE gets bigger or lower than 0, protein expression gets more definitive. Absolute CE threshold value was set to 0.5 and w was set to 2 (see Supplementary Information). Proteins with the CE value lower than -0.5 was defined as absent and the ones with the CE value higher than 0.5 was defined as present. Proteins with binary (present/absent) expression in all cancer types were specified by using the CE values. Secondly, differentially expressed proteins were determined by comparing the cancer expression to normal. The normal dataset has scoring values for 12100-13500 proteins, for different cell types from different tissues. Unlike the cancer dataset, the normal dataset has only a single value, therefore, proteins with ‘not detected’ score was defined as absent, and the ones with ‘high’ score was defined as present. Cancer types and tissues and cell types were matched manually (see Supplementary Information). For each cancer type-normal tissue match, proteins which are present in cancer and absent in normal tissue were defined as ON (upregulated), and proteins which are absent in cancer and present in normal tissue was defined as OFF (downregulated). MAPP were defined as those proteins which are ON-OFF in at least one cancer type, and OFF-ON in at least one other cancer type.

Monte Carlo simulation was done to observe the random distribution of CE. Random values were assigned for each protein as the distribution of the total values for each protein (h+m+l+n) were kept constant. For each protein pair, the p-value was defined as the number of random iterations, for which the number antagonist relationships across all cancer type pairs was not higher random. Multiple testing correction of the p-values was done by False Discovery Rate (FDR). 0.05 was chosen as the FDR correct p-value cutoff to define the significant MAPP.

### Differential Ratio Values of Protein Pairs and Cancer Types

For the proteins which were defined as differentially expressed in TCPA or HPA datasets, weighted ratio values based on differential expression, positive or negative correlation and mutually antagonistic relationships were calculated. Differential expression number of each protein was defined as the number of CS (or CT) for which differential expression was observed.

Positive correlation based protein pairs (PCPP) and negative correlation based protein pairs (NCPP) were analyzed. For correlation analysis, Jaccard index based correlation value for each protein was calculated as below.

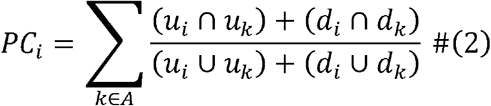

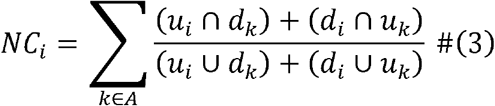

where PC_i_ is the positive correlation value, and NC_i_ is the negative correlation value of protein i, u_i_ and u_k_ are the sets of CS (or CT) in which protein i and protein k are ON, d_i_ and d_k_ are the sets of CS (or CT) in which protein i and protein k are OFF, A is the set of all proteins except for the protein i.

Finally, the mutually antagonistic value (MAPP number) for each protein was calculated the sum of all pairs in which the protein was defined as being in a mutually antagonistic pair. Ratio values for differential expression, correlation and mutual antagonism of each protein were calculated by dividing the individual value of each protein to the sum for all proteins. Differential ratio values for PCPP, NCPP and MAPP were calculated as the difference of the correlation and the mutual antagonism based values from the differential expression based values. Both CS and CT based values were calculated for TCPA and only CT based values were calculated for HPA.

### Molecular Interactions of MAPP

Molecular interaction neighborhood of significant MAPP was analyzed for both TCPA and HPA datasets. BioGrid Physical Interaction Dataset Version 4.3.194 was used for human protein-protein interactions (Stark et al. 2006, Oughtred et al. 2021). TRRUST (release note 2018.04.16) and The Human Transcriptional Regulation Interactions database (HTRIdb) were used for human transcriptional (PDI) interactions (Han et al. 2015, Han et al. 2018, Bovolenta et al. 2012). Antibody based protein names of TCPA were converted to NCBI Gene Symbols, which could be integrated with protein-protein and transcriptional interactions (Supplementary Table 12). Direct (first degree) and indirect (second degree) protein-protein interactions of MAPP were explored. One TCPA name could match to a maximum of 3 different NCBI Gene Symbols, therefore all possible interactions of the matching Symbols were considered separately. For transcriptional interactions, TRRUST and HTRIdb database pairs were combined in order to have a more comprehensive dataset. Then, directional transcription factor-gene interactions of MAPP were explored.

### Software and Code Availability

R scripts are available at https://github.com/ertuda/MAPP. Network visualization was done by Cytoscape (Shannon et al. 2003).

