## Supplemental Information for "Mutually Antagonistic Protein Pairs of Cancer"

### Supplementary Information

### Supplementary Methods

TCPA pan-cancer L4 dataset has 7694 cancer sample values for 258 proteins (antibodies). The proteins with more than 50% missing values were removed. The remaining dataset has 217 proteins. The expression values range from -5.6 to 19.5 with a median of 0. Median values for different samples range from -0.2 to 0.1. 95% of all values range from -1.3 to 1.5 (Supplementary Figure 4). The list of 217 proteins includes different modifications of the same protein, as specific antibodies target differentially modified proteins and they could have distinct biological functions. For example, CHK1_pS345, P27_pT198, SRC_pY527, etc. are present in addition to the unmodified version of the same proteins. If such modifications are neglected, there are 175 unique proteins in the dataset.

Different absolute threshold values for defining differential expression were tested. The threshold of 1 selected around 10% of all values, whereas the threshold of 2 selected around 2% (Supplementary Figure 4). As the threshold value increased the fraction of selected values decreased below 1%. The absolute threshold value of 2 (-2 for lower expression and 2 for higher expression) was chosen to define differentially expressed proteins. Proteins with an expression value higher than 2 were defined as ON and the ones with an expression value lower than -2 were defined as OFF for each cancer sample. First, mutually antagonistic protein pairs were defined based on the absolute threshold value by ignoring the cancer type of the sample. For example, MYH11 is ON and HER2 is OFF in the samples; TCGA-DD-A4NH-01A-21-A40M-20, TCGA-DD-A3A6-01A-21-A40L-20, and TCGA-FD-A6TB-01A-21-A458-20. While the cancer types of the first two samples are Liver hepatocellular carcinoma (LIHC), the cancer type of the third sample is Bladder Urothelial Carcinoma (BLCA). MYH11 is OFF and HER2 is ON in the sample TCGA-D8-A1JA-01A-21-A17J-20, which belongs to Breast invasive carcinoma (BRCA). When the cancer types are ignored, there are 3 pairs of cancer samples in which MYH11-HER2 pair is ON-OFF in the first case, and OFF-ON in the second case. All the protein pairs were explored throughout the dataset similarly, keeping the number of their occurrences which for instance is 3 for the MYH11-HER2 pair.

Permutation tests were performed by shuffling the expression values of each cancer sample. Mutually antagonist protein pairs were similarly explored in the randomized datasets. Significance of the cancer sample based protein pairs was tested by comparing the number of occurrences of each protein pair to their random occurrences, and p-values were defined as the fraction of random iterations with an equal or higher number of samples pair occurrences. Multiple testing correction of the p-values was done by False Discovery Rate (FDR).

Next, mutually antagonistic protein pairs were defined based on the cancer type of the sample. For example, MYH11 is ON and HER2 is OFF in 3 different samples but 2 different cancer types; LIHC and BLCA. MYH11 is OFF and HER2 is ON in only one sample which belongs to BRCA. Therefore, there are 2 unique cases where MYH11-HER2 is ON-OFF and OFF-ON, when cancer types are considered, between the cancer type pairs LIHC-BRCA and BLCA-BRCA. All the protein pairs were explored and compared to randomized datasets similarly. Significance of the cancer type based protein pairs were analyzed similarly.

The pathology dataset of HPA, has protein expression values for 19651 proteins, across 20 cancer types. The dataset has as different number of ‘not detected’, ‘low’, ‘medium’, and ‘high’ measurements for proteins. It has 15308 proteins after removing the ones with missing values. Total number of scoring values for different rows (measurements of a proteins in a cancer type) range from 1 to 12. Some proteins also have very few values. Different threshold values were tested for their effect on the dataset (Supplementary Figure 5). There is a significant drop in the number of all values in the dataset after the threshold of 10, whereas the coverage of genes does not change. Therefore rows (proteins for all cancer types) with less than 10 values were removed from the pathology dataset of HPA. After the removal, the dataset has expression values for 15308 proteins and 17 cancer types. Equation 1 in the Methods section was used to define the Cancer Expression parameter (CE) for the tumor samples of HPA (the pathology dataset). CE has a range of -1 to 1. An absolute threshold value for CE is used to assign binary values (present/absent) for all proteins in different cancer types. CE has a w factor which alters the effect of ‘high’ and ‘not detected’ counts as compared to ‘low’ and ‘medium’ counts. Different values for the absolute CE threshold and w were chosen to observe the number of proteins with binary values (fraction of selected rows) and MAPP. As the absolute CE threshold value increase from 0.1 to 1 the fraction of the binary proteins (including all instances for each cancer type) decrease from above 90% to lower than 50%, with mostly higher variance (Supplementary Figure 5). Also, the number of MAPP decrease from above 2500 to 152-273 in the CE threshold range of 0.1-0.5, and from 57-93 to 1-5 in the range of 0.6-1 (Supplementary Figure 5). The decline is sharper and the variation is larger before the threshold of 0.5. Weight parameter has a mild effect on the number of binary proteins and MAPP (Supplementary Figures 5). Variation of the number of MAPP increase as w increase from 2-5. As a result, the absolute CE threshold value was set to 0.5 (-0.5 for absence and 0.5 for presence) and w was set to 2.

The normal expression dataset of HPA has 15308 proteins across 63 tissue types and 120 cell types. Cancer types of the pathology dataset were manually matched to the tissues and cell types of the normal expression dataset (Supplementary Tables 13-14). Some cancer types like breast cancer can match to a single tissue but multiple cell types; i.e., glandular cells and myoepithelial cells. Some cancer types like glioma can match to a single cell type but multiple tissues; i.e., caudate, cerebral cortex, hippocampus, etc. When there were multiple normal values for a single cancer type as in the case of breast cancer and glioma, differential expression with at least one of them was required. For example, TSPAN6 protein has 2 ‘not detected’, 2 ‘low’, 7 ‘medium’, and 1 ‘high’ scoring values for breast cancer. On the other hand, TSPAN6 protein is labeled as ‘high’ in glandular cells, and ‘not detected’ in myoepithelial cells of normal breast tissue. CE value of TSPAN6 in breast cancer is 0.2 therefore it is not defined as binary as it is below the 0.5 CE threshold value. TSPAN6 protein has 0 ‘not detected’, 1 ‘low’, 5 ‘medium’, and 4 ‘high’ scoring values for liver cancer, and it is labeled as ‘not detected’ in hepatocytes of normal liver tissue. CE value of TSPAN6 in liver cancer is 0.86, therefore TSPAN is defined a binary (present) in liver cancer. Its normal value is binary since the expression in hepatocytes is defined as ‘not detected’. TSPAN is defined as differentially expressed (upregulated) since it is present in liver cancer and absent in normal liver. TSPAN6 protein has 0 ‘not detected’, 0 ‘low’, 2 ‘medium’, and 10 ‘high’ scoring values for endometrial cancer, and it is labeled as ‘high’ in glandular cells of normal endometrium. Although its CE value is 1 (thus present in cancer), TSPAN is not defined as differentially expressed (upregulated) in endometrial cancer, since it is also present in normal.

### Supplementary Figures

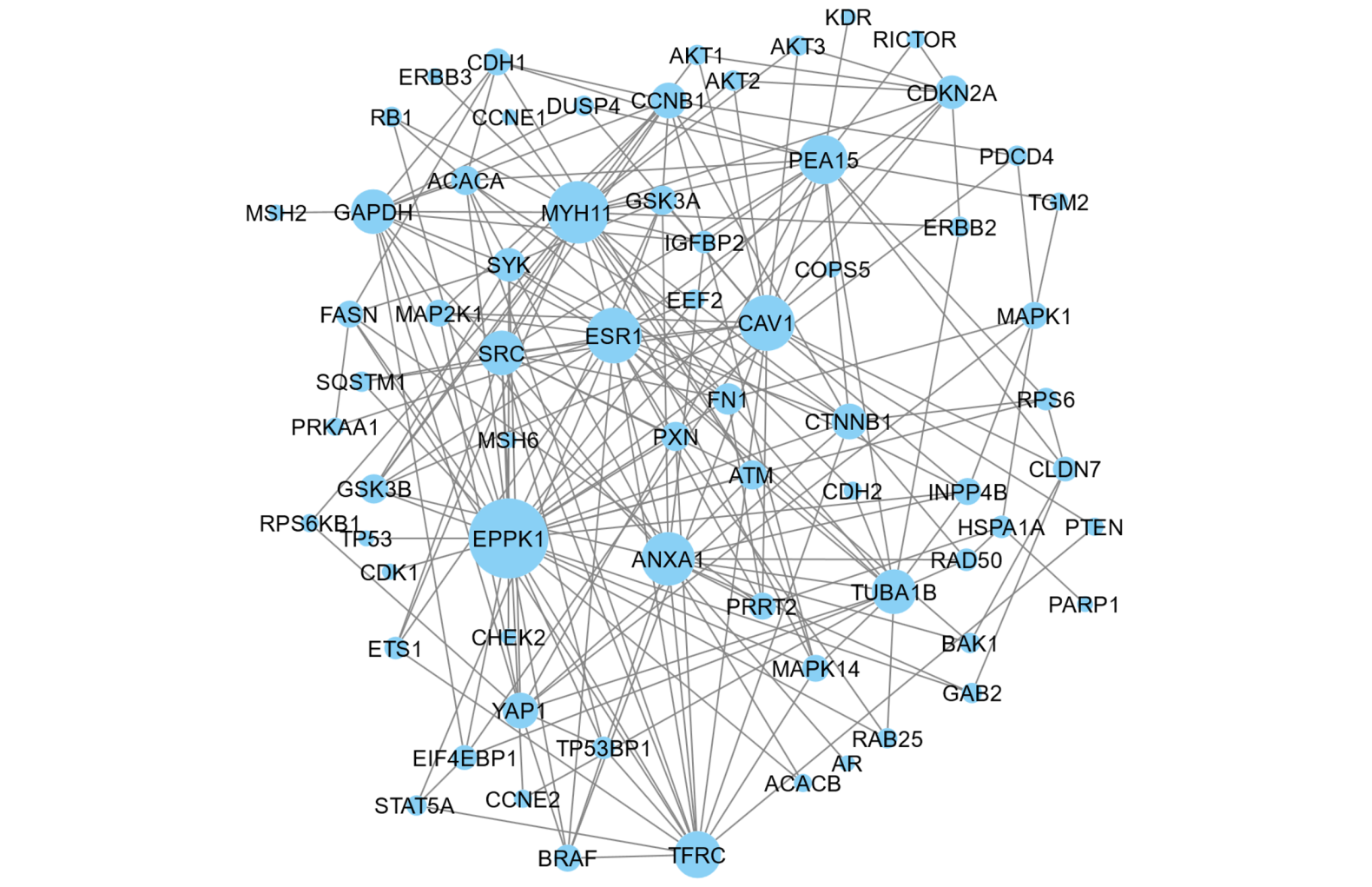

**Supplementary Figure 1.** Significant MAPP network of TCPA. CS and CT based MAPP were combined and unique pairs were built into a network. TCPA names were converted to NCBI symbols. Size of the nodes are correlated to their degree values (number of links).

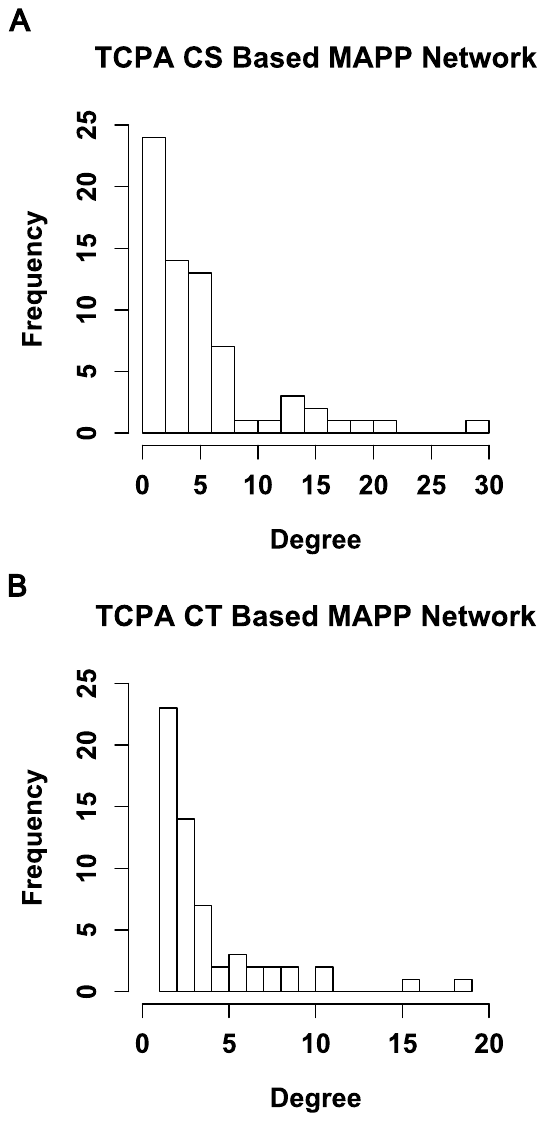

**Supplementary Figure 2.** Degree distribution of **(A)** the CS based significant MAPP network and **(B)** the CT based significant MAPP network.

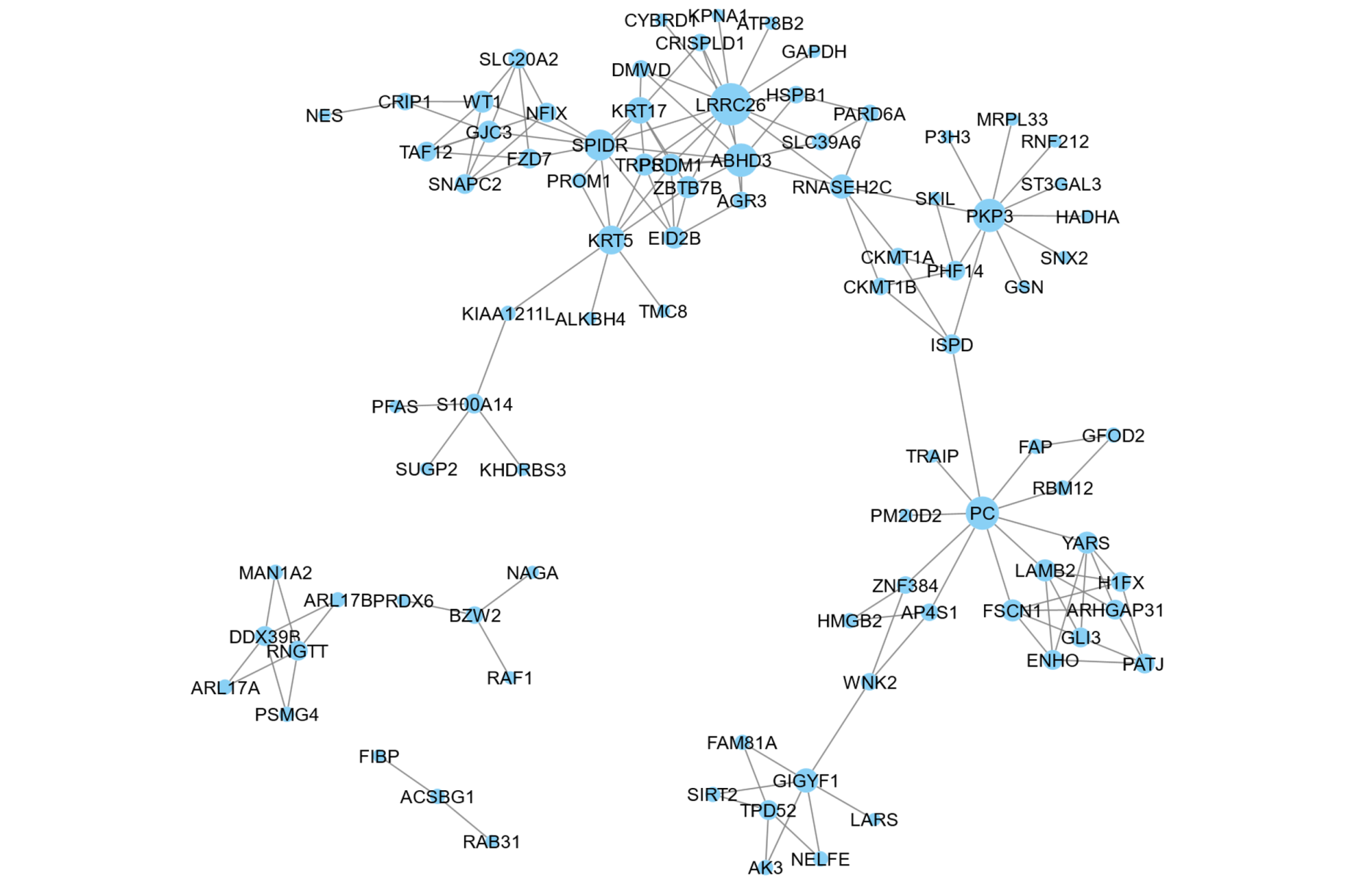

**Supplementary Figure 3.** Significant MAPP network of HPA. Size of the nodes are correlated to their degree values (number of links).

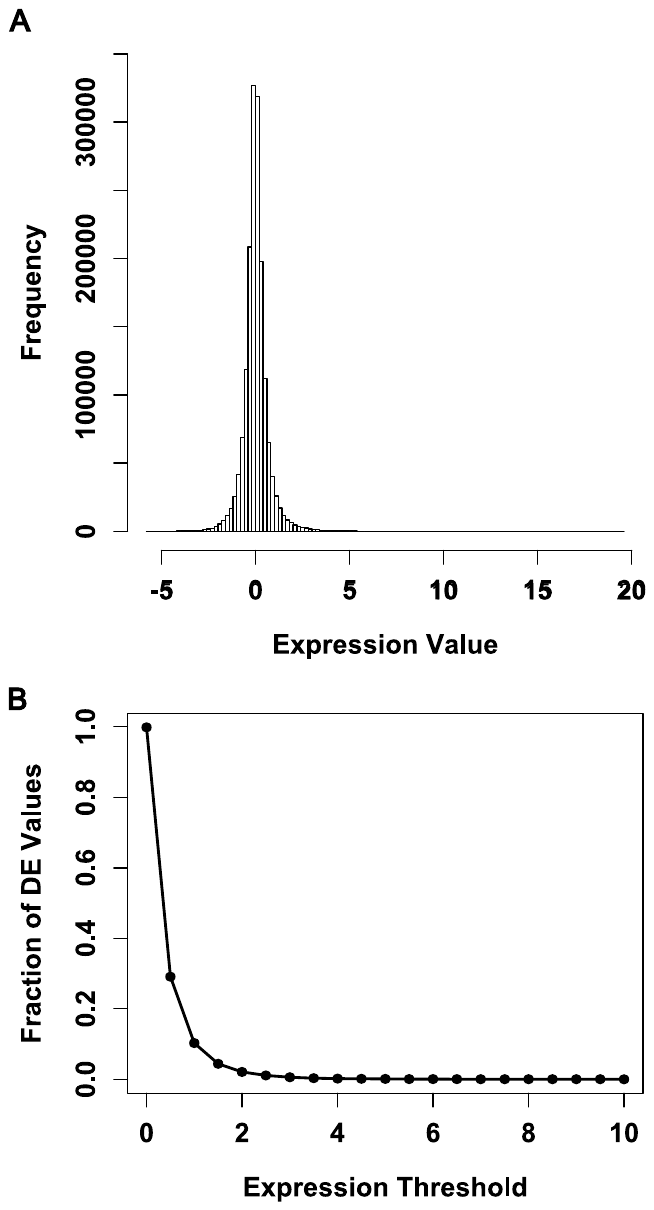

**Supplementary Figure 4. Exploratory analysis of the preprocessing steps of the TCPA dataset (A)** Distribution of expression values for TCPA after removing proteins with more than 50% missing values. **(B)** Fraction of differential expression values were calculated for various expression threshold values.

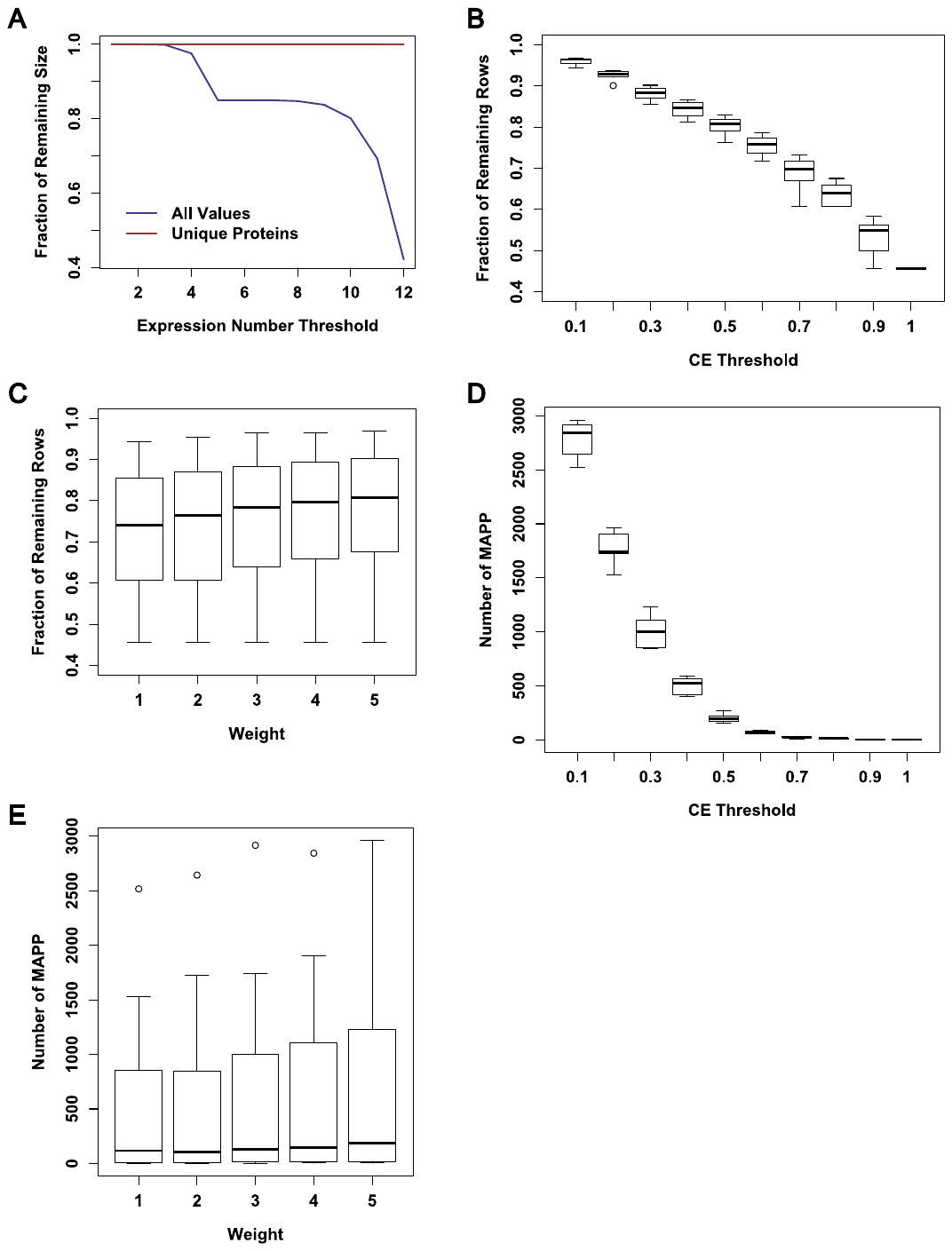

**Supplementary Figure 5. Exploratory analysis of the preprocessing steps of the HPA dataset.** **(A)** Fraction of remaining number of all values of the pathology dataset of HPA and unique HPA proteins based on different Expression Number Threshold values of removal. **(B)** Fraction of remaining rows of the pathology dataset of HPA based on different absolute CE Threshold values. For each CE Threshold value, different w parameters were used. **(C)** Number of MAPP generated by using the pathology dataset of HPA based on different absolute CE Threshold values. For each CE Threshold value, different w parameters were used. **(D)** Fraction of remaining rows of the pathology dataset of HPA based on different weight (w) parameter values. For each w, different CE Threshold values were used. **(E)** Number of MAPP generated by using the pathology dataset of HPA based on different weight (w) parameter values. For each w, different CE Threshold values were used.

### Supplementary Tables

**Supplementary Table 1.** TCPA MAPP list based on CS and CT. For each unique protein pair, number of occurrences (count) and p-values were given.

| Protein 1 | Protein 2 | CS count | CS p-value | CS FDR | CT count | CT p-value | CT FDR |
| --- | --- | --- | --- | --- | --- | --- | --- |
| X4EBP1_pT37T46 | STAT5ALPHA | 2 | 0.001 | 0.002426 | 2 | 0.001 | 0.007104 |
| X4EBP1_pT37T46 | EPPK1 | 96 | 0.001 | 0.002426 | 12 | 0.001 | 0.007104 |
| X53BP1 | YAP_pS127 | 1 | 0.001 | 0.002426 | 1 | 0.001 | 0.007104 |
| ATM | SRC_pY416 | 5 | 0.001 | 0.002426 | 2 | 0.001 | 0.007104 |
| ATM | SRC_pY527 | 4 | 0.001 | 0.002426 | 2 | 0.001 | 0.007104 |
| BAK | CLAUDIN7 | 2 | 0.001 | 0.002426 | 1 | 0.001 | 0.007104 |
| BETACATENIN | YAP_pS127 | 2 | 0.001 | 0.002426 | 2 | 0.001 | 0.007104 |
| CAVEOLIN1 | PTEN | 4 | 0.001 | 0.002426 | 3 | 0.001 | 0.007104 |
| CAVEOLIN1 | BRAF | 80 | 0.001 | 0.002426 | 8 | 0.001 | 0.007104 |
| CAVEOLIN1 | PKCPANBETAII_pS660 | 63 | 0.001 | 0.002426 | 1 | 0.001 | 0.007104 |
| CAVEOLIN1 | TFRC | 168 | 0.001 | 0.002426 | 16 | 0.001 | 0.007104 |
| CAVEOLIN1 | DUSP4 | 1 | 0.001 | 0.002426 | 1 | 0.001 | 0.007104 |
| CLAUDIN7 | S6_pS240S244 | 2 | 0.001 | 0.002426 | 1 | 0.001 | 0.007104 |
| CLAUDIN7 | PEA15_pS116 | 18 | 0.001 | 0.002426 | 3 | 0.001 | 0.007104 |
| CYCLINB1 | SYK | 1 | 0.001 | 0.002426 | 1 | 0.001 | 0.007104 |
| CYCLINE1 | MYH11 | 2 | 0.001 | 0.002426 | 1 | 0.001 | 0.007104 |
| EEF2 | PEA15_pS116 | 1 | 0.001 | 0.002426 | 1 | 0.001 | 0.007104 |
| ERALPHA | PAXILLIN | 1 | 0.001 | 0.002426 | 1 | 0.001 | 0.007104 |
| ERALPHA | STAT5ALPHA | 8 | 0.001 | 0.002426 | 2 | 0.001 | 0.007104 |
| ERK2 | PDCD4 | 13 | 0.001 | 0.002426 | 1 | 0.001 | 0.007104 |
| ERK2 | TRANSGLUTAMINASE | 1 | 0.001 | 0.002426 | 1 | 0.001 | 0.007104 |
| FIBRONECTIN | EPPK1 | 26 | 0.001 | 0.002426 | 8 | 0.001 | 0.007104 |
| HER3_pY1289 | MYH11 | 1 | 0.001 | 0.002426 | 1 | 0.001 | 0.007104 |
| HSP70 | PARPCLEAVED | 1 | 0.001 | 0.002426 | 1 | 0.001 | 0.007104 |
| IGFBP2 | MYH11 | 10 | 0.001 | 0.002426 | 3 | 0.001 | 0.007104 |
| MEK1 | YAP_pS127 | 3 | 0.001 | 0.002426 | 1 | 0.001 | 0.007104 |
| P38_pT180Y182 | MYH11 | 1 | 0.001 | 0.002426 | 1 | 0.001 | 0.007104 |
| PAXILLIN | SRC_pY416 | 15 | 0.001 | 0.002426 | 6 | 0.001 | 0.007104 |
| PAXILLIN | EPPK1 | 110 | 0.001 | 0.002426 | 24 | 0.001 | 0.007104 |
| PTEN | TFRC | 1 | 0.001 | 0.002426 | 1 | 0.001 | 0.007104 |
| RB_pS807S811 | MYH11 | 42 | 0.001 | 0.002426 | 8 | 0.001 | 0.007104 |
| SRC_pY416 | ETS1 | 1 | 0.001 | 0.002426 | 1 | 0.001 | 0.007104 |
| SRC_pY527 | PEA15_pS116 | 1 | 0.001 | 0.002426 | 1 | 0.001 | 0.007104 |
| SYK | FASN | 2 | 0.001 | 0.002426 | 2 | 0.001 | 0.007104 |
| SYK | EPPK1 | 30 | 0.001 | 0.002426 | 3 | 0.001 | 0.007104 |
| VEGFR2 | PEA15_pS116 | 36 | 0.001 | 0.002426 | 2 | 0.001 | 0.007104 |
| YAP_pS127 | BRAF | 1 | 0.001 | 0.002426 | 1 | 0.001 | 0.007104 |
| GSK3_pS9 | MYH11 | 12 | 0.001 | 0.002426 | 7 | 0.001 | 0.007104 |
| MYH11 | P16INK4A | 20 | 0.001 | 0.002426 | 12 | 0.001 | 0.007104 |
| PEA15_pS116 | RICTOR | 2 | 0.001 | 0.002426 | 1 | 0.001 | 0.007104 |
| PEA15_pS116 | DUSP4 | 4 | 0.001 | 0.002426 | 2 | 0.001 | 0.007104 |
| TFRC | ANNEXIN1 | 35 | 0.001 | 0.002426 | 3 | 0.001 | 0.007104 |
| EPPK1 | CDK1_pY15 | 2 | 0.001 | 0.002426 | 2 | 0.001 | 0.007104 |
| ATM | P38_pT180Y182 | 2 | 0.001 | 0.002426 | 2 | 0.002 | 0.011559 |
| BETACATENIN | MYH11 | 234 | 0.001 | 0.002426 | 33 | 0.002 | 0.011559 |
| ERALPHA | ACETYLATUBULINLYS40 | 448 | 0.001 | 0.002426 | 12 | 0.002 | 0.011559 |
| FIBRONECTIN | MYH11 | 2 | 0.001 | 0.002426 | 2 | 0.002 | 0.011559 |
| INPP4B | EPPK1 | 11 | 0.001 | 0.002426 | 6 | 0.002 | 0.011559 |
| MEK1 | GAPDH | 1 | 0.001 | 0.002426 | 1 | 0.002 | 0.011559 |
| STAT5ALPHA | TFRC | 2 | 0.001 | 0.002426 | 1 | 0.002 | 0.011559 |
| BRAF | TFRC | 6 | 0.001 | 0.002426 | 6 | 0.002 | 0.011559 |
| BRAF | EPPK1 | 256 | 0.001 | 0.002426 | 10 | 0.002 | 0.011559 |
| ACC_pS79 | EPPK1 | 1 | 0.001 | 0.002426 | 1 | 0.003 | 0.014826 |
| ERALPHA | P16INK4A | 6 | 0.001 | 0.002426 | 4 | 0.003 | 0.014826 |
| ERK2 | HSP70 | 6 | 0.001 | 0.002426 | 2 | 0.003 | 0.014826 |
| P38_pT180Y182 | TFRC | 2 | 0.001 | 0.002426 | 1 | 0.003 | 0.014826 |
| PAXILLIN | RAB25 | 9 | 0.001 | 0.002426 | 3 | 0.003 | 0.014826 |
| FASN | EPPK1 | 399 | 0.001 | 0.002426 | 27 | 0.003 | 0.014826 |
| PAXILLIN | PEA15_pS116 | 8 | 0.001 | 0.002426 | 4 | 0.004 | 0.018432 |
| YAP_pS127 | ACETYLATUBULINLYS40 | 161 | 0.001 | 0.002426 | 9 | 0.004 | 0.018432 |
| RAB25 | EPPK1 | 6 | 0.001 | 0.002426 | 4 | 0.004 | 0.018432 |
| X4EBP1_pT37T46 | ACETYLATUBULINLYS40 | 2 | 0.001 | 0.002426 | 2 | 0.005 | 0.021859 |
| GAB2 | EPPK1 | 19 | 0.001 | 0.002426 | 5 | 0.005 | 0.021859 |
| SRC_pY527 | MYH11 | 4 | 0.001 | 0.002426 | 2 | 0.005 | 0.021859 |
| RICTOR | P16INK4A | 3 | 0.001 | 0.002426 | 3 | 0.005 | 0.021859 |
| ACC1 | TFRC | 7 | 0.001 | 0.002426 | 2 | 0.006 | 0.024951 |
| AKT_pT308 | MYH11 | 9 | 0.001 | 0.002426 | 3 | 0.006 | 0.024951 |
| P70S6K_pT389 | MYH11 | 5 | 0.001 | 0.002426 | 1 | 0.006 | 0.024951 |
| ETS1 | MYH11 | 6 | 0.001 | 0.002426 | 6 | 0.006 | 0.024951 |
| TFRC | EPPK1 | 1007 | 0.001 | 0.002426 | 95 | 0.007 | 0.028417 |
| X53BP1 | TFRC | 10 | 0.001 | 0.002426 | 5 | 0.008 | 0.031 |
| ERALPHA | PKCPANBETAII_pS660 | 2 | 0.001 | 0.002426 | 1 | 0.008 | 0.031 |
| GSK3ALPHABETA_pS21S9 | SYK | 6 | 0.001 | 0.002426 | 2 | 0.011 | 0.040772 |
| SRC_pY416 | EPPK1 | 69 | 0.001 | 0.002426 | 12 | 0.011 | 0.040772 |
| ATM | ECADHERIN | 13 | 0.001 | 0.002426 | 4 | 0.012 | 0.044 |
| P38_pT180Y182 | EPPK1 | 3 | 0.001 | 0.002426 | 2 | 0.013 | 0.045701 |
| P53 | EPPK1 | 3 | 0.001 | 0.002426 | 3 | 0.013 | 0.045701 |
| YAP_pS127 | GAPDH | 5 | 0.001 | 0.002426 | 5 | 0.013 | 0.045701 |
| FASN | PKCPANBETAII_pS660 | 1 | 0.001 | 0.002426 | 1 | 0.013 | 0.045701 |
| X53BP1 | EPPK1 | 12 | 0.001 | 0.002426 | 4 | 0.014 | 0.046804 |
| ERALPHA | GAPDH | 2108 | 0.001 | 0.002426 | 20 | 0.014 | 0.046804 |
| BRAF | ANNEXIN1 | 128 | 0.001 | 0.002426 | 2 | 0.014 | 0.046804 |
| CAVEOLIN1 | SRC_pY527 | 44 | 0.001 | 0.002426 | 3 | 0.015 | 0.048255 |
| SRC_pY527 | EPPK1 | 116 | 0.001 | 0.002426 | 17 | 0.015 | 0.048255 |
| ERALPHA | BRAF | 2 | 0.001 | 0.002426 | 2 | 0.016 | 0.049153 |
| EPPK1 | P62LCKLIGAND | 12 | 0.001 | 0.002426 | 7 | 0.016 | 0.049153 |
| ACETYLATUBULINLYS40 | ANNEXIN1 | 3100 | 0.001 | 0.002426 | 12 | 0.016 | 0.049153 |
| HER2 | P16INK4A | 5 | 0.001 | 0.002426 | 4 | 0.019 | 0.057336 |
| ERALPHA | MEK1 | 2 | 0.001 | 0.002426 | 2 | 0.022 | 0.065235 |
| CLAUDIN7 | GAB2 | 1 | 0.001 | 0.002426 | 1 | 0.025 | 0.072246 |
| S6_pS240S244 | PEA15_pS116 | 2 | 0.001 | 0.002426 | 1 | 0.025 | 0.072246 |
| CAVEOLIN1 | INPP4B | 2 | 0.001 | 0.002426 | 1 | 0.031 | 0.087364 |
| ACC1 | BETACATENIN | 3 | 0.001 | 0.002426 | 1 | 0.034 | 0.093242 |
| ATM | MYH11 | 17 | 0.001 | 0.002426 | 7 | 0.035 | 0.093242 |
| CAVEOLIN1 | JAB1 | 4 | 0.001 | 0.002426 | 1 | 0.035 | 0.093242 |
| CAVEOLIN1 | CLAUDIN7 | 3 | 0.001 | 0.002426 | 1 | 0.036 | 0.09371 |
| GSK3ALPHABETA_pS21S9 | ANNEXIN1 | 12 | 0.001 | 0.002426 | 3 | 0.036 | 0.09371 |
| GAPDH | EPPK1 | 2241 | 0.001 | 0.002426 | 69 | 0.037 | 0.095583 |
| AR | ANNEXIN1 | 2 | 0.001 | 0.002426 | 2 | 0.038 | 0.095985 |
| EPPK1 | P16INK4A | 44 | 0.001 | 0.002426 | 9 | 0.038 | 0.095985 |
| BAK | ANNEXIN1 | 17 | 0.001 | 0.002426 | 2 | 0.039 | 0.097429 |
| RAD50 | ACETYLATUBULINLYS40 | 10 | 0.001 | 0.002426 | 3 | 0.04 | 0.097429 |
| CYCLINE2 | ACETYLATUBULINLYS40 | 1 | 0.001 | 0.002426 | 1 | 0.042 | 0.101574 |
| FIBRONECTIN | ACETYLATUBULINLYS40 | 64 | 0.001 | 0.002426 | 10 | 0.049 | 0.112899 |
| PAXILLIN | SRC_pY527 | 1 | 0.001 | 0.002426 | 1 | 0.049 | 0.112899 |
| CYCLINB1 | P38_pT180Y182 | 12 | 0.001 | 0.002426 | 8 | 0.05 | 0.113667 |
| NCADHERIN | ACETYLATUBULINLYS40 | 1 | 0.001 | 0.002426 | 1 | 0.05 | 0.113667 |
| SYK | YAP_pS127 | 1 | 0.001 | 0.002426 | 1 | 0.052 | 0.11743 |
| ERALPHA | GSK3ALPHABETA_pS21S9 | 4 | 0.001 | 0.002426 | 3 | 0.068 | 0.148641 |
| CYCLINB1 | MEK1 | 35 | 0.001 | 0.002426 | 2 | 0.07 | 0.151076 |
| GAPDH | DUSP4 | 4 | 0.001 | 0.002426 | 4 | 0.074 | 0.156733 |
| BAK | ERALPHA | 4 | 0.001 | 0.002426 | 1 | 0.076 | 0.158024 |
| ERK2 | YAP_pS127 | 6 | 0.001 | 0.002426 | 1 | 0.077 | 0.158175 |
| GSK3_pS9 | EPPK1 | 28 | 0.001 | 0.002426 | 15 | 0.085 | 0.161927 |
| HSP70 | RAD50 | 1 | 0.001 | 0.002426 | 1 | 0.088 | 0.16579 |
| CAVEOLIN1 | PEA15_pS116 | 1998 | 0.001 | 0.002426 | 6 | 0.091 | 0.169568 |
| INPP4B | SYK | 1 | 0.001 | 0.002426 | 1 | 0.104 | 0.191697 |
| X4EBP1_pT37T46 | CAVEOLIN1 | 3 | 0.001 | 0.002426 | 2 | 0.196 | 0.300021 |
| X53BP1 | SRC_pY527 | 1 | 0.001 | 0.002426 | 1 | 0.18 | 0.300021 |
| AMPKALPHA_pT172 | ERALPHA | 1 | 0.001 | 0.002426 | 1 | 0.205 | 0.300021 |
| AMPKALPHA_pT172 | FASN | 4 | 0.001 | 0.002426 | 1 | 0.191 | 0.300021 |
| BETACATENIN | RAD50 | 1 | 0.001 | 0.002426 | 1 | 0.183 | 0.300021 |
| BETACATENIN | EPPK1 | 660 | 0.001 | 0.002426 | 32 | 0.192 | 0.300021 |
| CYCLINB1 | ECADHERIN | 12 | 0.001 | 0.002426 | 5 | 0.195 | 0.300021 |
| FIBRONECTIN | SRC_pY527 | 1 | 0.001 | 0.002426 | 1 | 0.181 | 0.300021 |
| HER2 | ACETYLATUBULINLYS40 | 6 | 0.001 | 0.002426 | 3 | 0.176 | 0.300021 |
| IGFBP2 | GAPDH | 37 | 0.001 | 0.002426 | 13 | 0.181 | 0.300021 |
| PEA15_pS116 | TRANSGLUTAMINASE | 7 | 0.001 | 0.002426 | 3 | 0.189 | 0.300021 |
| MYH11 | ANNEXIN1 | 27 | 0.001 | 0.002426 | 8 | 0.208 | 0.301821 |
| ATM | EPPK1 | 44 | 0.001 | 0.002426 | 12 | 0.369 | 0.380346 |
| BETACATENIN | S6_pS235S236 | 1 | 0.001 | 0.002426 | 1 | 0.333 | 0.380346 |
| CHK2 | EPPK1 | 1 | 0.001 | 0.002426 | 1 | 0.356 | 0.380346 |
| CYCLINB1 | PDCD4 | 6 | 0.001 | 0.002426 | 1 | 0.333 | 0.380346 |
| ECADHERIN | FASN | 50 | 0.001 | 0.002426 | 10 | 0.36 | 0.380346 |
| ERK2 | FIBRONECTIN | 8 | 0.001 | 0.002426 | 4 | 0.323 | 0.380346 |
| INPP4B | ANNEXIN1 | 3 | 0.001 | 0.002426 | 3 | 0.343 | 0.380346 |
| P38_pT180Y182 | ACETYLATUBULINLYS40 | 1 | 0.001 | 0.002426 | 1 | 0.374 | 0.380346 |
| PAXILLIN | TFRC | 10 | 0.001 | 0.002426 | 3 | 0.355 | 0.380346 |
| S6_pS240S244 | EPPK1 | 15 | 0.001 | 0.002426 | 9 | 0.35 | 0.380346 |
| SRC_pY527 | GAPDH | 24 | 0.001 | 0.002426 | 12 | 0.336 | 0.380346 |
| HSP70 | PKCPANBETAII_pS660 | 2 | 0.001 | 0.002426 | 0 | 1 | 1 |
| CAVEOLIN1 | MEK1 | 25 | 0.002 | 0.004653 | 1 | 0.004 | 0.018432 |
| ERALPHA | SYK | 8 | 0.002 | 0.004653 | 2 | 0.008 | 0.031 |
| AKT_pS473 | CAVEOLIN1 | 11 | 0.002 | 0.004653 | 2 | 0.014 | 0.046804 |
| IGFBP2 | ANNEXIN1 | 10 | 0.002 | 0.004653 | 5 | 0.176 | 0.300021 |
| GAPDH | MYH11 | 1424 | 0.002 | 0.004653 | 45 | 0.353 | 0.380346 |
| SRC_pY527 | TFRC | 16 | 0.002 | 0.004653 | 9 | 0.383 | 0.384126 |
| SRC_pY416 | ANNEXIN1 | 16 | 0.003 | 0.006795 | 3 | 0.04 | 0.097429 |
| ACC1 | PEA15_pS116 | 15 | 0.003 | 0.006795 | 7 | 0.044 | 0.103476 |
| X4EBP1_pT37T46 | GAPDH | 5 | 0.003 | 0.006795 | 3 | 0.195 | 0.300021 |
| GAPDH | MSH2 | 2 | 0.003 | 0.006795 | 1 | 0.362 | 0.380346 |
| CAVEOLIN1 | P16INK4A | 10 | 0.004 | 0.008713 | 10 | 0.002 | 0.011559 |
| PAXILLIN | YAP_pS127 | 1 | 0.004 | 0.008713 | 1 | 0.007 | 0.028417 |
| RAB25 | ACETYLATUBULINLYS40 | 24 | 0.004 | 0.008713 | 6 | 0.026 | 0.073883 |
| CYCLINB1 | GAPDH | 150 | 0.004 | 0.008713 | 4 | 0.084 | 0.161831 |
| IGFBP2 | PEA15_pS116 | 21 | 0.004 | 0.008713 | 2 | 0.092 | 0.1705 |
| BETACATENIN | ERALPHA | 1380 | 0.004 | 0.008713 | 10 | 0.334 | 0.380346 |
| GSK3_pS9 | ANNEXIN1 | 14 | 0.005 | 0.010755 | 5 | 0.001 | 0.007104 |
| PEA15_pS116 | ACETYLATUBULINLYS40 | 5 | 0.005 | 0.010755 | 5 | 0.038 | 0.095985 |
| FIBRONECTIN | RB_pS807S811 | 3 | 0.006 | 0.012667 | 1 | 0.01 | 0.037889 |
| RAD50 | ANNEXIN1 | 6 | 0.006 | 0.012667 | 1 | 0.081 | 0.160587 |
| PKCPANBETAII_pS660 | ANNEXIN1 | 480 | 0.006 | 0.012667 | 3 | 0.361 | 0.380346 |
| ATM | ACETYLATUBULINLYS40 | 27 | 0.007 | 0.014687 | 4 | 0.067 | 0.1474 |
| ECADHERIN | GAPDH | 595 | 0.008 | 0.016683 | 14 | 0.032 | 0.088715 |
| SRC_pY416 | MYH11 | 54 | 0.009 | 0.018431 | 11 | 0.003 | 0.014826 |
| ATM | ANNEXIN1 | 5 | 0.009 | 0.018431 | 3 | 0.082 | 0.160701 |
| CAVEOLIN1 | GSK3ALPHABETA_pS21S9 | 9 | 0.009 | 0.018431 | 3 | 0.174 | 0.300021 |
| PDCD4 | EPPK1 | 90 | 0.01 | 0.020237 | 19 | 0.076 | 0.158024 |
| NCADHERIN | MYH11 | 10 | 0.01 | 0.020237 | 4 | 0.172 | 0.300021 |
| FASN | TFRC | 256 | 0.011 | 0.022 | 9 | 0.044 | 0.103476 |
| AKT_pS473 | MYH11 | 22 | 0.011 | 0.022 | 4 | 0.067 | 0.1474 |
| ERALPHA | TFRC | 1056 | 0.012 | 0.023723 | 29 | 0.02 | 0.059825 |
| SRC_pY527 | CYCLINE2 | 1 | 0.012 | 0.023723 | 1 | 0.079 | 0.160351 |
| CAVEOLIN1 | P62LCKLIGAND | 2 | 0.013 | 0.025406 | 2 | 0.197 | 0.300021 |
| CYCLINB1 | GSK3_pS9 | 6 | 0.013 | 0.025406 | 3 | 0.345 | 0.380346 |
| HER2 | MYH11 | 3 | 0.014 | 0.027205 | 2 | 0.178 | 0.300021 |
| EEF2 | EPPK1 | 6 | 0.017 | 0.032847 | 4 | 0.016 | 0.049153 |
| INPP4B | ACETYLATUBULINLYS40 | 12 | 0.019 | 0.036506 | 2 | 0.204 | 0.300021 |
| BETACATENIN | TFRC | 152 | 0.021 | 0.040123 | 15 | 0.035 | 0.093242 |
| RB_pS807S811 | EPPK1 | 204 | 0.023 | 0.043459 | 10 | 0.066 | 0.147098 |
| GAB2 | ANNEXIN1 | 14 | 0.023 | 0.043459 | 10 | 0.362 | 0.380346 |
| MYH11 | PEA15_pS116 | 700 | 0.024 | 0.044609 | 9 | 0.032 | 0.088715 |
| ERALPHA | ETS1 | 29 | 0.024 | 0.044609 | 4 | 0.186 | 0.300021 |
| ETS1 | TFRC | 28 | 0.024 | 0.044609 | 5 | 0.364 | 0.380346 |
| ACC1 | ECADHERIN | 3 | 0.025 | 0.046216 | 2 | 0.003 | 0.014826 |
| ACC_pS79 | ANNEXIN1 | 1 | 0.026 | 0.047806 | 1 | 0.359 | 0.380346 |
| EPPK1 | MSH6 | 3 | 0.027 | 0.04938 | 1 | 0.035 | 0.093242 |
| ERK2 | INPP4B | 1 | 0.028 | 0.0504 | 1 | 0.015 | 0.048255 |
| ERK2 | ANNEXIN1 | 107 | 0.028 | 0.0504 | 1 | 0.073 | 0.155581 |
| TRANSGLUTAMINASE | ACETYLATUBULINLYS40 | 6 | 0.028 | 0.0504 | 5 | 0.087 | 0.164817 |
| CAVEOLIN1 | EPPK1 | 117 | 0.029 | 0.051656 | 17 | 0.183 | 0.300021 |
| RAD50 | P16INK4A | 1 | 0.029 | 0.051656 | 1 | 0.365 | 0.380346 |
| PAXILLIN | P16INK4A | 6 | 0.03 | 0.052887 | 4 | 0.026 | 0.073883 |
| ECADHERIN | ERALPHA | 375 | 0.03 | 0.052887 | 12 | 0.34 | 0.380346 |
| FIBRONECTIN | PKCPANBETAII_pS660 | 20 | 0.031 | 0.053817 | 9 | 0.001 | 0.007104 |
| GAPDH | TFRC | 84 | 0.031 | 0.053817 | 23 | 0.017 | 0.051759 |
| GAPDH | ANNEXIN1 | 123 | 0.031 | 0.053817 | 16 | 0.2 | 0.300021 |
| ATM | BETACATENIN | 14 | 0.032 | 0.054995 | 1 | 0.075 | 0.15787 |
| ERALPHA | ERK2 | 8 | 0.032 | 0.054995 | 3 | 0.377 | 0.380346 |
| SRC_pY527 | ADAR1 | 1 | 0.033 | 0.055054 | 1 | 0.025 | 0.072246 |
| GSK3_pS9 | PEA15_pS116 | 1 | 0.033 | 0.055054 | 1 | 0.04 | 0.097429 |
| GAPDH | ACETYLATUBULINLYS40 | 28 | 0.033 | 0.055054 | 16 | 0.072 | 0.154415 |
| GATA3 | GAPDH | 60 | 0.033 | 0.055054 | 2 | 0.081 | 0.160587 |
| BETACATENIN | VEGFR2 | 1 | 0.033 | 0.055054 | 1 | 0.194 | 0.300021 |
| ACETYLATUBULINLYS40 | MSH6 | 4 | 0.033 | 0.055054 | 2 | 0.364 | 0.380346 |
| MEK1 | ANNEXIN1 | 62 | 0.034 | 0.056174 | 4 | 0.01 | 0.037889 |
| SYK | PREX1 | 5 | 0.034 | 0.056174 | 1 | 0.043 | 0.103261 |
| S6_pS235S236 | EPPK1 | 6 | 0.035 | 0.057548 | 6 | 0.189 | 0.300021 |
| CAVEOLIN1 | PARPCLEAVED | 4 | 0.036 | 0.058629 | 1 | 0.085 | 0.161927 |
| SRC_pY416 | TFRC | 17 | 0.036 | 0.058629 | 5 | 0.202 | 0.300021 |
| SRC_pY416 | GAPDH | 24 | 0.038 | 0.061302 | 9 | 0.044 | 0.103476 |
| ATM | MEK1 | 4 | 0.038 | 0.061302 | 3 | 0.07 | 0.151076 |
| MYH11 | TFRC | 1050 | 0.04 | 0.064225 | 25 | 0.372 | 0.380346 |
| CAVEOLIN1 | ANNEXIN1 | 2 | 0.051 | 0.081505 | 1 | 0.31 | 0.380346 |
| EPPK1 | PARPCLEAVED | 2 | 0.062 | 0.098623 | 1 | 0.077 | 0.158175 |
| ERALPHA_pS118 | ACETYLATUBULINLYS40 | 1 | 0.064 | 0.100866 | 1 | 0.08 | 0.160471 |
| SYK | MYH11 | 19 | 0.064 | 0.100866 | 6 | 0.203 | 0.300021 |
| PTEN | VEGFR2 | 1 | 0.069 | 0.108248 | 1 | 0.195 | 0.300021 |
| STAT5ALPHA | EPPK1 | 36 | 0.07 | 0.108326 | 3 | 0.047 | 0.109774 |
| X4EBP1_pT37T46 | EEF2 | 3 | 0.07 | 0.108326 | 1 | 0.197 | 0.300021 |
| MYH11 | EPPK1 | 1935 | 0.07 | 0.108326 | 99 | 0.182 | 0.300021 |
| CYCLINB1 | RAB25 | 16 | 0.071 | 0.108402 | 1 | 0.351 | 0.380346 |
| ERALPHA | EPPK1 | 1950 | 0.071 | 0.108402 | 47 | 0.361 | 0.380346 |
| VEGFR2 | ACETYLATUBULINLYS40 | 50 | 0.071 | 0.108402 | 2 | 0.343 | 0.380346 |
| ERK2 | MYH11 | 54 | 0.075 | 0.114 | 6 | 0.198 | 0.300021 |
| PKCPANBETAII_pS660 | RAB25 | 1 | 0.077 | 0.116009 | 1 | 0.04 | 0.097429 |
| CYCLINB1 | EPPK1 | 560 | 0.077 | 0.116009 | 25 | 0.079 | 0.160351 |
| MYH11 | DUSP4 | 16 | 0.081 | 0.1215 | 2 | 0.08 | 0.160471 |
| CAVEOLIN1 | ERALPHA | 130 | 0.085 | 0.126943 | 9 | 0.329 | 0.380346 |
| GAB2 | TFRC | 14 | 0.091 | 0.135313 | 6 | 0.333 | 0.380346 |
| EEF2 | ACETYLATUBULINLYS40 | 18 | 0.092 | 0.136208 | 8 | 0.307 | 0.380346 |
| SYK | ACETYLATUBULINLYS40 | 141 | 0.101 | 0.148249 | 12 | 0.176 | 0.300021 |
| CYCLINE2 | ANNEXIN1 | 1 | 0.101 | 0.148249 | 1 | 0.334 | 0.380346 |
| FASN | CASPASE3 | 1 | 0.102 | 0.149077 | 1 | 0.064 | 0.143579 |
| ETS1 | ANNEXIN1 | 3 | 0.105 | 0.152809 | 1 | 0.347 | 0.380346 |
| CYCLINB1 | ANNEXIN1 | 22 | 0.109 | 0.157958 | 2 | 0.358 | 0.380346 |
| X4EBP1_pT37T46 | ANNEXIN1 | 18 | 0.11 | 0.158734 | 4 | 0.355 | 0.380346 |
| EEF2K | MYH11 | 24 | 0.114 | 0.163815 | 7 | 0.192 | 0.300021 |
| EPPK1 | ACETYLATUBULINLYS40 | 10469 | 0.115 | 0.164561 | 62 | 0.187 | 0.300021 |
| P70S6K_pT389 | YAP_pS127 | 5 | 0.117 | 0.166725 | 1 | 0.001 | 0.007104 |
| SYK | GSK3_pS9 | 2 | 0.12 | 0.168889 | 2 | 0.083 | 0.160813 |
| CAVEOLIN1 | ACETYLATUBULINLYS40 | 5208 | 0.12 | 0.168889 | 22 | 0.377 | 0.380346 |
| EPPK1 | DIRAS3 | 5 | 0.12 | 0.168889 | 3 | 0.353 | 0.380346 |
| RICTOR | ACETYLATUBULINLYS40 | 21 | 0.121 | 0.169598 | 4 | 0.353 | 0.380346 |
| HER2 | SMAC | 1 | 0.123 | 0.171698 | 1 | 0.368 | 0.380346 |
| ERALPHA | ANNEXIN1 | 164 | 0.126 | 0.175171 | 18 | 0.326 | 0.380346 |
| BETACATENIN | GAPDH | 8 | 0.139 | 0.192462 | 6 | 0.214 | 0.309212 |
| ACC1 | GAPDH | 40 | 0.205 | 0.282702 | 4 | 0.008 | 0.031 |
| BETACATENIN | S6_pS240S244 | 1 | 0.209 | 0.28706 | 1 | 0.361 | 0.380346 |
| S6_pS235S236 | ACETYLATUBULINLYS40 | 2 | 0.212 | 0.290016 | 2 | 0.336 | 0.380346 |
| MYH11 | MSH6 | 2 | 0.219 | 0.29622 | 1 | 0.178 | 0.300021 |
| TFRC | ACETYLATUBULINLYS40 | 528 | 0.22 | 0.29622 | 45 | 0.173 | 0.300021 |
| MEK1 | MYH11 | 1 | 0.22 | 0.29622 | 1 | 0.321 | 0.380346 |
| ETS1 | P62LCKLIGAND | 1 | 0.22 | 0.29622 | 1 | 0.337 | 0.380346 |
| BETACATENIN | PEA15_pS116 | 48 | 0.221 | 0.2964 | 6 | 0.014 | 0.046804 |
| CAVEOLIN1 | IGFBP2 | 4 | 0.223 | 0.297914 | 4 | 0.323 | 0.380346 |
| MYH11 | ACETYLATUBULINLYS40 | 2275 | 0.225 | 0.299416 | 24 | 0.179 | 0.300021 |
| BAK | CAVEOLIN1 | 2 | 0.23 | 0.30269 | 2 | 0.34 | 0.380346 |
| BETACATENIN | HSP70 | 21 | 0.231 | 0.30269 | 10 | 0.346 | 0.380346 |
| ERALPHA | SRC_pY527 | 2 | 0.231 | 0.30269 | 1 | 0.368 | 0.380346 |
| P53 | MYH11 | 6 | 0.23 | 0.30269 | 4 | 0.353 | 0.380346 |
| IGFBP2 | EPPK1 | 152 | 0.234 | 0.30545 | 11 | 0.186 | 0.300021 |
| X4EBP1_pT37T46 | MYH11 | 15 | 0.235 | 0.305589 | 5 | 0.183 | 0.300021 |
| INPP4B | PEA15_pS116 | 9 | 0.236 | 0.305727 | 2 | 0.363 | 0.380346 |
| CAVEOLIN1 | RB_pS807S811 | 39 | 0.241 | 0.308697 | 6 | 0.183 | 0.300021 |
| CYCLINB1 | ACETYLATUBULINLYS40 | 959 | 0.24 | 0.308697 | 24 | 0.189 | 0.300021 |
| STAT5ALPHA | YAP_pS127 | 1 | 0.241 | 0.308697 | 1 | 0.344 | 0.380346 |
| EEF2 | ERALPHA | 276 | 0.248 | 0.310014 | 8 | 0.003 | 0.014826 |
| AMPKALPHA_pT172 | CYCLINB1 | 2 | 0.243 | 0.310014 | 1 | 0.082 | 0.160701 |
| ATM | GSK3ALPHABETA_pS21S9 | 1 | 0.249 | 0.310014 | 1 | 0.203 | 0.300021 |
| CHK2 | ACETYLATUBULINLYS40 | 2 | 0.252 | 0.310014 | 1 | 0.199 | 0.300021 |
| ACC1 | EPPK1 | 5 | 0.249 | 0.310014 | 3 | 0.208 | 0.301821 |
| BAK | PDCD4 | 4 | 0.247 | 0.310014 | 3 | 0.362 | 0.380346 |
| CAVEOLIN1 | HSP70 | 5 | 0.252 | 0.310014 | 4 | 0.361 | 0.380346 |
| CYCLINB1 | MYH11 | 850 | 0.247 | 0.310014 | 30 | 0.354 | 0.380346 |
| ERALPHA | GSK3_pS9 | 3 | 0.25 | 0.310014 | 2 | 0.337 | 0.380346 |
| GSK3ALPHABETA_pS21S9 | EPPK1 | 56 | 0.251 | 0.310014 | 6 | 0.359 | 0.380346 |
| EPPK1 | ANNEXIN1 | 76 | 0.244 | 0.310014 | 19 | 0.357 | 0.380346 |
| SYK | ANNEXIN1 | 2 | 0.256 | 0.313806 | 1 | 0.176 | 0.300021 |
| PDCD4 | TFRC | 24 | 0.258 | 0.315129 | 5 | 0.202 | 0.300021 |
| ACETYLATUBULINLYS40 | ADAR1 | 1 | 0.26 | 0.316441 | 1 | 0.205 | 0.300021 |
| BETACATENIN | CYCLINB1 | 162 | 0.364 | 0.398164 | 4 | 0.001 | 0.007104 |
| CYCLINB1 | SRC_pY527 | 22 | 0.365 | 0.398164 | 4 | 0.001 | 0.007104 |
| SYK | GAPDH | 1 | 0.356 | 0.398164 | 1 | 0.002 | 0.011559 |
| ERALPHA | P62LCKLIGAND | 5 | 0.365 | 0.398164 | 5 | 0.003 | 0.014826 |
| ERALPHA | FIBRONECTIN | 4 | 0.364 | 0.398164 | 2 | 0.004 | 0.018432 |
| AKT_pS473 | P16INK4A | 1 | 0.372 | 0.398164 | 1 | 0.015 | 0.048255 |
| ECADHERIN | PEA15_pS116 | 294 | 0.385 | 0.398164 | 7 | 0.016 | 0.049153 |
| AKT_pS473 | ANNEXIN1 | 6 | 0.357 | 0.398164 | 1 | 0.036 | 0.09371 |
| PDCD4 | ANNEXIN1 | 5 | 0.378 | 0.398164 | 4 | 0.083 | 0.160813 |
| CYCLINB1 | ERALPHA | 45 | 0.363 | 0.398164 | 12 | 0.09 | 0.168626 |
| RB_pS807S811 | ANNEXIN1 | 22 | 0.383 | 0.398164 | 3 | 0.162 | 0.297 |
| BETACATENIN | FIBRONECTIN | 32 | 0.362 | 0.398164 | 7 | 0.171 | 0.300021 |
| BETACATENIN | FASN | 18 | 0.371 | 0.398164 | 6 | 0.176 | 0.300021 |
| FASN | ANNEXIN1 | 118 | 0.397 | 0.398164 | 6 | 0.182 | 0.300021 |
| X4EBP1_pT37T46 | FIBRONECTIN | 2 | 0.338 | 0.398164 | 2 | 0.366 | 0.380346 |
| ACC1 | ACETYLATUBULINLYS40 | 3 | 0.371 | 0.398164 | 1 | 0.352 | 0.380346 |
| AKT_pT308 | EPPK1 | 44 | 0.359 | 0.398164 | 7 | 0.344 | 0.380346 |
| BETACATENIN | CAVEOLIN1 | 531 | 0.394 | 0.398164 | 26 | 0.373 | 0.380346 |
| CAVEOLIN1 | NCADHERIN | 5 | 0.383 | 0.398164 | 2 | 0.355 | 0.380346 |
| CAVEOLIN1 | P70S6K_pT389 | 74 | 0.373 | 0.398164 | 1 | 0.354 | 0.380346 |
| CAVEOLIN1 | SRC_pY416 | 32 | 0.383 | 0.398164 | 6 | 0.373 | 0.380346 |
| CLAUDIN7 | GAPDH | 1356 | 0.376 | 0.398164 | 9 | 0.355 | 0.380346 |
| CLAUDIN7 | MYH11 | 50 | 0.395 | 0.398164 | 4 | 0.371 | 0.380346 |
| CLAUDIN7 | EPPK1 | 192 | 0.382 | 0.398164 | 5 | 0.358 | 0.380346 |
| CYCLINB1 | GSK3ALPHABETA_pS21S9 | 8 | 0.353 | 0.398164 | 2 | 0.36 | 0.380346 |
| CYCLINB1 | YAP_pS127 | 8 | 0.384 | 0.398164 | 2 | 0.365 | 0.380346 |
| CYCLINE1 | S6_pS235S236 | 1 | 0.397 | 0.398164 | 1 | 0.373 | 0.380346 |
| ECADHERIN | TFRC | 8 | 0.392 | 0.398164 | 7 | 0.372 | 0.380346 |
| EEF2 | INPP4B | 1 | 0.368 | 0.398164 | 1 | 0.34 | 0.380346 |
| EEF2K | ACETYLATUBULINLYS40 | 4 | 0.358 | 0.398164 | 2 | 0.338 | 0.380346 |
| EGFR_pY1068 | P16INK4A | 1 | 0.379 | 0.398164 | 1 | 0.369 | 0.380346 |
| ERALPHA | P53 | 1 | 0.357 | 0.398164 | 1 | 0.347 | 0.380346 |
| ERALPHA | MYH11 | 741 | 0.361 | 0.398164 | 20 | 0.342 | 0.380346 |
| GAB2 | MYH11 | 15 | 0.388 | 0.398164 | 10 | 0.359 | 0.380346 |
| HER2 | PEA15_pS116 | 8 | 0.378 | 0.398164 | 4 | 0.361 | 0.380346 |
| HSP70 | PEA15_pS116 | 1 | 0.372 | 0.398164 | 1 | 0.341 | 0.380346 |
| HSP70 | ACETYLATUBULINLYS40 | 34 | 0.371 | 0.398164 | 6 | 0.35 | 0.380346 |
| MEK1 | TFRC | 12 | 0.395 | 0.398164 | 2 | 0.375 | 0.380346 |
| NCADHERIN | ANNEXIN1 | 3 | 0.364 | 0.398164 | 1 | 0.349 | 0.380346 |
| PAXILLIN | MYH11 | 30 | 0.375 | 0.398164 | 3 | 0.349 | 0.380346 |
| S6_pS240S244 | ACETYLATUBULINLYS40 | 6 | 0.361 | 0.398164 | 6 | 0.341 | 0.380346 |
| SMAD1 | ANNEXIN1 | 1 | 0.386 | 0.398164 | 1 | 0.342 | 0.380346 |
| SRC_pY416 | P16INK4A | 2 | 0.391 | 0.398164 | 2 | 0.347 | 0.380346 |
| SRC_pY527 | FASN | 1 | 0.356 | 0.398164 | 1 | 0.33 | 0.380346 |
| STAT5ALPHA | ACETYLATUBULINLYS40 | 8 | 0.381 | 0.398164 | 4 | 0.365 | 0.380346 |
| SYK | TFRC | 6 | 0.381 | 0.398164 | 4 | 0.358 | 0.380346 |
| CYCLINE2 | MYH11 | 1 | 0.379 | 0.398164 | 1 | 0.352 | 0.380346 |
| ETS1 | EPPK1 | 105 | 0.366 | 0.398164 | 9 | 0.343 | 0.380346 |
| ETS1 | ACETYLATUBULINLYS40 | 5 | 0.373 | 0.398164 | 5 | 0.36 | 0.380346 |
| FASN | GAPDH | 5600 | 0.39 | 0.398164 | 30 | 0.369 | 0.380346 |
| FASN | ACETYLATUBULINLYS40 | 48 | 0.338 | 0.398164 | 3 | 0.32 | 0.380346 |
| GAPDH | PEA15_pS116 | 30 | 0.375 | 0.398164 | 2 | 0.358 | 0.380346 |
| GAPDH | RAB25 | 3 | 0.387 | 0.398164 | 3 | 0.362 | 0.380346 |
| MYH11 | PKCPANBETAII_pS660 | 12 | 0.367 | 0.398164 | 4 | 0.345 | 0.380346 |
| PDCD4 | RAB25 | 1 | 0.375 | 0.398164 | 1 | 0.349 | 0.380346 |
| PEA15_pS116 | TFRC | 357 | 0.379 | 0.398164 | 4 | 0.359 | 0.380346 |
| PEA15_pS116 | EPPK1 | 4921 | 0.378 | 0.398164 | 17 | 0.358 | 0.380346 |
| PEA15_pS116 | ANNEXIN1 | 504 | 0.385 | 0.398164 | 2 | 0.341 | 0.380346 |
| BETACATENIN | ANNEXIN1 | 220 | 0.387 | 0.398164 | 7 | 0.38 | 0.382242 |
| SRC_pY527 | ANNEXIN1 | 36 | 0.366 | 0.398164 | 3 | 0.396 | 0.396 |
| ERALPHA | DUSP4 | 8 | 0.401 | 0.401 | 5 | 0.347 | 0.380346 |

**Supplementary Table 2.** HPA MAPP list. For each unique protein pair, number of occurrences (count) and p-values were given.

| Protein 1 | Protein 2 | Count | p-value | FDR |
| --- | --- | --- | --- | --- |
| TRPS1 | KRT17 | 2 | 0.005 | 0.036245 |
| TRPS1 | LRRC26 | 2 | 0.004 | 0.036245 |
| TRPS1 | ABHD3 | 2 | 0.002 | 0.036245 |
| PC | ZNF384 | 2 | 0.011 | 0.036245 |
| PRDM1 | KRT17 | 1 | 0.023 | 0.036245 |
| PRDM1 | EID2B | 1 | 0.022 | 0.036245 |
| PRDM1 | ABHD3 | 1 | 0.013 | 0.036245 |
| TRPS1 | EID2B | 1 | 0.019 | 0.036245 |
| ZBTB7B | KRT17 | 1 | 0.018 | 0.036245 |
| ZBTB7B | EID2B | 1 | 0.007 | 0.036245 |
| ZBTB7B | ABHD3 | 1 | 0.01 | 0.036245 |
| SPIDR | KRT5 | 1 | 0.015 | 0.036245 |
| SPIDR | KRT17 | 1 | 0.02 | 0.036245 |
| SPIDR | EID2B | 1 | 0.022 | 0.036245 |
| SPIDR | ABHD3 | 1 | 0.015 | 0.036245 |
| SPIDR | GJC3 | 1 | 0.005 | 0.036245 |
| SPIDR | WT1 | 1 | 0.011 | 0.036245 |
| SPIDR | NFIX | 1 | 0.017 | 0.036245 |
| SPIDR | FZD7 | 1 | 0.014 | 0.036245 |
| AGR3 | EID2B | 1 | 0.014 | 0.036245 |
| AGR3 | LRRC26 | 1 | 0.019 | 0.036245 |
| AGR3 | ABHD3 | 1 | 0.012 | 0.036245 |
| CYBRD1 | LRRC26 | 1 | 0.016 | 0.036245 |
| ATP8B2 | LRRC26 | 1 | 0.019 | 0.036245 |
| KPNA1 | LRRC26 | 1 | 0.005 | 0.036245 |
| HSPB1 | ABHD3 | 1 | 0.009 | 0.036245 |
| HSPB1 | PARD6A | 1 | 0.009 | 0.036245 |
| SLC39A6 | LRRC26 | 1 | 0.007 | 0.036245 |
| SLC39A6 | ABHD3 | 1 | 0.009 | 0.036245 |
| SLC39A6 | PARD6A | 1 | 0.007 | 0.036245 |
| RNASEH2C | CKMT1B | 1 | 0.018 | 0.036245 |
| RNASEH2C | ABHD3 | 1 | 0.015 | 0.036245 |
| RNASEH2C | PARD6A | 1 | 0.016 | 0.036245 |
| SNAPC2 | GJC3 | 1 | 0.007 | 0.036245 |
| SNAPC2 | NFIX | 1 | 0.023 | 0.036245 |
| SNAPC2 | FZD7 | 1 | 0.01 | 0.036245 |
| TAF12 | GJC3 | 1 | 0.021 | 0.036245 |
| TAF12 | NFIX | 1 | 0.021 | 0.036245 |
| TAF12 | FZD7 | 1 | 0.02 | 0.036245 |
| SLC20A2 | GJC3 | 1 | 0.022 | 0.036245 |
| CRIP1 | GJC3 | 1 | 0.021 | 0.036245 |
| CRIP1 | WT1 | 1 | 0.004 | 0.036245 |
| CRIP1 | NES | 1 | 0.006 | 0.036245 |
| CRISPLD1 | ABHD3 | 1 | 0.024 | 0.036245 |
| ENHO | FSCN1 | 1 | 0.014 | 0.036245 |
| ENHO | LAMB2 | 1 | 0.009 | 0.036245 |
| H1FX | FSCN1 | 1 | 0.02 | 0.036245 |
| H1FX | PATJ | 1 | 0.015 | 0.036245 |
| H1FX | YARS | 1 | 0.012 | 0.036245 |
| ARHGAP31 | PATJ | 1 | 0.007 | 0.036245 |
| ARHGAP31 | YARS | 1 | 0.021 | 0.036245 |
| ARHGAP31 | LAMB2 | 1 | 0.019 | 0.036245 |
| GLI3 | FSCN1 | 1 | 0.017 | 0.036245 |
| GLI3 | PATJ | 1 | 0.017 | 0.036245 |
| GLI3 | YARS | 1 | 0.018 | 0.036245 |
| GLI3 | LAMB2 | 1 | 0.005 | 0.036245 |
| PC | YARS | 1 | 0.012 | 0.036245 |
| PC | ISPD | 1 | 0.012 | 0.036245 |
| PC | FAP | 1 | 0.01 | 0.036245 |
| PC | RBM12 | 1 | 0.024 | 0.036245 |
| PC | PM20D2 | 1 | 0.007 | 0.036245 |
| PC | TRAIP | 1 | 0.006 | 0.036245 |
| KRT5 | KIAA1211L | 1 | 0.015 | 0.036245 |
| KRT5 | TMC8 | 1 | 0.023 | 0.036245 |
| HMGB2 | AP4S1 | 1 | 0.016 | 0.036245 |
| HMGB2 | ZNF384 | 1 | 0.018 | 0.036245 |
| WNK2 | AP4S1 | 1 | 0.011 | 0.036245 |
| WNK2 | ZNF384 | 1 | 0.024 | 0.036245 |
| WNK2 | GIGYF1 | 1 | 0.018 | 0.036245 |
| PKP3 | P3H3 | 1 | 0.019 | 0.036245 |
| PKP3 | GSN | 1 | 0.014 | 0.036245 |
| PKP3 | RNF212 | 1 | 0.01 | 0.036245 |
| PKP3 | HADHA | 1 | 0.013 | 0.036245 |
| PKP3 | ISPD | 1 | 0.022 | 0.036245 |
| CKMT1B | PHF14 | 1 | 0.024 | 0.036245 |
| CKMT1B | ISPD | 1 | 0.024 | 0.036245 |
| GFOD2 | FAP | 1 | 0.014 | 0.036245 |
| GFOD2 | RBM12 | 1 | 0.012 | 0.036245 |
| RNGTT | PSMG4 | 1 | 0.006 | 0.036245 |
| RNGTT | ARL17A | 1 | 0.012 | 0.036245 |
| RNGTT | MAN1A2 | 1 | 0.01 | 0.036245 |
| RNGTT | ARL17B | 1 | 0.006 | 0.036245 |
| DDX39B | PSMG4 | 1 | 0.011 | 0.036245 |
| DDX39B | ARL17A | 1 | 0.019 | 0.036245 |
| DDX39B | MAN1A2 | 1 | 0.011 | 0.036245 |
| DDX39B | ARL17B | 1 | 0.024 | 0.036245 |
| FIBP | ACSBG1 | 1 | 0.01 | 0.036245 |
| RAB31 | ACSBG1 | 1 | 0.019 | 0.036245 |
| AK3 | TPD52 | 1 | 0.008 | 0.036245 |
| AK3 | GIGYF1 | 1 | 0.008 | 0.036245 |
| NELFE | TPD52 | 1 | 0.009 | 0.036245 |
| NELFE | GIGYF1 | 1 | 0.009 | 0.036245 |
| PHF14 | SKIL | 1 | 0.012 | 0.036245 |
| PFAS | S100A14 | 1 | 0.016 | 0.036245 |
| KIAA1211L | S100A14 | 1 | 0.015 | 0.036245 |
| LARS | GIGYF1 | 1 | 0.02 | 0.036245 |
| BZW2 | RAF1 | 1 | 0.014 | 0.036245 |
| BZW2 | NAGA | 1 | 0.008 | 0.036245 |
| PRDM1 | LRRC26 | 1 | 0.025 | 0.036634 |
| H1FX | LAMB2 | 1 | 0.025 | 0.036634 |
| FAM81A | GIGYF1 | 1 | 0.025 | 0.036634 |
| ZBTB7B | LRRC26 | 1 | 0.026 | 0.037 |
| HSPB1 | LRRC26 | 1 | 0.026 | 0.037 |
| SIRT2 | GIGYF1 | 1 | 0.026 | 0.037 |
| ZBTB7B | KRT5 | 1 | 0.028 | 0.037649 |
| GAPDH | LRRC26 | 1 | 0.029 | 0.037649 |
| TAF12 | WT1 | 1 | 0.027 | 0.037649 |
| SLC20A2 | WT1 | 1 | 0.028 | 0.037649 |
| SLC20A2 | FZD7 | 1 | 0.029 | 0.037649 |
| DMWD | ABHD3 | 1 | 0.028 | 0.037649 |
| ARHGAP31 | FSCN1 | 1 | 0.029 | 0.037649 |
| PC | AP4S1 | 1 | 0.028 | 0.037649 |
| CKMT1A | PHF14 | 1 | 0.028 | 0.037649 |
| CKMT1A | ISPD | 1 | 0.029 | 0.037649 |
| CRISPLD1 | KRT17 | 1 | 0.03 | 0.037949 |
| PKP3 | SNX2 | 1 | 0.03 | 0.037949 |
| BZW2 | PRDX6 | 1 | 0.03 | 0.037949 |
| FAM81A | TPD52 | 1 | 0.031 | 0.038881 |
| KHDRBS3 | S100A14 | 1 | 0.032 | 0.039798 |
| PRDM1 | KRT5 | 1 | 0.033 | 0.039846 |
| TRPS1 | KRT5 | 1 | 0.034 | 0.039846 |
| RNASEH2C | PKP3 | 1 | 0.035 | 0.039846 |
| RNASEH2C | LRRC26 | 1 | 0.035 | 0.039846 |
| SNAPC2 | WT1 | 1 | 0.033 | 0.039846 |
| SLC20A2 | NFIX | 1 | 0.034 | 0.039846 |
| ENHO | PATJ | 1 | 0.035 | 0.039846 |
| PC | LAMB2 | 1 | 0.034 | 0.039846 |
| KRT5 | PROM1 | 1 | 0.035 | 0.039846 |
| KRT17 | PROM1 | 1 | 0.033 | 0.039846 |
| SUGP2 | S100A14 | 1 | 0.034 | 0.039846 |
| DMWD | LRRC26 | 1 | 0.036 | 0.040364 |
| PKP3 | PHF14 | 1 | 0.036 | 0.040364 |
| PC | FSCN1 | 1 | 0.037 | 0.041173 |
| RNASEH2C | CKMT1A | 1 | 0.038 | 0.04197 |
| SPIDR | LRRC26 | 1 | 0.04 | 0.043529 |
| KRT5 | ALKBH4 | 1 | 0.04 | 0.043529 |
| DMWD | KRT17 | 1 | 0.041 | 0.043971 |
| SIRT2 | TPD52 | 1 | 0.041 | 0.043971 |
| ENHO | YARS | 1 | 0.042 | 0.044719 |
| PKP3 | ST3GAL3 | 1 | 0.043 | 0.045457 |
| PKP3 | MRPL33 | 1 | 0.044 | 0.046184 |
| CRISPLD1 | LRRC26 | 1 | 0.046 | 0.047944 |
| PC | PROM1 | 1 | 0.052 | 0.053818 |
| PKP3 | FITM1 | 1 | 0.053 | 0.054472 |
| AGR3 | KRT5 | 1 | 0.055 | 0.056138 |
| PC | PATJ | 1 | 0.061 | 0.061836 |
| NAP1L1 | ACSBG1 | 1 | 0.063 | 0.063429 |
| AGR3 | KRT17 | 1 | 0.068 | 0.068 |

**Supplementary Table 3.** Differential ratio values for the CS based analysis of the TCPA dataset. Differential expression number, positive and negative correlation numbers, MAPP number and differential ratio values for the PCPP, NCPP, and MAPP were listed for all the proteins with a nonzero differential ratio value. Proteins were ranked from the lowest MAPP based differential ratio value to the highest.

| Protein symbol | Differential expression number | PC number | NC number | MAPP number | Differential ratio for PCPP | Differential ratio for NCPP | Differential ratio for MAPP |
| --- | --- | --- | --- | --- | --- | --- | --- |
| ECADHERIN | 1864 | 1.73837 | 3.532909 | 484 | -0.0467 | -0.02465 | -0.03889 |
| NDRG1_pT346 | 1175 | 1.823323 | 1.064704 | 0 | -0.02634 | -0.02523 | -0.0341 |
| RICTOR | 908 | 1.844018 | 0.735709 | 139 | -0.0185 | -0.02022 | -0.02199 |
| PAI1 | 702 | 1.560319 | 1.200945 | 0 | -0.01373 | -0.01036 | -0.02037 |
| PKCALPHA | 670 | 4.95509 | 1.283458 | 0 | 0.00165 | -0.00875 | -0.01945 |
| PKCALPHA_pS657 | 641 | 5.008151 | 1.253057 | 0 | 0.002717 | -0.00816 | -0.0186 |
| EGFR_pY1068 | 604 | 2.474254 | 0.751512 | 1 | -0.007 | -0.01127 | -0.0175 |
| FASN | 1142 | 1.66178 | 1.636092 | 522 | -0.02607 | -0.01951 | -0.01674 |
| MAPK_pT202Y204 | 519 | 4.202907 | 1.164481 | 0 | 0.00283 | -0.00536 | -0.01506 |
| MYH11 | 2441 | 2.114939 | 1.193788 | 1817 | -0.06184 | -0.0609 | -0.01375 |
| TIGAR | 428 | 1.142539 | 0.881199 | 0 | -0.00756 | -0.00508 | -0.01242 |
| CASPASE7CLEAVEDD198 | 414 | 1.025423 | 0.586865 | 0 | -0.00765 | -0.00712 | -0.01202 |
| AR | 446 | 1.498264 | 0.419172 | 39 | -0.00657 | -0.00945 | -0.01172 |
| CLAUDIN7 | 714 | 1.464793 | 0.719332 | 304 | -0.01449 | -0.01473 | -0.01117 |
| GATA3 | 464 | 1.316073 | 0.204726 | 91 | -0.00786 | -0.01176 | -0.01061 |
| PREX1 | 400 | 3.590087 | 0.687821 | 79 | 0.003675 | -0.00588 | -0.00913 |
| CKIT | 312 | 1.370841 | 0.648687 | 0 | -0.00322 | -0.00365 | -0.00906 |
| HER2_pY1248 | 252 | 2.085031 | 0.295454 | 0 | 0.001563 | -0.00485 | -0.00731 |
| PEA15 | 215 | 3.532408 | 0.936392 | 0 | 0.008799 | 0.001566 | -0.00624 |
| RBM15 | 201 | 2.39613 | 1.592523 | 0 | 0.004367 | 0.007442 | -0.00583 |
| BRD4 | 178 | 1.236056 | 2.007377 | 0 | 9.61E-05 | 0.011568 | -0.00517 |
| IGFBP2 | 401 | 1.211837 | 0.524848 | 208 | -0.00648 | -0.00726 | -0.0051 |
| EPPK1 | 2743 | 2.061131 | 2.397394 | 2372 | -0.07084 | -0.05963 | -0.00507 |
| EGFR | 165 | 2.325907 | 0.354593 | 0 | 0.005113 | -0.00183 | -0.00479 |
| PKCDELTA_pS664 | 158 | 3.494346 | 0.816316 | 0 | 0.010291 | 0.002219 | -0.00459 |
| INPP4B | 392 | 1.307187 | 0.303436 | 229 | -0.00581 | -0.00885 | -0.00418 |
| EGFR_pY1173 | 144 | 2.657196 | 0.971312 | 0 | 0.007133 | 0.003918 | -0.00418 |
| COLLAGENVI | 131 | 0.830305 | 0.691594 | 0 | -0.00027 | 0.001963 | -0.0038 |
| RAD51 | 124 | 1.324852 | 1.6059 | 0 | 0.002041 | 0.009789 | -0.0036 |
| S6 | 119 | 1.86044 | 1.60648 | 0 | 0.004467 | 0.009938 | -0.00345 |
| MYOSINIIA_pS1943 | 105 | 0.956068 | 2.17461 | 0 | 0.001023 | 0.015081 | -0.00305 |
| AKT_pS473 | 259 | 2.023367 | 0.297175 | 144 | 0.001097 | -0.00504 | -0.00299 |
| XBP1 | 99 | 1.325615 | 0.788162 | 0 | 0.00277 | 0.003697 | -0.00287 |
| G6PD | 96 | 0.518465 | 0.396476 | 0 | -0.00058 | 0.000519 | -0.00279 |
| X53BP1 | 108 | 2.211928 | 1.274188 | 18 | 0.006282 | 0.007488 | -0.00257 |
| ASNS | 74 | 0.815026 | 0.438866 | 0 | 0.001322 | 0.001511 | -0.00215 |
| SMAC | 62 | 0.817877 | 0.710409 | 1 | 0.001683 | 0.004123 | -0.00177 |
| NFKBP65_pS536 | 56 | 0.437016 | 0.140354 | 0 | 0.000235 | -0.00046 | -0.00163 |
| CD26 | 54 | 0.354636 | 0.174262 | 0 | -5.75E-05 | -0.00011 | -0.00157 |
| AKT | 51 | 2.738966 | 0.777634 | 0 | 0.010181 | 0.005002 | -0.00148 |
| ACC_pS79 | 50 | 0.921607 | 0.502206 | 2 | 0.002472 | 0.002735 | -0.00139 |
| EIF4G | 47 | 1.344085 | 0.668523 | 0 | 0.004358 | 0.004209 | -0.00136 |
| HER3 | 41 | 0.709197 | 0.603598 | 0 | 0.001829 | 0.003842 | -0.00119 |
| KU80 | 40 | 1.669544 | 1.435741 | 0 | 0.005947 | 0.010808 | -0.00116 |
| BCL2 | 38 | 0.470201 | 0.151625 | 0 | 0.000899 | 0.000161 | -0.0011 |
| SHP2_pY542 | 38 | 1.233855 | 0.1811 | 0 | 0.00415 | 0.000407 | -0.0011 |
| PDL1 | 30 | 0.96125 | 0.838908 | 0 | 0.003222 | 0.006123 | -0.00087 |
| TUBERIN | 29 | 1.646469 | 0.387563 | 0 | 0.006168 | 0.002389 | -0.00084 |
| P70S6K1 | 28 | 1.916994 | 0.860682 | 0 | 0.007349 | 0.006362 | -0.00081 |
| RAB11 | 28 | 0.390398 | 0.192383 | 0 | 0.000849 | 0.000791 | -0.00081 |
| ERCC1 | 28 | 0.493636 | 0.077406 | 0 | 0.001289 | -0.00017 | -0.00081 |
| MEK1_pS217S221 | 25 | 2.04425 | 1.440543 | 0 | 0.007978 | 0.011283 | -0.00073 |
| ERALPHA_pS118 | 26 | 0.409628 | 0.554654 | 1 | 0.000989 | 0.003869 | -0.00072 |
| CMYC | 22 | 0.684622 | 0.43601 | 0 | 0.002276 | 0.002996 | -0.00064 |
| YAP | 22 | 0.641673 | 0.06073 | 0 | 0.002093 | -0.00013 | -0.00064 |
| CD20 | 20 | 0.708742 | 0.099453 | 0 | 0.002437 | 0.000249 | -0.00058 |
| X4EBP1 | 19 | 0.902811 | 0.326318 | 0 | 0.003292 | 0.002169 | -0.00055 |
| SETD2 | 19 | 0.371736 | 0.164115 | 0 | 0.001031 | 0.000817 | -0.00055 |
| LCK | 18 | 0.253452 | 0.038661 | 0 | 0.000557 | -0.0002 | -0.00052 |
| BAP1C4 | 18 | 0.572998 | 0.257205 | 0 | 0.001917 | 0.001622 | -0.00052 |
| BIM | 15 | 0.456463 | 0.069881 | 0 | 0.001508 | 0.000147 | -0.00044 |
| CHK2_pT68 | 15 | 2.198005 | 0.912773 | 0 | 0.008922 | 0.007174 | -0.00044 |
| P27 | 15 | 0.459338 | 0.128239 | 0 | 0.00152 | 0.000634 | -0.00044 |
| HEREGULIN | 15 | 1.036746 | 0.51401 | 0 | 0.003978 | 0.00385 | -0.00044 |
| SCD1 | 14 | 1.245544 | 0.19713 | 0 | 0.004896 | 0.001237 | -0.00041 |
| CD49B | 13 | 0.284465 | 0.216291 | 0 | 0.000834 | 0.001426 | -0.00038 |
| MIG6 | 13 | 0.22817 | 0.055896 | 0 | 0.000594 | 8.87E-05 | -0.00038 |
| PRAS40_pT246 | 13 | 0.373469 | 0.039372 | 0 | 0.001213 | -4.91E-05 | -0.00038 |
| SRC | 13 | 0.232006 | 0.141437 | 0 | 0.00061 | 0.000802 | -0.00038 |
| AMPKALPHA | 11 | 0.105089 | 0.024501 | 0 | 0.000128 | -0.00012 | -0.00032 |
| CYCLINE1 | 54 | 0.322398 | 0.175993 | 40 | -0.00019 | -0.0001 | -0.00031 |
| LKB1 | 10 | 0.539049 | 0.770789 | 0 | 0.002005 | 0.006135 | -0.00029 |
| SMAD3 | 10 | 0.414038 | 0.204946 | 0 | 0.001472 | 0.001418 | -0.00029 |
| SNAIL | 10 | 1.152065 | 0.082259 | 0 | 0.004615 | 0.000396 | -0.00029 |
| JNK_pT183Y185 | 9 | 0.559602 | 0.082919 | 0 | 0.002121 | 0.00043 | -0.00026 |
| STAT3_pY705 | 9 | 0.570052 | 0.265923 | 0 | 0.002166 | 0.001956 | -0.00026 |
| YB1 | 9 | 0.496793 | 0.00625 | 0 | 0.001854 | -0.00021 | -0.00026 |
| BRCA2 | 9 | 0.729663 | 0.094082 | 0 | 0.002845 | 0.000523 | -0.00026 |
| CDK1 | 8 | 0.307686 | 0.392395 | 0 | 0.001078 | 0.003039 | -0.00023 |
| GAB2 | 117 | 1.225601 | 0.303651 | 101 | 0.001822 | -0.00086 | -0.00022 |
| CIAP | 7 | 0.451782 | 0.020646 | 0 | 0.00172 | -3.11E-05 | -0.0002 |
| MRE11 | 7 | 2.530714 | 1.193143 | 0 | 0.010571 | 0.009743 | -0.0002 |
| BCLXL | 6 | 0.050127 | 0.052971 | 0 | 3.93E-05 | 0.000267 | -0.00017 |
| CRAF_pS338 | 6 | 1.233597 | 1.272719 | 0 | 0.005078 | 0.010436 | -0.00017 |
| SMAD1 | 7 | 0.150798 | 0.087531 | 1 | 0.000439 | 0.000527 | -0.00017 |
| NOTCH1 | 5 | 0.096993 | 0.02283 | 0 | 0.000268 | 4.52E-05 | -0.00015 |
| FOXM1 | 5 | 0.071253 | 0.023478 | 0 | 0.000158 | 5.06E-05 | -0.00015 |
| SF2 | 5 | 0.715875 | 0.347558 | 0 | 0.002903 | 0.002752 | -0.00015 |
| TAZ | 5 | 0.985398 | 0.171424 | 0 | 0.00405 | 0.001284 | -0.00015 |
| CMET | 5 | 0.965209 | 0.05167 | 0 | 0.003964 | 0.000286 | -0.00015 |
| CYCLINE2 | 9 | 0.800794 | 0.480099 | 4 | 0.003148 | 0.003741 | -0.00014 |
| CD31 | 4 | 1.567727 | 0.211353 | 0 | 0.006558 | 0.001646 | -0.00012 |
| FOXO3A | 4 | 1.397496 | 0.097737 | 0 | 0.005834 | 0.000699 | -0.00012 |
| MTOR | 4 | 2.090723 | 0.885828 | 0 | 0.008785 | 0.007269 | -0.00012 |
| NF2 | 4 | 0.022481 | 0.025695 | 0 | -2.04E-05 | 9.81E-05 | -0.00012 |
| PDK1_pS241 | 4 | 1.831858 | 0.82731 | 0 | 0.007683 | 0.006781 | -0.00012 |
| CABL | 4 | 0.221621 | 0.120737 | 0 | 0.000827 | 0.00089 | -0.00012 |
| ERCC5 | 4 | 0.220983 | 0.138472 | 0 | 0.000825 | 0.001038 | -0.00012 |
| CASPASE3 | 5 | 0.901656 | 0.157434 | 1 | 0.003694 | 0.001167 | -0.00011 |
| ADAR1 | 6 | 0.404681 | 0.535445 | 2 | 0.001549 | 0.00429 | -0.00011 |
| CHK1 | 3 | 0.213711 | 0.107618 | 0 | 0.000823 | 0.00081 | -8.71E-05 |
| P27_pT157 | 3 | 0.823746 | 0.094897 | 0 | 0.00342 | 0.000704 | -8.71E-05 |
| PDK1 | 3 | 0.276415 | 0.156515 | 0 | 0.00109 | 0.001218 | -8.71E-05 |
| PRDX1 | 3 | 0.440282 | 0.204216 | 0 | 0.001787 | 0.001615 | -8.71E-05 |
| RICTOR_pT1135 | 3 | 2.126486 | 0.469172 | 0 | 0.008966 | 0.003824 | -8.71E-05 |
| X1433BETA | 3 | 0.623797 | 0.008992 | 0 | 0.002569 | -1.21E-05 | -8.71E-05 |
| X4EBP1_pS65 | 2 | 0.066632 | 0 | 0 | 0.000226 | -5.80E-05 | -5.80E-05 |
| CMET_pY1235 | 2 | 0.34956 | 0.355431 | 0 | 0.00143 | 0.002905 | -5.80E-05 |
| DJ1 | 2 | 0.003074 | 0.081732 | 0 | -4.50E-05 | 0.000623 | -5.80E-05 |
| DVL3 | 2 | 0.071374 | 0.115571 | 0 | 0.000246 | 0.000905 | -5.80E-05 |
| GSK3ALPHABETA | 2 | 1.05245 | 2.20955 | 0 | 0.004423 | 0.018362 | -5.80E-05 |
| IRS1 | 2 | 0.758019 | 0.174209 | 0 | 0.003169 | 0.001394 | -5.80E-05 |
| JNK2 | 2 | 0.022278 | 0.003108 | 0 | 3.68E-05 | -3.21E-05 | -5.80E-05 |
| P38MAPK | 2 | 1.058 | 0.749372 | 0 | 0.004446 | 0.006189 | -5.80E-05 |
| P90RSK_pT359S363 | 2 | 0.651335 | 0.063429 | 0 | 0.002715 | 0.000471 | -5.80E-05 |
| STATHMIN | 2 | 1.668902 | 0.479574 | 0 | 0.007047 | 0.00394 | -5.80E-05 |
| YB1_pS102 | 2 | 0.02226 | 0.001265 | 0 | 3.67E-05 | -4.75E-05 | -5.80E-05 |
| PI3KP85 | 2 | 0.029523 | 0.674674 | 0 | 6.76E-05 | 0.005566 | -5.80E-05 |
| RAPTOR | 2 | 0.00085 | 0.0276 | 0 | -5.44E-05 | 0.000172 | -5.80E-05 |
| TSC1 | 2 | 2.185511 | 1.159071 | 0 | 0.009247 | 0.009605 | -5.80E-05 |
| TUBERIN_pT1462 | 2 | 1.09089 | 0.0392 | 0 | 0.004586 | 0.000269 | -5.80E-05 |
| X1433ZETA | 2 | 0.482191 | 0.379334 | 0 | 0.001995 | 0.003104 | -5.80E-05 |
| ACVRL1 | 2 | 0.307922 | 0.0154 | 0 | 0.001253 | 7.03E-05 | -5.80E-05 |
| BRAF_pS445 | 2 | 0.025621 | 0.05804 | 0 | 5.10E-05 | 0.000426 | -5.80E-05 |
| COG3 | 2 | 0.108633 | 0.007968 | 0 | 0.000404 | 8.38E-06 | -5.80E-05 |
| JAK2 | 2 | 0.021069 | 0.023671 | 0 | 3.16E-05 | 0.000139 | -5.80E-05 |
| HER3_pY1289 | 3 | 0.267294 | 0.063126 | 1 | 0.001051 | 0.000439 | -5.56E-05 |
| BAX | 1 | 0.246633 | 1.819427 | 0 | 0.001021 | 0.015139 | -2.90E-05 |
| CRAF | 1 | 0.000365 | 0 | 0 | -2.75E-05 | -2.90E-05 | -2.90E-05 |
| CYCLIND1 | 1 | 0.753523 | 0.159125 | 0 | 0.003179 | 0.001298 | -2.90E-05 |
| MTOR_pS2448 | 1 | 0.027044 | 0.052382 | 0 | 8.61E-05 | 0.000408 | -2.90E-05 |
| PI3KP110ALPHA | 1 | 0.000365 | 0 | 0 | -2.75E-05 | -2.90E-05 | -2.90E-05 |
| PR | 1 | 0.001964 | 0.258536 | 0 | -2.07E-05 | 0.002126 | -2.90E-05 |
| SHC_pY317 | 1 | 0.125398 | 0.006858 | 0 | 0.000505 | 2.81E-05 | -2.90E-05 |
| SMAD4 | 1 | 0.022157 | 0.003404 | 0 | 6.53E-05 | -6.50E-07 | -2.90E-05 |
| FOXO3A_pS318S321 | 1 | 0.136201 | 0.08354 | 0 | 0.000551 | 0.000667 | -2.90E-05 |
| NRAS | 1 | 0.853252 | 0.056844 | 0 | 0.003604 | 0.000445 | -2.90E-05 |
| P21 | 1 | 0 | 0.000901 | 0 | -2.90E-05 | -2.15E-05 | -2.90E-05 |
| CASPASE8 | 1 | 1.173361 | 0.038302 | 0 | 0.004966 | 0.00029 | -2.90E-05 |
| RB | 1 | 1.506514 | 0.610776 | 0 | 0.006385 | 0.005063 | -2.90E-05 |
| BCL2A1 | 1 | 0.095264 | 0.015273 | 0 | 0.000377 | 9.83E-05 | -2.90E-05 |
| IRF1 | 1 | 0.010317 | 0 | 0 | 1.49E-05 | -2.90E-05 | -2.90E-05 |
| MSH2 | 27 | 1.168131 | 0.158277 | 39 | 0.00419 | 0.000536 | 0.000442 |
| TFRC | 1171 | 1.313455 | 2.084415 | 1098 | -0.0284 | -0.01661 | 0.000517 |
| TRANSGLUTAMINASE | 180 | 0.886979 | 0.392298 | 188 | -0.00145 | -0.00195 | 0.000684 |
| CHK2 | 15 | 0.886733 | 0.494018 | 40 | 0.00334 | 0.003683 | 0.000822 |
| CDK1_pY15 | 9 | 0.387055 | 0.088741 | 39 | 0.001387 | 0.000479 | 0.000964 |
| RAD50 | 69 | 0.944891 | 0.450091 | 95 | 0.00202 | 0.001749 | 0.000983 |
| AMPKALPHA_pT172 | 76 | 1.007711 | 0.808924 | 110 | 0.002084 | 0.004538 | 0.001251 |
| P62LCKLIGAND | 97 | 0.704923 | 0.305638 | 133 | 0.000186 | -0.00027 | 0.001364 |
| P70S6K_pT389 | 231 | 3.849433 | 1.025614 | 257 | 0.009684 | 0.001845 | 0.001372 |
| VEGFR2 | 102 | 1.190431 | 1.167534 | 147 | 0.002108 | 0.006773 | 0.001659 |
| P53 | 103 | 0.348536 | 0.259322 | 149 | -0.00151 | -0.00083 | 0.001693 |
| AKT_pT308 | 132 | 1.177332 | 0.149552 | 179 | 0.001181 | -0.00258 | 0.001794 |
| PTEN | 16 | 1.129141 | 0.730761 | 72 | 0.004343 | 0.005628 | 0.001798 |
| JAB1 | 7 | 2.285078 | 0.151038 | 70 | 0.009525 | 0.001056 | 0.001997 |
| EEF2K | 54 | 1.439914 | 0.609644 | 119 | 0.004563 | 0.003515 | 0.002172 |
| DIRAS3 | 9 | 2.596217 | 0.36791 | 79 | 0.010792 | 0.002806 | 0.002221 |
| PKCPANBETAII_pS660 | 248 | 4.599621 | 1.12932 | 304 | 0.012384 | 0.002217 | 0.002355 |
| HSP70 | 252 | 1.140606 | 1.005559 | 315 | -0.00246 | 0.001069 | 0.002585 |
| S6_pS240S244 | 111 | 1.499646 | 0.663156 | 193 | 0.003163 | 0.002307 | 0.002843 |
| PARPCLEAVED | 16 | 2.022928 | 0.302433 | 110 | 0.008148 | 0.002057 | 0.002992 |
| S6_pS235S236 | 30 | 0.950954 | 0.420434 | 131 | 0.003178 | 0.002634 | 0.003246 |
| GAPDH | 1318 | 1.137142 | 1.530588 | 1321 | -0.03341 | -0.02549 | 0.003258 |
| EEF2 | 141 | 1.428061 | 1.068573 | 237 | 0.001987 | 0.004816 | 0.003355 |
| NCADHERIN | 33 | 2.029757 | 1.52656 | 140 | 0.007684 | 0.011768 | 0.003442 |
| MSH6 | 41 | 1.251447 | 0.236246 | 167 | 0.004138 | 0.000779 | 0.004058 |
| P38_pT180Y182 | 24 | 0.986961 | 0.532845 | 152 | 0.003505 | 0.003745 | 0.00408 |
| DUSP4 | 148 | 0.498775 | 0.192327 | 270 | -0.00217 | -0.00269 | 0.004189 |
| BAK | 118 | 1.190693 | 1.299494 | 250 | 0.001644 | 0.007408 | 0.004431 |
| ERK2 | 381 | 4.960262 | 1.325688 | 496 | 0.01006 | -6.68E-06 | 0.004528 |
| HER2 | 163 | 1.452897 | 0.242938 | 330 | 0.001455 | -0.00271 | 0.005639 |
| BRAF | 120 | 2.82181 | 0.94543 | 300 | 0.008531 | 0.004399 | 0.005945 |
| RB_pS807S811 | 96 | 1.666244 | 0.593182 | 283 | 0.004308 | 0.002159 | 0.006107 |
| GSK3_pS9 | 66 | 1.97009 | 0.7434 | 278 | 0.006472 | 0.004282 | 0.00682 |
| STAT5ALPHA | 137 | 1.92575 | 0.616116 | 348 | 0.004222 | 0.00116 | 0.00696 |
| RAB25 | 102 | 0.649719 | 0.739581 | 332 | -0.00019 | 0.003205 | 0.007473 |
| CYCLINB1 | 887 | 1.202805 | 2.841539 | 1059 | -0.02062 | -0.00206 | 0.007534 |
| ETS1 | 108 | 1.487716 | 0.710613 | 345 | 0.003199 | 0.002789 | 0.007707 |
| ACC1 | 195 | 1.084448 | 1.135635 | 445 | -0.00104 | 0.003807 | 0.008324 |
| PAXILLIN | 98 | 1.458367 | 0.837141 | 396 | 0.003364 | 0.004134 | 0.0096 |
| SRC_pY416 | 203 | 1.967572 | 0.755539 | 493 | 0.002485 | 0.000407 | 0.009601 |
| PDCD4 | 107 | 1.437957 | 1.186876 | 413 | 0.003016 | 0.006789 | 0.009873 |
| MEK1 | 75 | 1.728789 | 0.794764 | 388 | 0.005183 | 0.004449 | 0.010016 |
| FIBRONECTIN | 242 | 0.894297 | 0.88612 | 556 | -0.00322 | 0.000363 | 0.010448 |
| P16INK4A | 143 | 0.850245 | 1.044996 | 474 | -0.00053 | 0.004561 | 0.010745 |
| BETACATENIN | 445 | 2.561703 | 1.316244 | 783 | -0.00201 | -0.00194 | 0.01169 |
| SRC_pY527 | 83 | 1.351104 | 0.558484 | 455 | 0.003343 | 0.002247 | 0.011889 |
| YAP_pS127 | 101 | 1.16794 | 0.839313 | 480 | 0.002041 | 0.004065 | 0.012152 |
| X4EBP1_pT37T46 | 76 | 1.412508 | 0.594907 | 478 | 0.003808 | 0.002754 | 0.012815 |
| SYK | 204 | 1.455084 | 1.584287 | 597 | 0.000274 | 0.007286 | 0.01284 |
| GSK3ALPHABETA_pS21S9 | 52 | 1.311118 | 0.522894 | 469 | 0.004073 | 0.00285 | 0.013229 |
| ATM | 93 | 1.863475 | 1.558364 | 534 | 0.005234 | 0.010292 | 0.014082 |
| ERALPHA | 1192 | 1.144997 | 1.217854 | 1585 | -0.02972 | -0.02444 | 0.015211 |
| CAVEOLIN1 | 1159 | 2.038652 | 1.674761 | 1640 | -0.02496 | -0.01968 | 0.017898 |
| PEA15_pS116 | 559 | 2.178364 | 1.098564 | 1353 | -0.00695 | -0.00707 | 0.026293 |
| ANNEXIN1 | 529 | 1.506359 | 1.996937 | 1589 | -0.00894 | 0.001294 | 0.03458 |
| ACETYLATUBULINLYS40 | 879 | 4.274116 | 1.692934 | 1918 | -0.00732 | -0.0114 | 0.03476 |

**Supplementary Table 4.** Differential ratio values for the CT based analysis of the TCPA dataset. Differential expression number, positive and negative correlation numbers, MAPP number and differential ratio values for the PCPP, NCPP, and MAPP were listed for all the proteins with a nonzero differential ratio value. Proteins were ranked from the lowest MAPP based differential ratio value to the highest.

| Protein symbol | Differential expression number | PC number | NC number | MAPP number | Differential ratio for PCPP | Differential ratio for NCPP | Differential ratio for MAPP |
| --- | --- | --- | --- | --- | --- | --- | --- |
| PAI1 | 29 | 36.77177 | 24.2237 | 0 | -0.00297 | -0.00467 | -0.01245 |
| NDRG1_pT346 | 25 | 38.15393 | 22.3921 | 0 | -0.0009 | -0.00354 | -0.01073 |
| RBM15 | 24 | 22.22243 | 38.99293 | 0 | -0.00457 | 0.002217 | -0.0103 |
| CASPASE7CLEAVEDD198 | 22 | 34.80225 | 21.10492 | 0 | -0.00047 | -0.00267 | -0.00944 |
| AMPKALPHA_pT172 | 22 | 20.50052 | 30.0337 | 3 | -0.00416 | 0.000199 | -0.00913 |
| ASNS | 21 | 33.83468 | 20.95471 | 0 | -0.0003 | -0.00229 | -0.00901 |
| CKIT | 21 | 33.67491 | 18.95558 | 0 | -0.00034 | -0.00293 | -0.00901 |
| MAPK_pT202Y204 | 20 | 35.9757 | 19.89297 | 0 | 0.000686 | -0.0022 | -0.00858 |
| BRD4 | 20 | 22.28978 | 34.80755 | 0 | -0.00284 | 0.00259 | -0.00858 |
| NFKBP65_pS536 | 19 | 32.23513 | 20.35444 | 0 | 0.000151 | -0.00162 | -0.00815 |
| EGFR_pY1068 | 19 | 36.96146 | 21.1369 | 1 | 0.001369 | -0.00137 | -0.00805 |
| AKT | 17 | 21.39772 | 28.74993 | 0 | -0.00178 | 0.001933 | -0.0073 |
| S6 | 17 | 20.88998 | 36.91443 | 0 | -0.00191 | 0.004554 | -0.0073 |
| HER3 | 16 | 29.23676 | 18.61975 | 0 | 0.000666 | -0.00089 | -0.00687 |
| SMAC | 16 | 23.22597 | 19.81228 | 1 | -0.00088 | -0.00051 | -0.00676 |
| RICTOR | 27 | 37.65032 | 26.08836 | 47 | -0.00189 | -0.00321 | -0.00669 |
| HER2_pY1248 | 15 | 36.01367 | 17.52238 | 0 | 0.002842 | -0.00081 | -0.00644 |
| MEK1_pS217S221 | 15 | 30.64894 | 20.36217 | 0 | 0.001459 | 9.90E-05 | -0.00644 |
| PKCALPHA | 15 | 29.47736 | 16.52911 | 0 | 0.001157 | -0.00113 | -0.00644 |
| PKCALPHA_pS657 | 15 | 29.88885 | 15.23765 | 0 | 0.001263 | -0.00155 | -0.00644 |
| G6PD | 15 | 30.52454 | 16.83104 | 0 | 0.001427 | -0.00103 | -0.00644 |
| PDL1 | 15 | 32.57349 | 19.28524 | 0 | 0.001955 | -0.00025 | -0.00644 |
| ACC_pS79 | 15 | 25.51144 | 20.82153 | 2 | 0.000136 | 0.000246 | -0.00623 |
| KU80 | 14 | 16.91287 | 25.8527 | 0 | -0.00165 | 0.002291 | -0.00601 |
| EIF4G | 14 | 16.60502 | 35.658 | 0 | -0.00173 | 0.005438 | -0.00601 |
| MYOSINIIA_pS1943 | 14 | 18.45037 | 28.2569 | 0 | -0.00125 | 0.003063 | -0.00601 |
| TUBERIN | 13 | 14.95764 | 24.69931 | 0 | -0.00173 | 0.00235 | -0.00558 |
| COLLAGENVI | 12 | 26.65296 | 15.31451 | 0 | 0.001717 | -0.00023 | -0.00515 |
| EGFR | 12 | 33.37362 | 16.66874 | 0 | 0.003449 | 0.000201 | -0.00515 |
| EGFR_pY1173 | 12 | 30.62838 | 17.14261 | 0 | 0.002742 | 0.000353 | -0.00515 |
| XBP1 | 12 | 23.83625 | 15.85638 | 0 | 0.000991 | -6.00E-05 | -0.00515 |
| AKT_pS473 | 21 | 34.88162 | 22.42322 | 39 | -2.53E-05 | -0.00181 | -0.00495 |
| P70S6K1 | 11 | 18.18695 | 22.93895 | 0 | -3.50E-05 | 0.002643 | -0.00472 |
| RAD51 | 11 | 21.53252 | 15.04458 | 0 | 0.000827 | 0.000109 | -0.00472 |
| X53BP1 | 24 | 22.56313 | 38.09508 | 55 | -0.00449 | 0.001929 | -0.00457 |
| CYCLINE1 | 11 | 25.13532 | 14.76348 | 2 | 0.001755 | 1.84E-05 | -0.00451 |
| P70S6K_pT389 | 11 | 22.64861 | 15.61933 | 3 | 0.001115 | 0.000293 | -0.00441 |
| HEREGULIN | 10 | 23.29564 | 15.44955 | 0 | 0.001711 | 0.000668 | -0.00429 |
| S6_pS240S244 | 26 | 32.79831 | 26.95772 | 66 | -0.00271 | -0.0025 | -0.00428 |
| VEGFR2 | 16 | 28.46662 | 21.07481 | 26 | 0.000468 | -0.0001 | -0.00416 |
| AR | 12 | 23.65739 | 16.98862 | 12 | 0.000945 | 0.000304 | -0.0039 |
| CMYC | 9 | 23.64606 | 10.60694 | 0 | 0.00223 | -0.00046 | -0.00386 |
| SCD1 | 9 | 19.97156 | 11.27701 | 0 | 0.001283 | -0.00024 | -0.00386 |
| TIGAR | 9 | 21.44728 | 12.52762 | 0 | 0.001663 | 0.000159 | -0.00386 |
| PREX1 | 9 | 22.33922 | 14.73445 | 1 | 0.001893 | 0.000867 | -0.00376 |
| BIM | 8 | 25.32752 | 11.70403 | 0 | 0.003092 | 0.000324 | -0.00343 |
| CHK2_pT68 | 8 | 20.39415 | 13.51508 | 0 | 0.001821 | 0.000905 | -0.00343 |
| PARPCLEAVED | 8 | 20.77025 | 14.40039 | 3 | 0.001918 | 0.001189 | -0.00312 |
| INPP4B | 26 | 36.62747 | 27.27038 | 78 | -0.00172 | -0.0024 | -0.00303 |
| X4EBP1 | 7 | 24.64602 | 14.54118 | 0 | 0.003346 | 0.001664 | -0.003 |
| BCL2 | 7 | 24.37871 | 9.781849 | 0 | 0.003277 | 0.000136 | -0.003 |
| CD49B | 7 | 22.9017 | 10.46687 | 0 | 0.002897 | 0.000356 | -0.003 |
| LCK | 7 | 15.18729 | 6.357052 | 0 | 0.000909 | -0.00096 | -0.003 |
| SMAD3 | 7 | 23.33995 | 10.39149 | 0 | 0.003009 | 0.000332 | -0.003 |
| SETD2 | 7 | 22.04851 | 9.737643 | 0 | 0.002677 | 0.000122 | -0.003 |
| RAD50 | 13 | 25.36191 | 19.0246 | 25 | 0.000955 | 0.000528 | -0.00297 |
| TRANSGLUTAMINASE | 19 | 34.40838 | 20.58199 | 50 | 0.000711 | -0.00155 | -0.00294 |
| STAT5ALPHA | 24 | 25.79941 | 32.21582 | 72 | -0.00365 | 4.16E-05 | -0.0028 |
| PTEN | 12 | 20.16097 | 27.85802 | 23 | 4.45E-05 | 0.003793 | -0.00275 |
| EEF2K | 17 | 21.98517 | 26.85744 | 44 | -0.00163 | 0.001326 | -0.00271 |
| STAT3_pY705 | 6 | 20.8662 | 9.778732 | 0 | 0.002801 | 0.000564 | -0.00258 |
| ERCC1 | 6 | 19.81528 | 7.350386 | 0 | 0.002531 | -0.00022 | -0.00258 |
| SNAIL | 6 | 18.14729 | 11.07325 | 0 | 0.002101 | 0.00098 | -0.00258 |
| PDCD4 | 27 | 30.23036 | 28.69539 | 88 | -0.0038 | -0.00238 | -0.00242 |
| P62LCKLIGAND | 23 | 34.96529 | 23.12914 | 73 | -0.00086 | -0.00245 | -0.00226 |
| CYCLINE2 | 6 | 15.66003 | 9.656955 | 4 | 0.00146 | 0.000525 | -0.00216 |
| CIAP | 5 | 16.7657 | 9.36682 | 0 | 0.002174 | 0.000861 | -0.00215 |
| NOTCH1 | 5 | 15.73295 | 8.741988 | 0 | 0.001908 | 0.00066 | -0.00215 |
| SRC | 5 | 15.10421 | 12.10067 | 0 | 0.001746 | 0.001739 | -0.00215 |
| YAP | 5 | 20.0976 | 11.09587 | 0 | 0.003032 | 0.001416 | -0.00215 |
| BAP1C4 | 5 | 7.101492 | 10.6891 | 0 | -0.00032 | 0.001286 | -0.00215 |
| ERALPHA_pS118 | 5 | 16.04304 | 11.08198 | 1 | 0.001988 | 0.001412 | -0.00204 |
| S6_pS235S236 | 15 | 24.52539 | 22.73015 | 43 | -0.00012 | 0.000859 | -0.00196 |
| CHK2 | 5 | 9.331334 | 17.13129 | 2 | 0.000258 | 0.003354 | -0.00194 |
| EEF2 | 27 | 31.00168 | 32.31834 | 94 | -0.0036 | -0.00121 | -0.00179 |
| IGFBP2 | 27 | 36.79656 | 25.54318 | 94 | -0.00211 | -0.00339 | -0.00179 |
| GATA3 | 7 | 21.0369 | 13.21094 | 12 | 0.002416 | 0.001237 | -0.00175 |
| AMPKALPHA | 4 | 12.68017 | 4.800066 | 0 | 0.00155 | -0.00018 | -0.00172 |
| CDK1 | 4 | 12.85717 | 7.468792 | 0 | 0.001596 | 0.000681 | -0.00172 |
| MRE11 | 4 | 12.64561 | 9.72056 | 0 | 0.001542 | 0.001404 | -0.00172 |
| MTOR | 4 | 9.447903 | 13.81265 | 0 | 0.000718 | 0.002717 | -0.00172 |
| PDK1_pS241 | 4 | 12.79566 | 15.25653 | 0 | 0.00158 | 0.003181 | -0.00172 |
| PEA15 | 4 | 18.46315 | 9.322498 | 0 | 0.00304 | 0.001276 | -0.00172 |
| CD20 | 4 | 13.70306 | 6.825651 | 0 | 0.001814 | 0.000474 | -0.00172 |
| FOXM1 | 4 | 15.23122 | 5.923124 | 0 | 0.002208 | 0.000185 | -0.00172 |
| TAZ | 4 | 14.6822 | 5.720105 | 0 | 0.002066 | 0.00012 | -0.00172 |
| CMET | 4 | 12.25754 | 7.412353 | 0 | 0.001442 | 0.000663 | -0.00172 |
| ERCC5 | 4 | 11.36073 | 14.9963 | 0 | 0.00121 | 0.003097 | -0.00172 |
| SHP2_pY542 | 4 | 16.13566 | 8.669419 | 0 | 0.002441 | 0.001066 | -0.00172 |
| GAB2 | 19 | 25.28426 | 25.0156 | 62 | -0.00164 | -0.00012 | -0.00169 |
| ECADHERIN | 38 | 30.50338 | 35.90347 | 141 | -0.00845 | -0.00478 | -0.00162 |
| MSH2 | 4 | 6.424011 | 6.829493 | 1 | -6.15E-05 | 0.000476 | -0.00161 |
| P53 | 15 | 18.60773 | 21.64809 | 47 | -0.00164 | 0.000512 | -0.00154 |
| BCLXL | 3 | 11.51317 | 3.748193 | 0 | 0.001679 | -8.43E-05 | -0.00129 |
| CD31 | 3 | 10.74432 | 3.859435 | 0 | 0.001481 | -4.86E-05 | -0.00129 |
| JNK_pT183Y185 | 3 | 13.76088 | 9.275148 | 0 | 0.002258 | 0.00169 | -0.00129 |
| NF2 | 3 | 9.354397 | 5.244429 | 0 | 0.001123 | 0.000396 | -0.00129 |
| P27 | 3 | 8.120294 | 4.753921 | 0 | 0.000805 | 0.000239 | -0.00129 |
| PKCDELTA_pS664 | 3 | 12.71995 | 5.387529 | 0 | 0.00199 | 0.000442 | -0.00129 |
| YB1 | 3 | 11.07566 | 9.44243 | 0 | 0.001566 | 0.001744 | -0.00129 |
| BRCA2 | 3 | 14.21906 | 6.192743 | 0 | 0.002376 | 0.0007 | -0.00129 |
| PDK1 | 3 | 12.59776 | 7.424184 | 0 | 0.001958 | 0.001096 | -0.00129 |
| PRDX1 | 3 | 15.03124 | 6.047536 | 0 | 0.002585 | 0.000654 | -0.00129 |
| RICTOR_pT1135 | 3 | 10.91659 | 7.159239 | 0 | 0.001525 | 0.001011 | -0.00129 |
| SF2 | 3 | 8.351179 | 7.196741 | 0 | 0.000864 | 0.001023 | -0.00129 |
| CABL | 3 | 12.72788 | 4.635065 | 0 | 0.001992 | 0.0002 | -0.00129 |
| CD26 | 3 | 10.5782 | 3.997612 | 0 | 0.001438 | -4.23E-06 | -0.00129 |
| HSP70 | 20 | 37.06648 | 20.62705 | 71 | 0.000967 | -0.00196 | -0.00118 |
| HER3_pY1289 | 3 | 11.84093 | 8.543801 | 1 | 0.001763 | 0.001455 | -0.00118 |
| JAB1 | 3 | 7.727355 | 4.06251 | 1 | 0.000703 | 1.66E-05 | -0.00118 |
| CASPASE3 | 3 | 9.639759 | 12.87095 | 1 | 0.001196 | 0.002844 | -0.00118 |
| HER2 | 23 | 35.42507 | 27.03615 | 84 | -0.00074 | -0.00119 | -0.00112 |
| MSH6 | 6 | 17.71023 | 14.28987 | 14 | 0.001988 | 0.002012 | -0.00112 |
| NCADHERIN | 12 | 29.63753 | 17.50515 | 39 | 0.002486 | 0.000469 | -0.00109 |
| ADAR1 | 3 | 6.765181 | 7.564627 | 2 | 0.000456 | 0.001141 | -0.00108 |
| X4EBP1_pS65 | 2 | 7.959395 | 1.853678 | 0 | 0.001192 | -0.00026 | -0.00086 |
| CRAF_pS338 | 2 | 11.30009 | 6.653147 | 0 | 0.002053 | 0.001277 | -0.00086 |
| CHK1 | 2 | 9.772633 | 3.837932 | 0 | 0.00166 | 0.000374 | -0.00086 |
| DVL3 | 2 | 1.249558 | 2.591062 | 0 | -0.00054 | -2.66E-05 | -0.00086 |
| GSK3ALPHABETA | 2 | 5.653147 | 12.30009 | 0 | 0.000598 | 0.00309 | -0.00086 |
| LKB1 | 2 | 9.616057 | 4.788037 | 0 | 0.001619 | 0.000679 | -0.00086 |
| P90RSK_pT359S363 | 2 | 10.34623 | 3.207672 | 0 | 0.001807 | 0.000171 | -0.00086 |
| STATHMIN | 2 | 5.057069 | 5.328241 | 0 | 0.000445 | 0.000852 | -0.00086 |
| YB1_pS102 | 2 | 10.09448 | 4.559704 | 0 | 0.001743 | 0.000605 | -0.00086 |
| PI3KP85 | 2 | 4.715529 | 13.318 | 0 | 0.000357 | 0.003417 | -0.00086 |
| RAB11 | 2 | 11.79988 | 4.915778 | 0 | 0.002182 | 0.00072 | -0.00086 |
| TSC1 | 2 | 7.187402 | 14.64739 | 0 | 0.000994 | 0.003844 | -0.00086 |
| TUBERIN_pT1462 | 2 | 12.1839 | 5.581954 | 0 | 0.002281 | 0.000934 | -0.00086 |
| X1433BETA | 2 | 9.709952 | 2.635995 | 0 | 0.001644 | -1.22E-05 | -0.00086 |
| X1433ZETA | 2 | 11.21586 | 3.877478 | 0 | 0.002032 | 0.000386 | -0.00086 |
| ACVRL1 | 2 | 10.27249 | 3.936255 | 0 | 0.001788 | 0.000405 | -0.00086 |
| BRAF_pS445 | 2 | 6.858078 | 7.603502 | 0 | 0.000909 | 0.001583 | -0.00086 |
| SMAD1 | 2 | 6.826632 | 7.40241 | 1 | 0.000901 | 0.001518 | -0.00075 |
| BAK | 13 | 25.24467 | 13.77258 | 47 | 0.000925 | -0.00116 | -0.00068 |
| AKT_pT308 | 14 | 33.16884 | 17.26436 | 53 | 0.002538 | -0.00047 | -0.00049 |
| CDK1_pY15 | 4 | 11.46624 | 12.23314 | 12 | 0.001238 | 0.00221 | -0.00047 |
| BAX | 1 | 7.470814 | 4.632131 | 0 | 0.001496 | 0.001058 | -0.00043 |
| CMET_pY1235 | 1 | 5.539735 | 1.669477 | 0 | 0.000998 | 0.000107 | -0.00043 |
| CRAF | 1 | 6.382726 | 3.091278 | 0 | 0.001215 | 0.000563 | -0.00043 |
| CYCLIND1 | 1 | 9.924171 | 4.619625 | 0 | 0.002128 | 0.001054 | -0.00043 |
| DJ1 | 1 | 1.125485 | 7.525032 | 0 | -0.00014 | 0.001987 | -0.00043 |
| FOXO3A | 1 | 6.525032 | 2.125485 | 0 | 0.001252 | 0.000253 | -0.00043 |
| IRS1 | 1 | 9.924171 | 4.619625 | 0 | 0.002128 | 0.001054 | -0.00043 |
| JNK2 | 1 | 2.595844 | 0.751102 | 0 | 0.00024 | -0.00019 | -0.00043 |
| MIG6 | 1 | 7.456339 | 2.039371 | 0 | 0.001492 | 0.000226 | -0.00043 |
| MTOR_pS2448 | 1 | 7.470814 | 4.632131 | 0 | 0.001496 | 0.001058 | -0.00043 |
| P27_pT157 | 1 | 7.188689 | 2.582216 | 0 | 0.001423 | 0.0004 | -0.00043 |
| P38MAPK | 1 | 3.632131 | 8.470814 | 0 | 0.000507 | 0.00229 | -0.00043 |
| PI3KP110ALPHA | 1 | 7.043481 | 3.766032 | 0 | 0.001386 | 0.00078 | -0.00043 |
| PR | 1 | 9.924171 | 4.619625 | 0 | 0.002128 | 0.001054 | -0.00043 |
| PRAS40_pT246 | 1 | 7.456339 | 2.039371 | 0 | 0.001492 | 0.000226 | -0.00043 |
| SHC_pY317 | 1 | 3.697557 | 1.285598 | 0 | 0.000524 | -1.65E-05 | -0.00043 |
| SMAD4 | 1 | 7.456339 | 2.039371 | 0 | 0.001492 | 0.000226 | -0.00043 |
| FOXO3A_pS318S321 | 1 | 4.163469 | 2.022438 | 0 | 0.000644 | 0.00022 | -0.00043 |
| NRAS | 1 | 6.525032 | 2.125485 | 0 | 0.001252 | 0.000253 | -0.00043 |
| P21 | 1 | 4.857687 | 1.879062 | 0 | 0.000822 | 0.000174 | -0.00043 |
| RAPTOR | 1 | 2.222961 | 1.5678 | 0 | 0.000144 | 7.41E-05 | -0.00043 |
| CASPASE8 | 1 | 9.924171 | 4.619625 | 0 | 0.002128 | 0.001054 | -0.00043 |
| RB | 1 | 3.947989 | 2.68788 | 0 | 0.000588 | 0.000434 | -0.00043 |
| BCL2A1 | 1 | 7.043481 | 3.766032 | 0 | 0.001386 | 0.00078 | -0.00043 |
| COG3 | 1 | 1.672084 | 5.665217 | 0 | 1.65E-06 | 0.001389 | -0.00043 |
| IRF1 | 1 | 7.188689 | 2.582216 | 0 | 0.001423 | 0.0004 | -0.00043 |
| JAK2 | 1 | 2.766032 | 8.043481 | 0 | 0.000284 | 0.002153 | -0.00043 |
| ACC1 | 26 | 27.89245 | 30.36498 | 104 | -0.00397 | -0.00141 | -0.00032 |
| BRAF | 22 | 25.28808 | 33.66225 | 89 | -0.00293 | 0.001364 | -0.00017 |
| RAB25 | 24 | 27.40619 | 26.77481 | 99 | -0.00324 | -0.00171 | 1.64E-05 |
| DIRAS3 | 4 | 13.18571 | 9.855073 | 21 | 0.001681 | 0.001447 | 0.000472 |
| SYK | 31 | 27.10415 | 30.45535 | 133 | -0.00632 | -0.00353 | 0.000555 |
| P38_pT180Y182 | 13 | 21.22555 | 19.3196 | 60 | -0.00011 | 0.000623 | 0.000673 |
| YAP_pS127 | 21 | 25.29718 | 25.51981 | 94 | -0.00249 | -0.00082 | 0.000783 |
| RB_pS807S811 | 19 | 28.39949 | 20.62476 | 86 | -0.00084 | -0.00153 | 0.000808 |
| PKCPANBETAII_pS660 | 18 | 22.3644 | 22.92299 | 84 | -0.00196 | -0.00037 | 0.001028 |
| GSK3_pS9 | 25 | 27.55692 | 32.52321 | 113 | -0.00363 | -0.00029 | 0.001046 |
| MEK1 | 15 | 21.56738 | 22.79221 | 86 | -0.00088 | 0.000879 | 0.002524 |
| FASN | 34 | 31.09772 | 34.43028 | 172 | -0.00658 | -0.00354 | 0.003332 |
| X4EBP1_pT37T46 | 23 | 31.45845 | 30.20208 | 127 | -0.00177 | -0.00018 | 0.003363 |
| DUSP4 | 11 | 24.36554 | 17.44836 | 78 | 0.001557 | 0.00088 | 0.003407 |
| ERK2 | 14 | 21.8656 | 24.55581 | 92 | -0.00037 | 0.001874 | 0.003579 |
| GSK3ALPHABETA_pS21S9 | 20 | 22.28957 | 35.82311 | 117 | -0.00284 | 0.002916 | 0.003609 |
| SRC_pY527 | 27 | 32.41445 | 24.1229 | 146 | -0.00324 | -0.00384 | 0.003627 |
| P16INK4A | 28 | 35.91133 | 33.65613 | 151 | -0.00276 | -0.00121 | 0.003719 |
| CLAUDIN7 | 18 | 32.16897 | 18.17295 | 112 | 0.000563 | -0.00189 | 0.003946 |
| ATM | 29 | 23.27587 | 33.06239 | 166 | -0.00645 | -0.00183 | 0.004853 |
| ETS1 | 19 | 32.00713 | 24.87453 | 147 | 9.25E-05 | -0.00017 | 0.007164 |
| SRC_pY416 | 25 | 33.08427 | 22.52777 | 173 | -0.00221 | -0.0035 | 0.007299 |
| FIBRONECTIN | 17 | 28.37836 | 24.62565 | 164 | 1.59E-05 | 0.000609 | 0.009794 |
| BETACATENIN | 42 | 33.59742 | 35.38309 | 270 | -0.00937 | -0.00667 | 0.010111 |
| PAXILLIN | 14 | 23.40277 | 26.01094 | 161 | 2.14E-05 | 0.002342 | 0.010769 |
| CYCLINB1 | 31 | 28.89112 | 33.05461 | 268 | -0.00586 | -0.00269 | 0.014624 |
| GAPDH | 45 | 32.44055 | 33.49971 | 407 | -0.01095 | -0.00856 | 0.0231 |
| CAVEOLIN1 | 36 | 36.33681 | 28.89145 | 387 | -0.00609 | -0.00618 | 0.024879 |
| ERALPHA | 25 | 28.55452 | 38.68051 | 390 | -0.00337 | 0.001688 | 0.029912 |
| PEA15_pS116 | 19 | 34.53927 | 22.12048 | 388 | 0.000745 | -0.00105 | 0.032279 |
| TFRC | 42 | 31.40973 | 34.70321 | 517 | -0.00993 | -0.00689 | 0.035851 |
| ANNEXIN1 | 31 | 35.30833 | 29.89554 | 498 | -0.00421 | -0.00371 | 0.038592 |
| ACETYLATUBULINLYS40 | 41 | 33.82117 | 36.24073 | 579 | -0.00888 | -0.00596 | 0.042741 |
| MYH11 | 38 | 35.28035 | 32.61 | 717 | -0.00722 | -0.00584 | 0.05841 |
| EPPK1 | 56 | 35.45187 | 35.69887 | 859 | -0.0149 | -0.01257 | 0.065482 |

**Supplementary Table 5.** Differential ratio values for the CT based analysis of the HPA dataset. Differential expression number, positive and negative correlation numbers, MAPP number and differential ratio values for the PCPP, NCPP, and MAPP were listed for all the proteins with a nonzero differential ratio value. Proteins were ranked from the lowest MAPP based differential ratio value to the highest.

| Protein symbol | Differential expression number | PC number | NC number | MAPP number | Differential ratio for PCPP | Differential ratio for NCPP | Differential ratio for MAPP |
| --- | --- | --- | --- | --- | --- | --- | --- |
| SUGP2 | 4 | 433.7274 | 62.84821 | 1 | -0.00028 | -0.00048 | 0.002683 |
| SNX2 | 4 | 262.2635 | 87.95403 | 1 | -0.00041 | -0.00042 | 0.002683 |
| KHDRBS3 | 4 | 446.4952 | 55.63704 | 1 | -0.00027 | -0.00049 | 0.002683 |
| CYBRD1 | 3 | 353.5961 | 97.39691 | 1 | -0.00019 | -0.00025 | 0.002835 |
| ATP8B2 | 3 | 345.5912 | 77.51213 | 1 | -0.00019 | -0.00029 | 0.002835 |
| KPNA1 | 3 | 353.5961 | 97.39691 | 1 | -0.00019 | -0.00025 | 0.002835 |
| RAB31 | 3 | 223.4532 | 125.7163 | 1 | -0.00028 | -0.00019 | 0.002835 |
| PM20D2 | 3 | 275.7929 | 152.5707 | 1 | -0.00024 | -0.00014 | 0.002835 |
| TRAIP | 3 | 400.6711 | 125.8376 | 1 | -0.00015 | -0.00019 | 0.002835 |
| NAP1L1 | 3 | 446.8053 | 59.63838 | 1 | -0.00011 | -0.00033 | 0.002835 |
| GSN | 3 | 290.3832 | 54.42483 | 1 | -0.00023 | -0.00034 | 0.002835 |
| LARS | 3 | 235.6194 | 139.7773 | 1 | -0.00028 | -0.00017 | 0.002835 |
| MRPL33 | 3 | 525.2933 | 44.83575 | 1 | -5.46E-05 | -0.00036 | 0.002835 |
| SKIL | 3 | 321.0783 | 171.0094 | 1 | -0.00021 | -0.0001 | 0.002835 |
| PRDX6 | 3 | 286.0249 | 130.1058 | 1 | -0.00024 | -0.00019 | 0.002835 |
| RAF1 | 3 | 424.0977 | 124.0067 | 1 | -0.00013 | -0.0002 | 0.002835 |
| GAPDH | 2 | 285.4278 | 88.64367 | 1 | -8.58E-05 | -0.00012 | 0.002986 |
| NES | 2 | 207.7274 | 210.8945 | 1 | -0.00014 | 0.000133 | 0.002986 |
| ALKBH4 | 2 | 219.3786 | 136.5195 | 1 | -0.00014 | -2.05E-05 | 0.002986 |
| TMC8 | 2 | 219.3786 | 136.5195 | 1 | -0.00014 | -2.05E-05 | 0.002986 |
| HADHA | 2 | 256.5524 | 62.76867 | 1 | -0.00011 | -0.00017 | 0.002986 |
| FIBP | 2 | 244.3944 | 73.32068 | 1 | -0.00012 | -0.00015 | 0.002986 |
| P3H3 | 2 | 336.3504 | 52.32552 | 1 | -4.69E-05 | -0.00019 | 0.002986 |
| ST3GAL3 | 2 | 336.3504 | 52.32552 | 1 | -4.69E-05 | -0.00019 | 0.002986 |
| FITM1 | 2 | 336.3504 | 52.32552 | 1 | -4.69E-05 | -0.00019 | 0.002986 |
| RNF212 | 2 | 336.3504 | 52.32552 | 1 | -4.69E-05 | -0.00019 | 0.002986 |
| PFAS | 2 | 205.7242 | 41.11629 | 1 | -0.00015 | -0.00022 | 0.002986 |
| NAGA | 2 | 211.1404 | 168.8005 | 1 | -0.00014 | 4.63E-05 | 0.002986 |
| AK3 | 4 | 482.676 | 119.6816 | 2 | -0.00024 | -0.00036 | 0.005972 |
| KIAA1211L | 3 | 201.2535 | 103.5664 | 2 | -0.0003 | -0.00024 | 0.006124 |
| RBM12 | 3 | 71.47876 | 246.171 | 2 | -0.0004 | 5.49E-05 | 0.006124 |
| GFOD2 | 3 | 443.2914 | 57.37266 | 2 | -0.00012 | -0.00034 | 0.006124 |
| NELFE | 3 | 514.8013 | 130.235 | 2 | -6.26E-05 | -0.00019 | 0.006124 |
| HMGB2 | 2 | 236.6227 | 201.8908 | 2 | -0.00012 | 0.000115 | 0.006276 |
| FAP | 2 | 68.35171 | 223.81 | 2 | -0.00025 | 0.00016 | 0.006276 |
| SIRT2 | 2 | 345.4504 | 178.1623 | 2 | -4.00E-05 | 6.57E-05 | 0.006276 |
| FAM81A | 2 | 345.4504 | 178.1623 | 2 | -4.00E-05 | 6.57E-05 | 0.006276 |
| ARL17A | 2 | 196.1834 | 245.275 | 2 | -0.00015 | 0.000205 | 0.006276 |
| ARL17B | 2 | 196.1834 | 245.275 | 2 | -0.00015 | 0.000205 | 0.006276 |
| MAN1A2 | 2 | 196.1834 | 245.275 | 2 | -0.00015 | 0.000205 | 0.006276 |
| PSMG4 | 2 | 196.1834 | 245.275 | 2 | -0.00015 | 0.000205 | 0.006276 |
| DMWD | 4 | 164.8511 | 79.52492 | 3 | -0.00048 | -0.00044 | 0.009262 |
| CRISPLD1 | 3 | 420.7921 | 64.15508 | 3 | -0.00013 | -0.00032 | 0.009413 |
| WNK2 | 3 | 202.5984 | 166.9741 | 3 | -0.0003 | -0.00011 | 0.009413 |
| CKMT1A | 3 | 70.34573 | 168.5737 | 3 | -0.0004 | -0.00011 | 0.009413 |
| CKMT1B | 3 | 70.34573 | 168.5737 | 3 | -0.0004 | -0.00011 | 0.009413 |
| CRIP1 | 2 | 209.8945 | 208.7274 | 3 | -0.00014 | 0.000129 | 0.009565 |
| HSPB1 | 2 | 226.9874 | 96.59252 | 3 | -0.00013 | -0.0001 | 0.009565 |
| SLC39A6 | 2 | 226.9874 | 96.59252 | 3 | -0.00013 | -0.0001 | 0.009565 |
| PARD6A | 2 | 95.59252 | 227.9874 | 3 | -0.00023 | 0.000169 | 0.009565 |
| PROM1 | 2 | 96.0159 | 136.5334 | 3 | -0.00023 | -2.05E-05 | 0.009565 |
| AP4S1 | 2 | 200.8908 | 237.6227 | 3 | -0.00015 | 0.000189 | 0.009565 |
| ACSBG1 | 2 | 72.32068 | 245.3944 | 3 | -0.00025 | 0.000205 | 0.009565 |
| BZW2 | 2 | 167.8005 | 212.1404 | 3 | -0.00018 | 0.000136 | 0.009565 |
| NFIX | 4 | 293.7006 | 195.8505 | 4 | -0.00038 | -0.0002 | 0.012551 |
| ZNF384 | 4 | 159.5719 | 165.8826 | 4 | -0.00048 | -0.00026 | 0.012551 |
| ISPD | 4 | 195.7114 | 127.01 | 4 | -0.00046 | -0.00034 | 0.012551 |
| RNGTT | 4 | 248.7075 | 153.6355 | 4 | -0.00042 | -0.00029 | 0.012551 |
| ENHO | 3 | 239.7608 | 212.0289 | 4 | -0.00027 | -1.58E-05 | 0.012703 |
| PHF14 | 3 | 188.3645 | 68.70382 | 4 | -0.00031 | -0.00031 | 0.012703 |
| H1FX | 2 | 311.431 | 191.073 | 4 | -6.59E-05 | 9.24E-05 | 0.012855 |
| ARHGAP31 | 2 | 311.431 | 191.073 | 4 | -6.59E-05 | 9.24E-05 | 0.012855 |
| SNAPC2 | 2 | 224.3921 | 271.4833 | 4 | -0.00013 | 0.000259 | 0.012855 |
| GLI3 | 2 | 311.431 | 191.073 | 4 | -6.59E-05 | 9.24E-05 | 0.012855 |
| TAF12 | 2 | 224.3921 | 271.4833 | 4 | -0.00013 | 0.000259 | 0.012855 |
| SLC20A2 | 2 | 224.3921 | 271.4833 | 4 | -0.00013 | 0.000259 | 0.012855 |
| FZD7 | 2 | 270.4833 | 225.3921 | 4 | -9.72E-05 | 0.000164 | 0.012855 |
| DDX39B | 2 | 244.275 | 197.1834 | 4 | -0.00012 | 0.000105 | 0.012855 |
| TPD52 | 2 | 177.1623 | 346.4504 | 4 | -0.00017 | 0.000414 | 0.012855 |
| S100A14 | 2 | 40.11629 | 206.7242 | 4 | -0.00027 | 0.000125 | 0.012855 |
| PATJ | 4 | 425.2946 | 185.2324 | 5 | -0.00028 | -0.00022 | 0.015841 |
| ZBTB7B | 3 | 313.3353 | 129.2165 | 5 | -0.00022 | -0.00019 | 0.015992 |
| GJC3 | 3 | 299.7251 | 184.6579 | 5 | -0.00023 | -7.25E-05 | 0.015992 |
| WT1 | 3 | 299.7251 | 184.6579 | 5 | -0.00023 | -7.25E-05 | 0.015992 |
| PRDM1 | 2 | 248.0874 | 162.9945 | 5 | -0.00011 | 3.43E-05 | 0.016144 |
| AGR3 | 2 | 248.0874 | 162.9945 | 5 | -0.00011 | 3.43E-05 | 0.016144 |
| FSCN1 | 2 | 190.073 | 312.431 | 5 | -0.00016 | 0.000344 | 0.016144 |
| YARS | 2 | 190.073 | 312.431 | 5 | -0.00016 | 0.000344 | 0.016144 |
| LAMB2 | 2 | 190.073 | 312.431 | 5 | -0.00016 | 0.000344 | 0.016144 |
| EID2B | 2 | 161.9945 | 249.0874 | 5 | -0.00018 | 0.000213 | 0.016144 |
| GIGYF1 | 5 | 292.8103 | 181.5983 | 6 | -0.00054 | -0.00038 | 0.018979 |
| RNASEH2C | 3 | 275.9014 | 94.57424 | 6 | -0.00024 | -0.00026 | 0.019282 |
| TRPS1 | 3 | 198.2924 | 132.0949 | 8 | -0.0003 | -0.00018 | 0.025861 |
| KRT5 | 4 | 129.7756 | 244.0795 | 9 | -0.00051 | -0.0001 | 0.028999 |
| KRT17 | 4 | 122.9086 | 171.8879 | 9 | -0.00051 | -0.00025 | 0.028999 |
| SPIDR | 3 | 211.5596 | 264.719 | 9 | -0.00029 | 9.33E-05 | 0.02915 |
| PKP3 | 5 | 72.09583 | 252.0518 | 11 | -0.0007 | -0.00024 | 0.035426 |
| ABHD3 | 4 | 125.7461 | 186.7768 | 11 | -0.00051 | -0.00022 | 0.035578 |
| PC | 8 | 334.9578 | 114.2466 | 13 | -0.00096 | -0.00098 | 0.04155 |
| LRRC26 | 5 | 119.1873 | 206.4925 | 15 | -0.00067 | -0.00033 | 0.048584 |

**Supplementary Table 6.** Significant MAPP of the TCPA dataset. Whether the pair is present in only CS, only CT or both CS and CT based analyses were listed. Presence of direct PPI, indirect PPI and direct PDI were also given.

| Protein 1 | Protein 2 | MAPP type | Direct PPI type | Indirect PPI type | PDI type |
| --- | --- | --- | --- | --- | --- |
| TUBA1B | ANXA1 | Both | None | None | None |
| TUBA1B | ESR1 | Both | None | Both | None |
| TUBA1B | FN1 | CS | None | None | None |
| TUBA1B | RAB25 | CS | None | None | None |
| TUBA1B | PEA15 | CS | None | None | None |
| TUBA1B | EIF4EBP1 | Both | None | None | None |
| TUBA1B | ATM | CS | None | None | None |
| TUBA1B | YAP1 | Both | None | Both | None |
| TUBA1B | CCNE2 | CS | None | None | None |
| TUBA1B | INPP4B | CS | None | None | None |
| TUBA1B | ERBB2 | CS | None | CS | None |
| TUBA1B | CDH2 | CS | None | None | None |
| TUBA1B | MAPK14 | CS | None | None | None |
| TUBA1B | RAD50 | CS | None | None | None |
| EPPK1 | TFRC | Both | None | None | None |
| EPPK1 | BRAF | Both | None | None | None |
| EPPK1 | GAPDH | CS | None | None | None |
| EPPK1 | RB1 | CS | None | None | None |
| EPPK1 | CTNNB1 | CS | None | None | None |
| EPPK1 | FN1 | Both | None | None | None |
| EPPK1 | CDKN2A | CS | None | None | None |
| EPPK1 | FASN | Both | None | None | None |
| EPPK1 | SRC | Both | None | None | None |
| EPPK1 | RAB25 | Both | None | None | None |
| EPPK1 | SYK | Both | None | None | None |
| EPPK1 | EIF4EBP1 | Both | None | None | None |
| EPPK1 | PXN | Both | None | None | None |
| EPPK1 | ATM | CS | None | None | None |
| EPPK1 | INPP4B | Both | None | None | None |
| EPPK1 | MAPK14 | Both | None | None | None |
| EPPK1 | EEF2 | Both | None | None | None |
| EPPK1 | SQSTM1 | Both | None | None | None |
| EPPK1 | GAB2 | Both | None | None | None |
| EPPK1 | PDCD4 | CS | None | None | None |
| EPPK1 | MSH6 | CS | None | None | None |
| EPPK1 | RPS6 | CS | None | None | None |
| EPPK1 | TP53BP1 | Both | None | None | None |
| EPPK1 | GSK3A | CS | None | None | None |
| EPPK1 | GSK3B | CS | None | None | None |
| EPPK1 | ACACA | Both | None | None | None |
| EPPK1 | ACACB | Both | None | None | None |
| EPPK1 | TP53 | Both | None | None | None |
| EPPK1 | CHEK2 | CS | None | None | None |
| EPPK1 | CDK1 | Both | None | None | None |
| CAV1 | TFRC | Both | None | None | None |
| CAV1 | BRAF | Both | None | None | None |
| CAV1 | CDKN2A | Both | None | None | None |
| CAV1 | SRC | Both | Both | Both | None |
| CAV1 | CLDN7 | CS | None | None | None |
| CAV1 | PEA15 | CS | None | None | None |
| CAV1 | EIF4EBP1 | CS | None | None | None |
| CAV1 | MAP2K1 | Both | None | Both | None |
| CAV1 | INPP4B | CS | None | None | None |
| CAV1 | SQSTM1 | CS | CS | CS | None |
| CAV1 | GSK3A | CS | None | None | None |
| CAV1 | GSK3B | CS | None | CS | None |
| CAV1 | PRRT2 | Both | None | None | None |
| CAV1 | AKT1 | Both | None | Both | None |
| CAV1 | AKT2 | Both | None | Both | None |
| CAV1 | AKT3 | Both | None | None | None |
| CAV1 | DUSP4 | Both | None | None | None |
| CAV1 | COPS5 | CS | None | CS | None |
| CAV1 | PTEN | Both | None | Both | None |
| MYH11 | GAPDH | CS | None | None | None |
| MYH11 | RB1 | Both | None | None | None |
| MYH11 | IGFBP2 | Both | None | None | None |
| MYH11 | ANXA1 | CS | None | None | None |
| MYH11 | CTNNB1 | Both | None | None | None |
| MYH11 | FN1 | Both | None | None | None |
| MYH11 | CDKN2A | Both | None | None | None |
| MYH11 | SRC | Both | None | None | None |
| MYH11 | PEA15 | CS | None | None | None |
| MYH11 | ATM | CS | None | None | None |
| MYH11 | ERBB2 | CS | None | None | None |
| MYH11 | CDH2 | CS | None | None | None |
| MYH11 | MAPK14 | Both | None | None | None |
| MYH11 | GSK3A | Both | None | None | None |
| MYH11 | GSK3B | Both | None | None | None |
| MYH11 | AKT1 | Both | None | None | None |
| MYH11 | AKT2 | Both | None | None | None |
| MYH11 | AKT3 | Both | None | None | None |
| MYH11 | RPS6KB1 | Both | None | None | None |
| MYH11 | ETS1 | Both | None | None | None |
| MYH11 | ERBB3 | Both | None | None | None |
| MYH11 | CCNE1 | Both | None | None | None |
| TFRC | BRAF | Both | None | Both | None |
| TFRC | ANXA1 | Both | None | None | None |
| TFRC | ESR1 | CS | None | None | None |
| TFRC | CTNNB1 | CS | None | CS | None |
| TFRC | FASN | CS | None | None | None |
| TFRC | SRC | CS | None | CS | None |
| TFRC | PXN | CS | None | None | None |
| TFRC | MAPK14 | Both | None | None | None |
| TFRC | TP53BP1 | Both | None | None | None |
| TFRC | ACACA | Both | None | None | None |
| TFRC | PTEN | Both | None | None | None |
| TFRC | ETS1 | CS | None | None | None |
| TFRC | STAT5A | Both | None | None | None |
| BRAF | ANXA1 | Both | None | None | None |
| BRAF | ESR1 | Both | None | Both | None |
| BRAF | YAP1 | Both | None | Both | None |
| GAPDH | IGFBP2 | CS | None | CS | None |
| GAPDH | ESR1 | Both | None | Both | None |
| GAPDH | CCNB1 | CS | None | CS | None |
| GAPDH | CDH1 | CS | None | CS | None |
| GAPDH | SRC | CS | None | CS | None |
| GAPDH | SYK | CT | None | None | None |
| GAPDH | EIF4EBP1 | CS | None | None | None |
| GAPDH | MAP2K1 | Both | None | None | None |
| GAPDH | YAP1 | Both | None | Both | None |
| GAPDH | ACACA | CT | None | None | None |
| GAPDH | DUSP4 | CS | None | None | None |
| GAPDH | MSH2 | CS | None | CS | None |
| RB1 | FN1 | Both | None | Both | None |
| IGFBP2 | ANXA1 | CS | None | None | None |
| IGFBP2 | PEA15 | CS | None | None | None |
| ANXA1 | SRC | CS | None | CS | None |
| ANXA1 | BAK1 | CS | None | None | None |
| ANXA1 | MAP2K1 | CT | None | None | None |
| ANXA1 | ATM | CS | None | CS | None |
| ANXA1 | INPP4B | CS | None | None | None |
| ANXA1 | RAD50 | CS | None | CS | None |
| ANXA1 | GAB2 | CS | None | None | None |
| ANXA1 | GSK3A | Both | None | None | None |
| ANXA1 | GSK3B | Both | None | Both | None |
| ANXA1 | ACACA | CS | None | None | None |
| ANXA1 | ACACB | CS | None | None | None |
| ANXA1 | PRRT2 | CS | None | None | None |
| ANXA1 | AR | CS | None | CS | CS |
| ESR1 | CTNNB1 | CS | None | CS | CS |
| ESR1 | FN1 | CT | None | None | None |
| ESR1 | CDKN2A | Both | None | Both | None |
| ESR1 | BAK1 | CS | None | CS | None |
| ESR1 | SYK | Both | None | Both | None |
| ESR1 | PXN | Both | None | Both | None |
| ESR1 | MAP2K1 | CS | None | CS | None |
| ESR1 | EEF2 | CT | None | None | None |
| ESR1 | SQSTM1 | CT | None | CT | None |
| ESR1 | GSK3A | CS | None | CS | None |
| ESR1 | GSK3B | CS | None | CS | None |
| ESR1 | PRRT2 | Both | None | None | None |
| ESR1 | ETS1 | CS | None | CS | None |
| ESR1 | STAT5A | Both | Both | Both | CS |
| ESR1 | PRKAA1 | CS | None | CS | None |
| CCNB1 | CTNNB1 | CT | None | CT | None |
| CCNB1 | CDH1 | CS | None | None | None |
| CCNB1 | SRC | CT | None | CT | None |
| CCNB1 | SYK | Both | None | None | None |
| CCNB1 | MAP2K1 | CS | None | CS | None |
| CCNB1 | MAPK14 | CS | None | CS | None |
| CCNB1 | PDCD4 | CS | None | CS | None |
| CCNB1 | GSK3A | CS | None | None | None |
| CCNB1 | GSK3B | CS | None | CS | None |
| CTNNB1 | PEA15 | CT | None | CT | None |
| CTNNB1 | YAP1 | Both | Both | Both | None |
| CTNNB1 | RAD50 | CS | None | CS | None |
| CTNNB1 | RPS6 | CS | None | None | None |
| CTNNB1 | ACACA | CS | None | CS | None |
| CDH1 | FASN | CS | None | None | None |
| CDH1 | PEA15 | CT | None | None | None |
| CDH1 | ATM | Both | None | Both | None |
| CDH1 | ACACA | Both | None | None | None |
| FN1 | SRC | CS | None | CS | None |
| FN1 | MAPK1 | CS | None | None | None |
| FN1 | PRRT2 | CT | None | None | None |
| CDKN2A | ERBB2 | CS | None | CS | None |
| CDKN2A | AKT1 | CT | None | CT | None |
| CDKN2A | AKT2 | CT | None | None | None |
| CDKN2A | AKT3 | CT | None | None | None |
| CDKN2A | RICTOR | Both | None | Both | None |
| HSPA1A | MAPK1 | Both | None | Both | None |
| HSPA1A | RAD50 | CS | None | None | None |
| HSPA1A | PRRT2 | CS | None | None | None |
| HSPA1A | PARP1 | Both | None | Both | None |
| FASN | SYK | Both | None | None | None |
| FASN | PRRT2 | Both | None | None | None |
| FASN | PRKAA1 | CS | None | CS | None |
| SRC | PEA15 | Both | None | Both | None |
| SRC | PXN | Both | None | Both | None |
| SRC | ATM | Both | None | Both | None |
| SRC | CCNE2 | CS | None | CS | None |
| SRC | TP53BP1 | CS | None | None | None |
| SRC | ETS1 | Both | None | None | None |
| BAK1 | CLDN7 | Both | None | None | None |
| RAB25 | PXN | Both | None | None | None |
| CLDN7 | PEA15 | Both | None | None | None |
| CLDN7 | GAB2 | CS | None | None | None |
| CLDN7 | RPS6 | Both | None | None | None |
| PEA15 | PXN | Both | None | Both | None |
| PEA15 | EEF2 | Both | None | None | None |
| PEA15 | RPS6 | CS | None | CS | None |
| PEA15 | ACACA | CS | None | None | None |
| PEA15 | DUSP4 | Both | None | Both | None |
| PEA15 | RICTOR | Both | None | None | None |
| PEA15 | TGM2 | CS | None | None | None |
| PEA15 | KDR | Both | None | None | None |
| SYK | YAP1 | CS | None | CS | None |
| SYK | INPP4B | CS | None | None | None |
| SYK | GSK3A | Both | None | Both | None |
| SYK | GSK3B | Both | None | Both | None |
| MAPK1 | YAP1 | CS | None | CS | None |
| MAPK1 | INPP4B | CT | None | None | None |
| MAPK1 | PDCD4 | Both | None | None | None |
| MAPK1 | TGM2 | Both | None | None | None |
| EIF4EBP1 | STAT5A | Both | None | None | None |
| PXN | YAP1 | Both | None | Both | None |
| MAP2K1 | YAP1 | Both | Both | Both | None |
| ATM | MAPK14 | Both | None | Both | None |
| YAP1 | TP53BP1 | Both | None | Both | None |
| YAP1 | RPS6KB1 | CT | None | None | None |

**Supplementary Table 7.** Significant MAPP of the HPA dataset. Presence of PPI were also given.

| Protein 1 | Protein 2 | PPI type |
| --- | --- | --- |
| PRDM1 | KRT5 | None |
| PRDM1 | KRT17 | None |
| PRDM1 | EID2B | None |
| PRDM1 | LRRC26 | None |
| PRDM1 | ABHD3 | None |
| TRPS1 | KRT5 | None |
| TRPS1 | KRT17 | None |
| TRPS1 | EID2B | None |
| TRPS1 | LRRC26 | None |
| TRPS1 | ABHD3 | None |
| ZBTB7B | KRT5 | None |
| ZBTB7B | KRT17 | None |
| ZBTB7B | EID2B | None |
| ZBTB7B | LRRC26 | None |
| ZBTB7B | ABHD3 | None |
| SPIDR | KRT5 | None |
| SPIDR | KRT17 | None |
| SPIDR | EID2B | None |
| SPIDR | LRRC26 | None |
| SPIDR | ABHD3 | None |
| SPIDR | GJC3 | None |
| SPIDR | WT1 | None |
| SPIDR | NFIX | None |
| SPIDR | FZD7 | None |
| AGR3 | EID2B | None |
| AGR3 | LRRC26 | None |
| AGR3 | ABHD3 | None |
| CYBRD1 | LRRC26 | None |
| ATP8B2 | LRRC26 | None |
| GAPDH | LRRC26 | None |
| KPNA1 | LRRC26 | None |
| HSPB1 | LRRC26 | None |
| HSPB1 | ABHD3 | None |
| HSPB1 | PARD6A | Indirect |
| SLC39A6 | LRRC26 | None |
| SLC39A6 | ABHD3 | None |
| SLC39A6 | PARD6A | None |
| RNASEH2C | PKP3 | None |
| RNASEH2C | CKMT1A | None |
| RNASEH2C | CKMT1B | None |
| RNASEH2C | LRRC26 | None |
| RNASEH2C | ABHD3 | None |
| RNASEH2C | PARD6A | None |
| SNAPC2 | GJC3 | None |
| SNAPC2 | WT1 | None |
| SNAPC2 | NFIX | None |
| SNAPC2 | FZD7 | None |
| TAF12 | GJC3 | None |
| TAF12 | WT1 | None |
| TAF12 | NFIX | None |
| TAF12 | FZD7 | None |
| SLC20A2 | GJC3 | None |
| SLC20A2 | WT1 | None |
| SLC20A2 | NFIX | None |
| SLC20A2 | FZD7 | None |
| CRIP1 | GJC3 | None |
| CRIP1 | WT1 | None |
| CRIP1 | NES | None |
| CRISPLD1 | KRT17 | None |
| CRISPLD1 | LRRC26 | None |
| CRISPLD1 | ABHD3 | None |
| DMWD | KRT17 | None |
| DMWD | LRRC26 | None |
| DMWD | ABHD3 | None |
| ENHO | FSCN1 | None |
| ENHO | PATJ | None |
| ENHO | YARS | None |
| ENHO | LAMB2 | None |
| H1FX | FSCN1 | None |
| H1FX | PATJ | None |
| H1FX | YARS | None |
| H1FX | LAMB2 | None |
| ARHGAP31 | FSCN1 | None |
| ARHGAP31 | PATJ | None |
| ARHGAP31 | YARS | None |
| ARHGAP31 | LAMB2 | None |
| GLI3 | FSCN1 | None |
| GLI3 | PATJ | None |
| GLI3 | YARS | None |
| GLI3 | LAMB2 | None |
| PC | FSCN1 | None |
| PC | YARS | None |
| PC | LAMB2 | None |
| PC | AP4S1 | None |
| PC | ZNF384 | None |
| PC | ISPD | None |
| PC | FAP | None |
| PC | RBM12 | None |
| PC | PM20D2 | None |
| PC | TRAIP | None |
| KRT5 | KIAA1211L | None |
| KRT5 | ALKBH4 | None |
| KRT5 | TMC8 | None |
| KRT5 | PROM1 | None |
| KRT17 | PROM1 | None |
| HMGB2 | AP4S1 | None |
| HMGB2 | ZNF384 | None |
| WNK2 | AP4S1 | None |
| WNK2 | ZNF384 | None |
| WNK2 | GIGYF1 | None |
| PKP3 | PHF14 | None |
| PKP3 | ISPD | None |
| PKP3 | P3H3 | None |
| PKP3 | ST3GAL3 | None |
| PKP3 | GSN | None |
| PKP3 | RNF212 | None |
| PKP3 | SNX2 | None |
| PKP3 | MRPL33 | None |
| PKP3 | HADHA | None |
| CKMT1A | PHF14 | None |
| CKMT1A | ISPD | None |
| CKMT1B | PHF14 | None |
| CKMT1B | ISPD | None |
| GFOD2 | FAP | None |
| GFOD2 | RBM12 | None |
| RNGTT | PSMG4 | None |
| RNGTT | ARL17A | None |
| RNGTT | MAN1A2 | None |
| RNGTT | ARL17B | None |
| DDX39B | PSMG4 | None |
| DDX39B | ARL17A | None |
| DDX39B | MAN1A2 | None |
| DDX39B | ARL17B | None |
| FIBP | ACSBG1 | None |
| RAB31 | ACSBG1 | None |
| SIRT2 | GIGYF1 | None |
| SIRT2 | TPD52 | None |
| AK3 | GIGYF1 | None |
| AK3 | TPD52 | None |
| FAM81A | GIGYF1 | None |
| FAM81A | TPD52 | None |
| NELFE | GIGYF1 | None |
| NELFE | TPD52 | None |
| PHF14 | SKIL | None |
| SUGP2 | S100A14 | None |
| KHDRBS3 | S100A14 | None |
| PFAS | S100A14 | None |
| KIAA1211L | S100A14 | None |
| LARS | GIGYF1 | None |
| BZW2 | PRDX6 | None |
| BZW2 | RAF1 | None |
| BZW2 | NAGA | None |

**Supplementary Table 8.** Unique cancer type pairs of the significant MAPP of the TCPA dataset. TCPA dataset cancer symbols were given.

| Cancer type 1 | Cancer type 2 |
| --- | --- |
| LGG | CESC |
| LGG | ESCA |
| LGG | HNSC |
| LGG | THYM |
| LGG | BLCA |
| LGG | LUSC |
| PCPG | HNSC |
| PCPG | CESC |
| PCPG | ESCA |
| PCPG | THYM |
| PCPG | BLCA |
| PCPG | LUSC |
| GBM | BRCA |
| GBM | SARC |
| GBM | CESC |
| GBM | OV |
| LGG | BRCA |
| LGG | SARC |
| LGG | OV |
| KIRC | BRCA |
| KIRC | OV |
| KIRC | SARC |
| CESC | KIRC |
| SKCM | BLCA |
| LUSC | SKCM |
| PCPG | OV |
| GBM | HNSC |
| CESC | SKCM |
| CESC | UCEC |
| CESC | MESO |
| CESC | KICH |
| CESC | LUAD |
| CESC | LUSC |
| PRAD | SKCM |
| PRAD | UCEC |
| PRAD | MESO |
| PRAD | KICH |
| PRAD | KIRC |
| PRAD | LUAD |
| PRAD | LUSC |
| KIRP | UCEC |
| KIRP | MESO |
| KIRP | SKCM |
| KIRP | KICH |
| KIRP | KIRC |
| KIRP | LUAD |
| KIRP | LUSC |
| MESO | SKCM |
| MESO | KICH |
| MESO | KIRC |
| MESO | LUAD |
| MESO | LUSC |
| MESO | UCEC |
| BRCA | MESO |
| BRCA | SKCM |
| BRCA | KICH |
| BRCA | LUAD |
| BRCA | LUSC |
| BRCA | UCEC |
| OV | MESO |
| OV | SKCM |
| OV | KICH |
| OV | LUAD |
| OV | LUSC |
| OV | UCEC |
| THYM | SKCM |
| THYM | MESO |
| THYM | KICH |
| THYM | KIRC |
| THYM | LUAD |
| THYM | LUSC |
| THYM | UCEC |
| KIRC | KICH |
| KIRC | SKCM |
| KIRC | LUAD |
| KIRC | LUSC |
| KIRC | UCEC |
| CHOL | KICH |
| CHOL | SKCM |
| CHOL | KIRC |
| CHOL | LUAD |
| CHOL | LUSC |
| CHOL | MESO |
| CHOL | UCEC |
| THCA | KICH |
| THCA | SKCM |
| THCA | KIRC |
| THCA | LUAD |
| THCA | LUSC |
| THCA | MESO |
| THCA | UCEC |
| BLCA | KIRC |
| BLCA | LUAD |
| BLCA | LUSC |
| BLCA | MESO |
| BLCA | UCEC |
| SARC | MESO |
| SARC | SKCM |
| SARC | UCEC |
| SARC | LUSC |
| PAAD | SKCM |
| PAAD | UCEC |
| PAAD | LUSC |
| LIHC | SKCM |
| LIHC | UCEC |
| LIHC | LUSC |
| MESO | PAAD |
| MESO | LIHC |
| KICH | BLCA |
| KICH | SARC |
| KICH | PAAD |
| KICH | LIHC |
| KIRC | PAAD |
| KIRC | LIHC |
| LUAD | SARC |
| LUAD | PAAD |
| LUAD | LIHC |
| BRCA | PCPG |
| STAD | LGG |
| UCEC | LGG |
| PCPG | STAD |
| PCPG | UCEC |
| OV | SARC |
| OV | READ |
| OV | THCA |
| OV | BLCA |
| OV | UCS |
| KIRC | STAD |
| SARC | LIHC |
| SARC | STAD |
| MESO | STAD |
| KIRP | STAD |
| KIRP | LIHC |
| HNSC | SKCM |
| HNSC | LIHC |
| STAD | SKCM |
| STAD | LIHC |
| UCEC | SKCM |
| BRCA | SARC |
| BRCA | KIRP |
| BRCA | HNSC |
| BRCA | STAD |
| STAD | HNSC |
| STAD | UCEC |
| SARC | THYM |
| SARC | CESC |
| UCS | SARC |
| DLBC | SARC |
| THCA | SARC |
| COAD | SARC |
| TGCT | CESC |
| SARC | TGCT |
| ESCA | GBM |
| ESCA | THCA |
| ESCA | SARC |
| ESCA | COAD |
| ESCA | READ |
| SARC | READ |
| CESC | READ |
| THCA | CESC |
| COAD | CESC |
| SARC | KIRP |
| CESC | LIHC |
| CESC | CHOL |
| BLCA | CESC |
| GBM | BLCA |
| STAD | OV |
| STAD | BLCA |
| COAD | PCPG |
| COAD | SKCM |
| COAD | MESO |
| ESCA | SKCM |
| ESCA | MESO |
| READ | MESO |
| READ | PCPG |
| HNSC | SARC |
| HNSC | MESO |
| SKCM | READ |
| ESCA | TGCT |
| HNSC | KIRC |
| HNSC | LUAD |
| HNSC | KICH |
| CESC | HNSC |
| HNSC | OV |
| BRCA | TGCT |
| HNSC | TGCT |
| LUSC | TGCT |
| PRAD | BLCA |
| CHOL | BLCA |
| BRCA | BLCA |
| SARC | BLCA |
| THYM | BLCA |
| THCA | BLCA |
| PAAD | BLCA |
| TGCT | BLCA |
| LUAD | LGG |
| PAAD | KIRP |
| LGG | PAAD |
| KIRP | BLCA |
| KICH | LGG |
| PCPG | LGG |
| MESO | LGG |
| LGG | GBM |
| KICH | GBM |
| MESO | GBM |
| LUSC | GBM |
| LGG | LIHC |
| GBM | LIHC |
| GBM | KIRC |
| KIRP | LGG |
| STAD | LUSC |
| COAD | LGG |
| COAD | LUSC |
| HNSC | LUSC |
| UCEC | LUSC |
| LUAD | LUSC |
| KIRC | LGG |
| READ | LGG |
| READ | LUSC |
| KICH | LUSC |
| LGG | THCA |
| LGG | SKCM |
| PAAD | SARC |
| PAAD | OV |
| STAD | GBM |
| KICH | SKCM |
| PCPG | LUAD |
| UVM | BLCA |
| PCPG | SKCM |
| KICH | PCPG |
| OV | CESC |
| STAD | CESC |
| COAD | OV |
| COAD | BRCA |
| OV | BRCA |
| PRAD | OV |
| PAAD | CESC |
| PAAD | HNSC |
| UCEC | KICH |
| BRCA | READ |
| DLBC | LGG |
| DLBC | PRAD |
| TGCT | PRAD |
| TGCT | LGG |
| DLBC | GBM |
| KIRP | ACC |
| LUAD | STAD |
| LUAD | COAD |
| LUAD | SKCM |
| LUAD | GBM |
| READ | LIHC |
| READ | DLBC |
| READ | ACC |
| LGG | UCS |
| GBM | UCS |
| OV | LIHC |
| LUAD | READ |
| DLBC | BLCA |
| HNSC | DLBC |

**Supplementary Table 9.** Unique cancer type pairs of the significant MAPP of the HPA dataset.

| Cancer type 1 | Cancer type 2 |
| --- | --- |
| breast cancer | cervical cancer |
| breast cancer | colorectal cancer |
| breast cancer | endometrial cancer |
| breast cancer | glioma |
| breast cancer | melanoma |
| breast cancer | skin cancer |
| breast cancer | testis cancer |
| cervical cancer | melanoma |
| cervical cancer | prostate cancer |
| cervical cancer | testis cancer |
| colorectal cancer | melanoma |
| endometrial cancer | melanoma |
| endometrial cancer | stomach cancer |
| endometrial cancer | testis cancer |
| glioma | melanoma |
| glioma | prostate cancer |
| glioma | testis cancer |
| melanoma | pancreatic cancer |
| melanoma | prostate cancer |
| melanoma | stomach cancer |
| melanoma | urothelial cancer |
| ovarian cancer | testis cancer |
| pancreatic cancer | testis cancer |

**Supplementary Table 10.** MAPP and dataset ratios of the CT of the TCPA dataset were given. MAPP ratio was defined as the number of times a CT was used for MAPP, divided by the total MAPP number. Dataset ratio was defined as the number of counts of a CT in the dataset, divided by the total numbers of CT values of the dataset.

| Cancer type (CT) | MAPP ratio | Dataset ratio |
| --- | --- | --- |
| UVM | 0.001669 | 0.00156 |
| ACC | 0.003339 | 0.005979 |
| UCS | 0.006678 | 0.006239 |
| CHOL | 0.015025 | 0.003899 |
| TGCT | 0.015025 | 0.015337 |
| DLBC | 0.016694 | 0.004289 |
| THYM | 0.021703 | 0.011697 |
| PRAD | 0.025042 | 0.04562 |
| THCA | 0.026711 | 0.048349 |
| KIRP | 0.036728 | 0.027034 |
| PAAD | 0.036728 | 0.013647 |
| LIHC | 0.040067 | 0.023915 |
| ESCA | 0.041736 | 0.016376 |
| COAD | 0.043406 | 0.0464 |
| READ | 0.048414 | 0.016896 |
| KICH | 0.053422 | 0.008188 |
| UCEC | 0.063439 | 0.052508 |
| GBM | 0.068447 | 0.026644 |
| LUAD | 0.071786 | 0.04705 |
| STAD | 0.071786 | 0.050949 |
| PCPG | 0.073456 | 0.010398 |
| HNSC | 0.073456 | 0.04497 |
| BRCA | 0.080134 | 0.113595 |
| MESO | 0.09182 | 0.007928 |
| OV | 0.093489 | 0.053418 |
| LUSC | 0.095159 | 0.042241 |
| KIRC | 0.095159 | 0.057837 |
| BLCA | 0.105175 | 0.04471 |
| SKCM | 0.105175 | 0.04588 |
| CESC | 0.136895 | 0.022225 |
| SARC | 0.141903 | 0.028724 |
| LGG | 0.200334 | 0.055498 |

**Supplementary Table 11.** MAPP and dataset ratios of the CT of the HPA dataset were given. MAPP ratio was defined as the number of times a CT was used for MAPP, divided by the total MAPP number. Dataset ratio was defined as the number of counts of a CT in the dataset, divided by the total numbers of CT values of the dataset.

| Cancer type (CT) | MAPP ratio | Dataset ratio |
| --- | --- | --- |
| ovarian cancer | 0.013699 | 0.061095 |
| pancreatic cancer | 0.027397 | 0.052875 |
| urothelial cancer | 0.027397 | 0.059954 |
| colorectal cancer | 0.034247 | 0.060504 |
| prostate cancer | 0.047945 | 0.056166 |
| skin cancer | 0.061644 | 0.058097 |
| stomach cancer | 0.068493 | 0.054786 |
| endometrial cancer | 0.157534 | 0.060907 |
| cervical cancer | 0.232877 | 0.059632 |
| glioma | 0.232877 | 0.057824 |
| melanoma | 0.239726 | 0.059021 |
| testis cancer | 0.287671 | 0.058264 |
| breast cancer | 0.568493 | 0.059531 |

**Supplementary Table 12.** Symbol match between TCPA and NCBI.

| TCPA Symbol | NCBI Symbol 1 | NCBI Symbol 2 | NCBI Symbol 3 |
| --- | --- | --- | --- |
| X1433EPSILON | YWHAE | - | - |
| X4EBP1 | EIF4EBP1 | - | - |
| X4EBP1_pS65 | EIF4EBP1 | - | - |
| X4EBP1_pT37T46 | EIF4EBP1 | - | - |
| X53BP1 | TP53BP1 | - | - |
| ACC_pS79 | ACACA | ACACB | - |
| ACC1 | ACACA | - | - |
| AKT | AKT1 | AKT2 | AKT3 |
| AKT_pS473 | AKT1 | AKT2 | AKT3 |
| AKT_pT308 | AKT1 | AKT2 | AKT3 |
| AMPKALPHA | PRKAA1 | - | - |
| AMPKALPHA_pT172 | PRKAA1 | - | - |
| AR | AR | - | - |
| ASNS | ASNS | - | - |
| ATM | ATM | - | - |
| BAK | BAK1 | - | - |
| BAX | BAX | - | - |
| BCL2 | BCL2 | - | - |
| BCLXL | BCL2L1 | - | - |
| BECLIN | BECN1 | - | - |
| BETACATENIN | CTNNB1 | - | - |
| BID | BID | - | - |
| BIM | BCL2L11 | - | - |
| CJUN_pS73 | JUN | - | - |
| CKIT | KIT | - | - |
| CMET_pY1235 | MET | - | - |
| CMYC | MYC | - | - |
| CRAF | RAF1 | - | - |
| CRAF_pS338 | RAF1 | - | - |
| CASPASE7CLEAVEDD198 | CASP7 | - | - |
| CAVEOLIN1 | CAV1 | - | - |
| CD31 | PECAM1 | - | - |
| CD49B | ITGA2 | - | - |
| CDK1 | CDK1 | - | - |
| CHK1 | CHEK1 | - | - |
| CHK1_pS345 | CHEK1 | - | - |
| CHK2 | CHEK2 | - | - |
| CHK2_pT68 | CHEK2 | - | - |
| CIAP | BIRC2 | - | - |
| CLAUDIN7 | CLDN7 | - | - |
| COLLAGENVI | COL6A1 | - | - |
| CYCLINB1 | CCNB1 | - | - |
| CYCLIND1 | CCND1 | - | - |
| CYCLINE1 | CCNE1 | - | - |
| DJ1 | PARK7 | - | - |
| DVL3 | DVL3 | - | - |
| ECADHERIN | CDH1 | - | - |
| EEF2 | EEF2 | - | - |
| EEF2K | EEF2K | - | - |
| EGFR | EGFR | - | - |
| EGFR_pY1068 | EGFR | - | - |
| EGFR_pY1173 | EGFR | - | - |
| EIF4E | EIF4E | - | - |
| ERALPHA | ESR1 | - | - |
| ERALPHA_pS118 | ESR1 | - | - |
| ERK2 | MAPK1 | - | - |
| FIBRONECTIN | FN1 | - | - |
| FOXO3A | FOXO3 | - | - |
| GAB2 | GAB2 | - | - |
| GATA3 | GATA3 | - | - |
| GSK3ALPHABETA | GSK3A | GSK3B | - |
| GSK3ALPHABETA_pS21S9 | GSK3A | GSK3B | - |
| HER2 | ERBB2 | - | - |
| HER2_pY1248 | ERBB2 | - | - |
| HER3 | ERBB3 | - | - |
| HER3_pY1289 | ERBB3 | - | - |
| HSP70 | HSPA1A | - | - |
| IGFBP2 | IGFBP2 | - | - |
| INPP4B | INPP4B | - | - |
| IRS1 | IRS1 | - | - |
| JNK_pT183Y185 | MAPK8 | - | - |
| JNK2 | MAPK9 | - | - |
| KU80 | XRCC5 | - | - |
| LCK | LCK | - | - |
| LKB1 | STK11 | - | - |
| MAPK_pT202Y204 | MAPK1 | MAPK3 | - |
| MEK1 | MAP2K1 | - | - |
| MEK1_pS217S221 | MAP2K1 | - | - |
| MIG6 | ERRFI1 | - | - |
| MRE11 | MRE11 | - | - |
| MTOR | MTOR | - | - |
| MTOR_pS2448 | MTOR | - | - |
| NCADHERIN | CDH2 | - | - |
| NFKBP65_pS536 | NFKB1 | - | - |
| NF2 | NF2 | - | - |
| NOTCH1 | NOTCH1 | - | - |
| PCADHERIN | CDH3 | - | - |
| P27 | CDKN1B | - | - |
| P27_pT157 | CDKN1B | - | - |
| P38MAPK | MAPK14 | - | - |
| P38_pT180Y182 | MAPK14 | - | - |
| P53 | TP53 | - | - |
| P70S6K1 | RPS6KB1 | - | - |
| P70S6K_pT389 | RPS6KB1 | - | - |
| P90RSK_pT359S363 | RPS6KA1 | - | - |
| PAI1 | SERPINE1 | - | - |
| PAXILLIN | PXN | - | - |
| PCNA | PCNA | - | - |
| PDK1_pS241 | PDPK1 | - | - |
| PEA15 | PEA15 | - | - |
| PI3KP110ALPHA | PIK3CA | - | - |
| PKCALPHA | PRKCA | - | - |
| PKCALPHA_pS657 | PRKCA | - | - |
| PKCDELTA_pS664 | PRKCD | - | - |
| PR | PGR | - | - |
| PRAS40_pT246 | AKT1S1 | - | - |
| PTEN | PTEN | - | - |
| RAD50 | RAD50 | - | - |
| RAD51 | RAD51 | - | - |
| RB_pS807S811 | RB1 | - | - |
| S6 | RPS6 | - | - |
| S6_pS235S236 | RPS6 | - | - |
| S6_pS240S244 | RPS6 | - | - |
| SHC_pY317 | SHC1 | - | - |
| SMAD1 | SMAD1 | - | - |
| SMAD3 | SMAD3 | - | - |
| SMAD4 | SMAD4 | - | - |
| SRC | SRC | - | - |
| SRC_pY416 | SRC | - | - |
| SRC_pY527 | SRC | - | - |
| STAT3_pY705 | STAT3 | - | - |
| STAT5ALPHA | STAT5A | - | - |
| STATHMIN | STMN1 | - | - |
| SYK | SYK | - | - |
| TUBERIN | TSC2 | - | - |
| VEGFR2 | KDR | - | - |
| XRCC1 | XRCC1 | - | - |
| YAP | YAP1 | - | - |
| YAP_pS127 | YAP1 | - | - |
| YB1 | YBX1 | - | - |
| YB1_pS102 | YBX1 | - | - |
| X4EBP1_pT70 | EIF4EBP1 | - | - |
| ARAF_pS299 | ARAF | - | - |
| ANNEXINVII | ANXA7 | - | - |
| ARID1A | ARID1A | - | - |
| BRAF | BRAF | - | - |
| BAD_pS112 | BAD | - | - |
| BAP1C4 | BAP1 | - | - |
| BRCA2 | BRCA2 | - | - |
| CD20 | MS4A1 | - | - |
| CYCLINE2 | CCNE2 | - | - |
| ETS1 | ETS1 | - | - |
| EIF4G | EIF4G1 | - | - |
| FASN | FASN | - | - |
| FOXO3A_pS318S321 | FOXO3 | - | - |
| FOXM1 | FOXM1 | - | - |
| G6PD | G6PD | - | - |
| GAPDH | GAPDH | - | - |
| GSK3_pS9 | GSK3A | GSK3B | - |
| HEREGULIN | NRG1 | - | - |
| MYH11 | MYH11 | - | - |
| MYOSINIIA_pS1943 | MYH9 | - | - |
| NRAS | NRAS | - | - |
| NDRG1_pT346 | NDRG1 | - | - |
| P21 | CDKN1A | - | - |
| P27_pT198 | CDKN1B | - | - |
| P90RSK | RPS6KA1 | - | - |
| PDCD4 | PDCD4 | - | - |
| PDK1 | PDPK1 | - | - |
| PEA15_pS116 | PEA15 | - | - |
| PI3KP85 | PIK3R1 | - | - |
| PKCPANBETAII_pS660 | PRRT2 | - | - |
| PRDX1 | PRDX1 | - | - |
| RAB11 | RAB11A | RAB11B | - |
| RAB25 | RAB25 | - | - |
| RAPTOR | RPTOR | - | - |
| RBM15 | RBM15 | - | - |
| RICTOR | RICTOR | - | - |
| RICTOR_pT1135 | RICTOR | - | - |
| SCD1 | SCD | - | - |
| SF2 | SRSF1 | - | - |
| TAZ | WWTR1 | - | - |
| TIGAR | TIGAR | - | - |
| TRANSGLUTAMINASE | TGM2 | - | - |
| TFRC | TFRC | - | - |
| TSC1 | TSC1 | - | - |
| TUBERIN_pT1462 | TSC2 | - | - |
| EPPK1 | EPPK1 | - | - |
| XBP1 | XBP1 | - | - |
| ACETYLATUBULINLYS40 | TUBA1B | - | - |
| P62LCKLIGAND | SQSTM1 | - | - |
| X1433BETA | YWHAB | - | - |
| X1433ZETA | YWHAZ | - | - |
| ACVRL1 | ACVRL1 | - | - |
| DIRAS3 | DIRAS3 | - | - |
| ANNEXIN1 | ANXA1 | - | - |
| PREX1 | PREX1 | - | - |
| ENY2 | ENY2 | - | - |
| GCN5L2 | KAT2A | - | - |
| ADAR1 | ADAR | - | - |
| JAB1 | COPS5 | - | - |
| CMET | MET | - | - |
| CASPASE8 | CASP8 | - | - |
| ERCC1 | ERCC1 | - | - |
| MSH2 | MSH2 | - | - |
| MSH6 | MSH6 | - | - |
| PARPCLEAVED | PARP1 | - | - |
| RB | RB1 | - | - |
| SETD2 | SETD2 | - | - |
| SMAC | DIABLO | - | - |
| SNAIL | SNAI1 | - | - |
| AXL | AXL | - | - |
| MYOSINIIA | MYH9 | - | - |
| SLC1A5 | SLC1A5 | - | - |
| GATA6 | GATA6 | - | - |
| BRD4 | BRD4 | - | - |
| CDK1_pY15 | CDK1 | - | - |
| ARAF | ARAF | - | - |
| BRAF_pS445 | BRAF | - | - |
| BCL2A1 | BCL2A1 | - | - |
| CABL | ABL1 | - | - |
| CASPASE3 | CASP3 | - | - |
| CD26 | DPP4 | - | - |
| CHK1_pS296 | CHEK1 | - | - |
| COG3 | COG3 | - | - |
| DUSP4 | DUSP4 | - | - |
| ERCC5 | ERCC5 | - | - |
| IGF1R_pY1135Y1136 | IGF1R | INSR | - |
| IRF1 | IRF1 | - | - |
| JAK2 | JAK2 | - | - |
| P16INK4A | CDKN2A | - | - |
| SHP2_pY542 | PTPN11 | - | - |
| PDL1 | CD274 | - | - |
| PARP1 | PARP1 | - | - |
| CA9 | CA9 | - | - |
| COMPLEXIISUBUNIT30 | SDHB | - | - |
| GYGGLYCOGENIN1 | GYG1 | - | - |
| GYS | GYS1 | - | - |
| GYS_pS641 | GYS1 | - | - |
| HIF1ALPHA | HIF1A | - | - |
| LDHA | LDHA | - | - |
| LDHB | LDHB | - | - |
| MITOCHONDRIA | - | - | - |
| OXPHOSCOMPLEXVSUBUNITB | ATP5F1B | - | - |
| PKM2 | PKM | - | - |
| PYGB | PYGB | - | - |
| PYGBAB2 | PYGB | - | - |
| PYGL | PYGL | - | - |
| PYGM | PYGM | - | - |
| CTLA4 | CTLA4 | - | - |
| PDCD1 | PDCD1 | - | - |
| CASPASE9 | CASP9 | - | - |
| E2F1 | E2F1 | - | - |
| EZH2 | EZH2 | - | - |
| KEAP1 | KEAP1 | - | - |
| LCN2A | LCN2 | - | - |
| MACC1 | MACC1 | - | - |
| NRF2 | NFE2L2 | - | - |
| PARPAB3 | PARP1 | - | - |
| THYMIDILATESYNTHASE | TYMS | - | - |
| TTF1 | TTF1 | - | - |
| CHROMOGRANINANTERM | CHGA | - | - |
| CK5 | KRT5 | - | - |
| NAPSINA | NAPSA | - | - |
| P63 | TP63 | - | - |
| RET_pY905 | RET | - | - |
| SYNAPTOPHYSIN | SYP | - | - |
| ALPHACATENIN | CTNNA1 | CTNNA3 | - |

**Supplementary Table 13.** Normal tissue names for the cancer types of the HPA dataset.

| Cancer type | Normal tissue 1 | Normal tissue 2 | Normal tissue 3 | Normal tissue 4 | Normal tissue 5 | Normal tissue 6 | Normal tissue 7 |
| --- | --- | --- | --- | --- | --- | --- | --- |
| breast cancer | breast | - | - | - | - | - | - |
| cervical cancer | cervix, uterine | - | - | - | - | - | - |
| colorectal cancer | colon | rectum | - | - | - | - | - |
| endometrial cancer | endometrium 1 | endometrium 2 | - | - | - | - | - |
| glioma | caudate | cerebral cortex | hippocampus | hypothalamus | dorsal raphe | cerebellum | substantia nigra |
| liver cancer | liver | - | - | - | - | - | - |
| lung cancer | lung | - | - | - | - | - | - |
| lymphoma | lymph node | - | - | - | - | - | - |
| melanoma | skin 1 | - | - | - | - | - | - |
| ovarian cancer | ovary | - | - | - | - | - | - |
| pancreatic cancer | pancreas | - | - | - | - | - | - |
| prostate cancer | prostate | - | - | - | - | - | - |
| renal cancer | kidney | - | - | - | - | - | - |
| skin cancer | skin 1 | - | - | - | - | - | - |
| stomach cancer | stomach 1 | stomach 2 | - | - | - | - | - |
| testis cancer | testis | - | - | - | - | - | - |
| urothelial cancer | urinary bladder | - | - | - | - | - | - |

**Supplementary Table 14.** Normal cell type names for the cancer types of the HPA dataset.

| Cancer type | Normal cell type 1 | Normal cell type 2 |
| --- | --- | --- |
| breast cancer | glandular cells | myoepithelial cells |
| cervical cancer | glandular cells | squamous epithelial cells |
| colorectal cancer | glandular cells | - |
| endometrial cancer | glandular cells | - |
| glioma | glial cells | - |
| liver cancer | hepatocytes | - |
| lung cancer | pneumocytes | - |
| lymphoma | germinal center cells | non-germinal center cells |
| melanoma | melanocytes | - |
| ovarian cancer | follicle cells | - |
| pancreatic cancer | exocrine glandular cells | - |
| prostate cancer | glandular cells | - |
| renal cancer | cells in tubules | - |
| skin cancer | keratinocytes | - |
| stomach cancer | glandular cells | - |
| testis cancer | spermatogonia cells | - |
| urothelial cancer | urothelial cells | - |
